# Altered interaction between RBD and ACE2 receptor contributes towards the increased transmissibility of SARS CoV-2 delta, kappa, beta, and gamma strains with RBD double mutations

**DOI:** 10.1101/2021.08.30.458303

**Authors:** Siddharth Sinha, Benjamin Tam, San Ming Wang

**Affiliations:** Frontiers Science Center for Precision Oncology, Faculty of Health Sciences, University of Macau, Macau

**Keywords:** SARS_CoV-2, Variants, MD simulations, Free energy, Antibody escape.

## Abstract

The COVID-19 pandemics by SARS-CoV-2 causes catastrophic damage for global human health. The initial step of SARS-CoV-2 infection is the binding of the receptor-binding domain (RBD) in its spike protein to ACE2 receptor in host cell membrane. The evolving of SARS-CoV-2 constantly generates new mutations across its genome including RBD. Besides the well-known single mutation in RBD, the recent new mutation strains with RBD “double mutation” is causing new outbreaks globally, as represented by the delta strain containing RBD L452R/T478K. Although it is considered that the increased transmissibility of the double mutated strains could be attributed to the alteration of mutated RBD to ACE2 receptor, the molecular details remains to be unclear. Using the methods of molecular dynamics simulation, superimposed structural comparison, free binding energy estimation and antibody escaping, we investigated the relationship between ACE2 receptor and the RBD double mutant L452R/T478K (delta), L452R/E484Q (kappa) and E484K/N501Y (beta, gamma). The results demonstrated that each of the three RBD double mutants altered RBD structure, led to enhanced binding affinity of mutated RBD to ACE2 receptor, leading to increased transmissibility of SARS-CoV-2 to the host cells.

## INTRODUCTION

SARS-CoV-2 pandemic has caused devastating consequences on global public health, with over 207 million people infected and over 4.3 million loss of life globally since the COVID-19 started (https://covid19.who.int accessed August 17, 2021). SARS-CoV-2 infects human cells through its spike (S) protein. In the process, the receptor-binding domain (RBD, residues 318–526) of the S protein (residues 1-1273)binds to the angiotensin-converting enzyme 2 (ACE2) receptor on host cell membrane to release its genome into the cell (Lan et al., 2020; Q. Wang et al., 2020). Therefore, RBD is a determinant for SARS-CoV-2 infection into host cells. As RNA virus, the genome of SARS-CoV-2 is constantly evolving with new mutations generated across its genome including RBD. Since the first SARS-CoV-2 genome sequences reported in Jan 5, 2020, there have been 942 coding-changing mutations identified within the 193 positions of RBD, more than one mutation per day (942 mutations in 932 days), and 4.9 mutations per position on average (Supplementary table 1, http://cov-glue.cvr.gla.ac.uk/#/replacement, access July 31, 2021). While most of the mutations do not show pathogenic significance, the strains containing several RBD mutations, namely L452R (epsilon), T478K, E484K, E484Q, and N501Y, were selected (Figure 1) and have caused multiple outbreaks much due to the increased transmissibility of SARS-CoV-2 contributed by these RBD mutations (Leung et al., 2021), https://www.who.int/en/activities/tracking-SARS-CoV-2-variants/). Recently, several new SARS-CoV-2 strains with RBD “double mutations” are causing new challenge. The double mutations basically contain the same single mutations above, but they are more transmissible than the strains with corresponding single mutation. For example, the “delta” strain with RBD mutation L452R/T478K rapidly spreads to over 130 countries since it was identified in late 2020 (Torjesen, 2021), https://www.cdc.gov/mmwr/volumes/70/wr/mm7031e2). L452R/E484Q (kalpa) and E484K/N501Y (beta, gamma). Thus, understanding the mechanism of increased transmissibility in RBD double mutations is urgently needed to develop strategies to control their spreading. While studies revealed how RBD single mutations increased SARS-CoV-2 transmissibility, it remains to know whether the SARS-CoV-2 with RBD double mutations adopted the same or similar manners as the SARS-CoV-2 with RBD single mutations or gains new features considering the more aggressive behavior of SARS-CoV-2 with RBD double mutations than SARS-CoV-2 with single RBD mutations.

**Figure 1.**
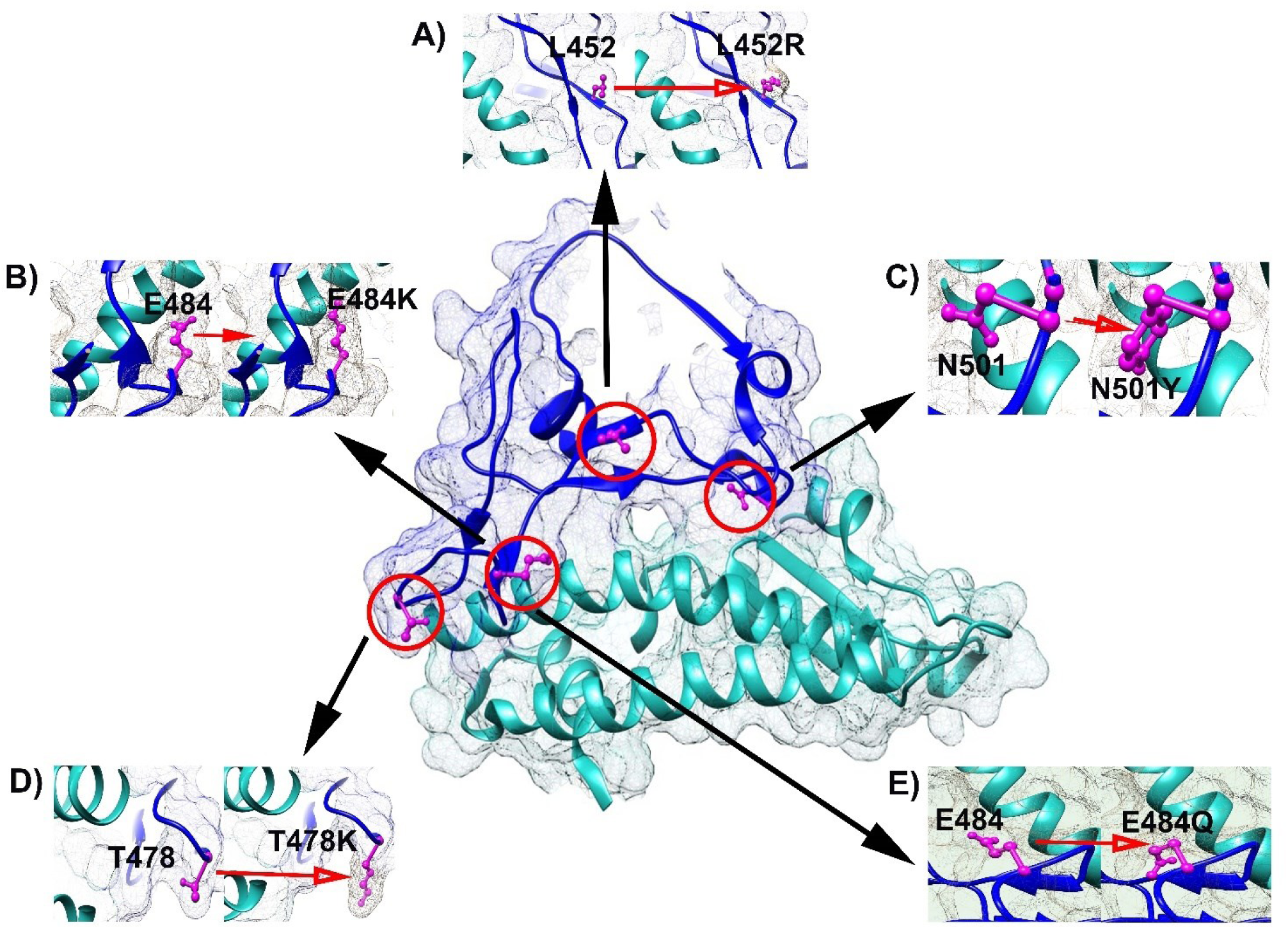
Locations of single mutated RBD and ACE2. The center shows the RBD-ACE2 complex, the surrounding shows the single mutations in RBD, with the left being the wild type and the right being the mutated residue. A) L452 and L452R; B) E484 and E484K; C) N501 and N501Y; D) T478 and T478K; E) E484 and E484Q.

In this study, we investigated the relationship between ACE2 receptor and three major RBD double mutant L452R/T478K (delta), L452R/E484Q (kappa) and E484K/N501Y (beta, gamma). We used multiple methods to address the topic including molecular dynamics simulation (MDS), superimposed structural comparison, free binding energy estimation and antibody escaping. Molecular dynamics simulation measures conformational change of protein across a time period and uses the trajectories to describe the thermodynamics changes in RBD mutant structure (Karplus, 2002; Sinha & Wang, 2020), superimposed structural comparison allows direct visualization of the altered RBD structure by the double mutated residues (Kufareva & Abagyan, 2012), free binding energy change estimates the affinity change caused by the double mutated residues between RBD and ACE2 receptor (Gumbart et al., 2013), and mapping antibody binding site explains whether RBD structural change caused by mutated residues can result in escape of mutants from neutralizing antibodies (NAbs). The results from our study provided evidence that the three RBD double mutants altered RBD structure in the ways much different from these caused by the corresponding RBD single mutants, enhanced binding of mutated RBD to ACE2 receptor, leading to the high transmissibility of SARS-CoV-2 to the host cells.

## RESULTS

### Conformation changes in RBD double mutants

To investigate the effects of double mutations on RBD structure, we examined conformational changes of the RBD double mutant - ACE2 receptor complex in RBD double mutant L452R/T478K, L452R/E484Q, E484K/N501Y using MD simulations for 100 ns. We performed the study by using 5 different types of methods including simulations of RMSD (Root mean square deviation), RMSF (Root mean square fluctuation), Rg (radius of gyration), SASA (solvent accessible surface area) and MSD (mean square displacement) for each type of RBD double mutant, with the wild type RBD and 5 RBD single mutants as the controls. The results revealed substantial structural differences between the double mutants, wild type, and single mutants:

#### RMSD

It demonstrates the change in the atomic coordinates between the wildtype and mutant structures (Dong et al., 2018). The wild type RBD-ACE2 structure stabilized around ∼0.3-0.4 nm with a deviation around 45 ns. L452R/T478K showed deviation upto ∼30 ns and thereafter stabilized at ∼0.3-0.4 nm; L452R/E484Q stabilized ∼0.3-0.5 nm with a deviation at around 35 ns; E484K/N501Y reached equilibrium distance at ∼0.23 nm. Single mutant L452R, T478K, E484Q and E484K all but N501Y showed stable configuration post 60 ns: L452R stabilized ∼0.25 nm, T478K ∼0.3 nm, E484Q ∼0.2 – 0.3 nm, E484K ∼0.3 nm. N501Y had 0.5 nm between 80 −100 ns. The results showed the trajectories in each of the RBD double mutants differed from the wild type and the corresponding single mutants (Figure 2).

**Figure 2.**
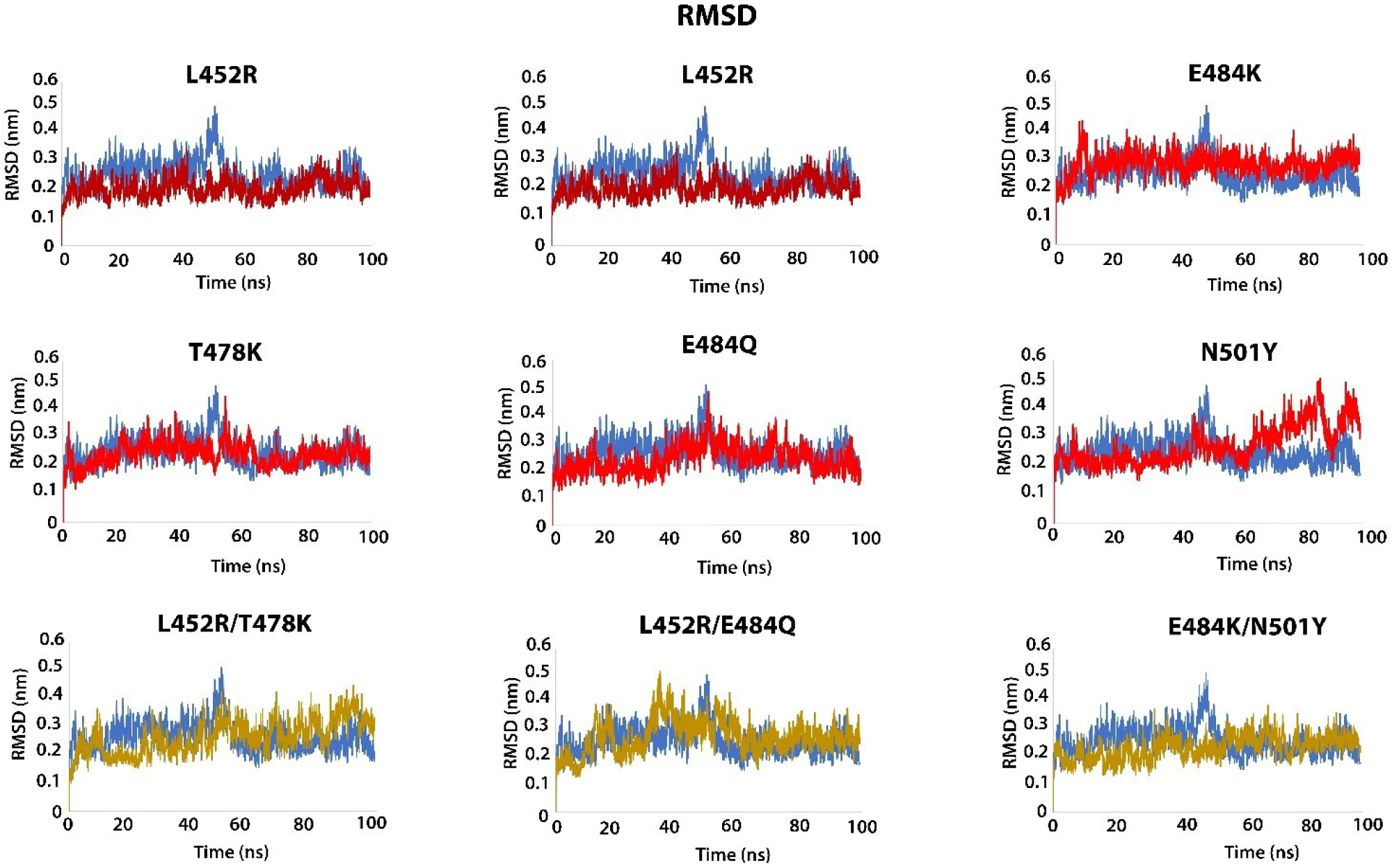
Dynamic changes of RBD structure by RMSD analysis. Blue: Wildtype, Red: single mutations (L452R, T478K, E484K, E484Y, N501Y); Yellow: double mutations (L452R/T478K, L452R/E484Q, E484K/N501Y). x-axis shows the simulation lasted time (100 ns), y-axis shows the RMSD value for the structures. The results show substantial differences between single and double mutated RBD structure.

#### RMSF

It determines the resilience of residues to analyze the effects of substitution in native and mutant structures (Benson & Daggett, 2012). The flexibility in polypeptide chain through RMSF showed that double mutant L452R/E484Q, L452R/T478K, and E484K/N501Y had high to medium resilience ∼ 0.4 nm, ∼0.15 nm, and ∼0.25 nm, respectively, at residue position 360-380 and 470-490. In comparison, the single mutants had no greater flexibility in the backbone C-α atom than the native structure (Supplementary Figure 1). The higher resilience in the double mutant L452R/E484Q and E484K/N501Y can be attributed to their differences in RMSD trajectory around residue position 360- 380 and 470-490.

#### Rg

It measures the compactness between the wild type and mutant structures (Daidone et al., 2003). Rg value for the wild type RBD was ∼3.10-3.15 nm. Double mutant L452R/T478K showed a greater Rg value ∼3.2-3.25 nm for the entire trajectory than the wild type; L452R/E484Q had ∼3.13-3.18 nm; E484K/N501Y had ∼3.15-3.25 nm after 35 ns; L452R/E484Q and E484K/N501Y showed the change in Rg value ∼80-100 ns and ∼10-30 ns respectively. Their larger hydrodynamic radius implies the change in the shape of structure during protein folding and unfolding. T478K, E484K, and E484Q had higher Rg values than the wild type RBD but lower than the double mutants, except N501Y had higher value ∼3.2-3.3 nm (Supplementary Figure 2).

#### SASA

SASA defines the solvent accessible surface area thereby measuring the relative expansion of the native and mutant structures (Zhang & Lazim, 2017). The wild type RBD had a surface area of ∼430 nm^2^, all single mutant L452R, T478K, E484K, E484Q and N501Y had decreased value of ∼426 nm^2^, ∼426 nm^2^, ∼424 nm^2^, ∼428 nm^2^ and ∼429 nm^2^, respectively. The double mutants also decreased values to these in single mutants but their patterns were different (Supplementary Figure 3).

#### MSD

MSD defines the mean square displacement of overall atoms from a set of initial positions between wild type RBD and RBD mutants (Yousefpour et al., 2015). The wild type structure showed an average displacement value ∼11.31. The double mutant L452R/T478K, L452R/E484Q and E484K/N501Y showed average displacement value ∼10.88, ∼7.33 and ∼9.06, respectively, lower than the wild type but higher than these in the single mutants (Supplementary Figure 4).

### Conformational changes in RBD double mutants

To directly visualize the conformational changes in double mutated RBD, we superimposed structure between wildtype, double mutants and the corresponding single mutants. We observed that the conformational changes in each of the double mutant RBD were substantially different from these in their corresponding single mutants (Figure 3). L452R/T478K showed conformational change at residue position 475-482 and 518-521 (Figure 3G), whereas both L452R/E484Q and E484K/N501Y were at residue position 475-485 (Figure 3H, 3I). L452 at the middle and N501 at the end of RBD-ACE2 interface of the β strand attributes to the change in conformations of related mutants. L452R induced conformational change at residue position 474-485 and 517-526 (Figure 3B), whereas N501Y at residue position 439-453 and 498-502 (Figure 3F). The conformational changes in double mutants comprising of L452R and N501Y were very different from their corresponding single mutants.

**Figure 3.**
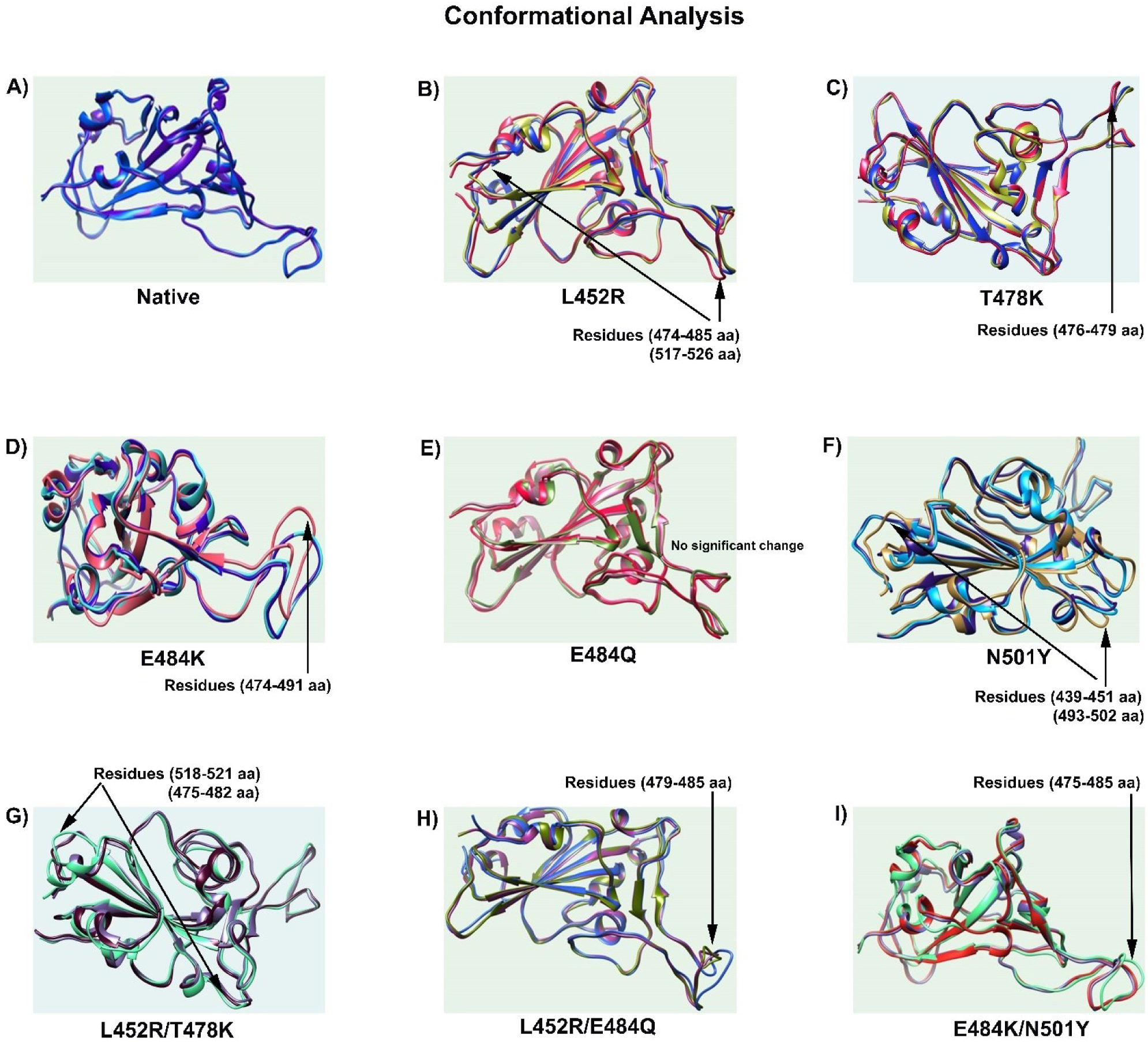
Conformational changes in mutated RBD. Mutant structures were superimposed with the wildtype RBD. Arrow shows conformation changes at specific residue positions. A) Native; B) L452R; C) T478K; D) E484K; E) E484Q; F) N501Y; G) L452R/T478K; H) L452R/E484Q; I) E484K/N501Y. The results show that the RBD conformational changes in double mutations were substantially different from single mutations.

### Free energy changes in RBD double mutants

We used MM/GBSA method to estimate overall changes of free binding energy between RBD mutants and ACE2 receptor. The high binding energy of RBD to ACE2 receptor is largely contributed by several key residues including N501, L452 and T478, which had 4, 2 and 2 contact regions, respectively, whereas other residues such as E484 contributed less binding energy than N501 to ACE2 receptor (Figure 4). Comparing to the overall free binding energy of 212.5 kJ mol^-1^ between the wild type RBD and ACE2 receptor, all single and double RBD mutants showed increased free binding energy except N501Y decreased to −204.6 kJ mol^-1^ (Table 1A). However, the changed levels between double mutants and their corresponding single mutants were at similar levels, implying double mutants didn’t generate higher binding energy than the single mutants.

**Figure 4.**
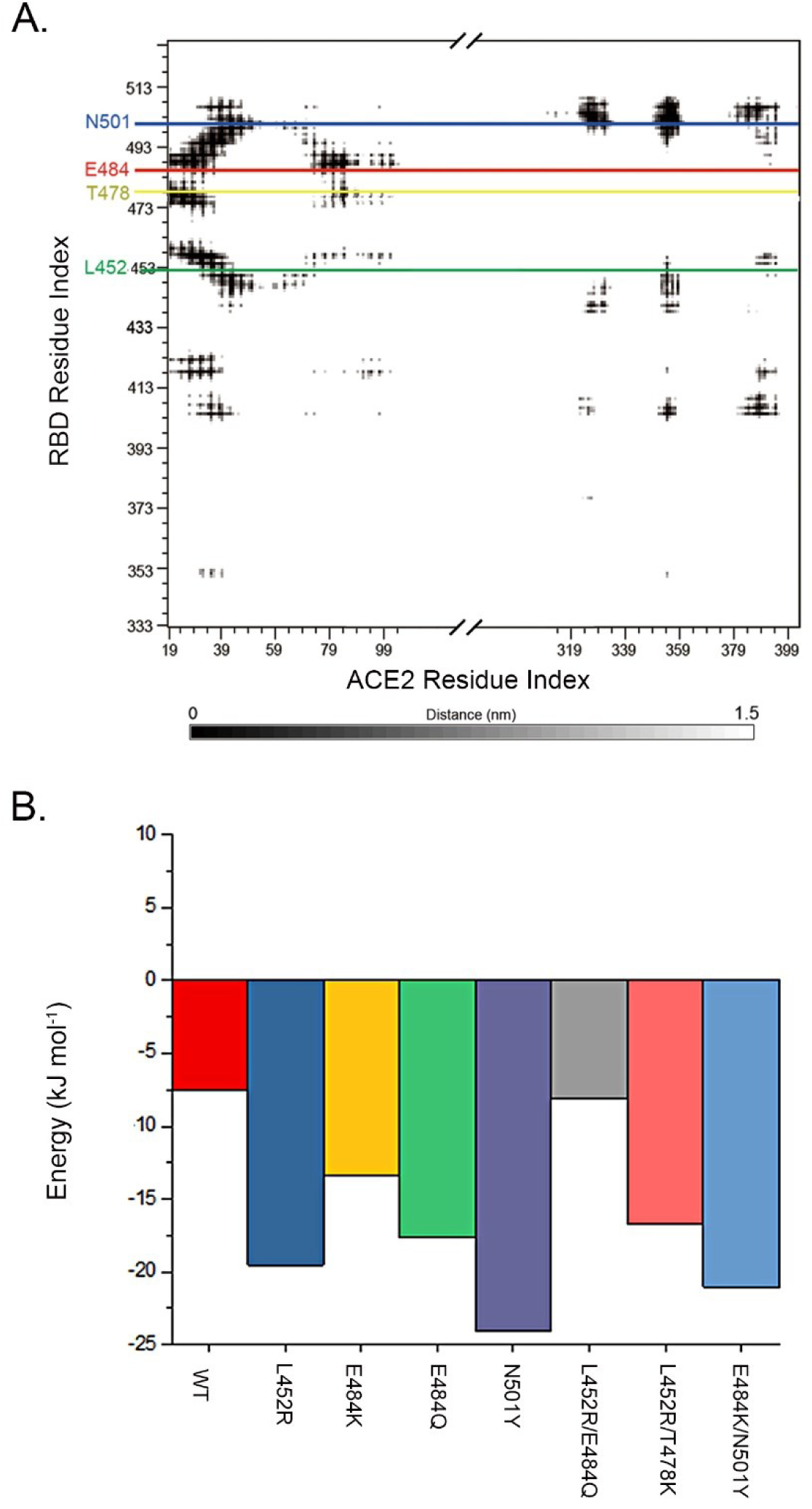
Free binding energy changes between mutated RBD and ACE2 receptor. A. RBD and ACE2 residue interaction map. Residues in close contact and the contact distance further than 1.5 nm are represented by diminishing grey scale. The key mutation positions (L452, T478, E484, N501) in RBD are highlighted in green, yellow, red and blue; B) binding energy by MM/GBSA for single RBD residues at position 501. It shows that while N501Y caused the highest increase of the binding energy at position 501, all single mutation (L452R, E484K, E484Q) and double mutation (L452R/E484Q, L452R/T478K, E484K/N501Y) also increased the binding energy at position 501. The red, blue, gold, green, purple, gray, pink and light blue represented WT, L452R, E484K, E484Q, N501Y, L452R/E484Q, L452R/T478K and E484K/N501Y, respectively.

**Table 1.** Free binding energy changes between mutated RBD and ACE2

| Mutant | Residue | Free binding energy (kJ mol<sup>-1</sup>) |
| --- | --- | --- |
| A, Overall free binding energy changes |  |  |
| Wild type |  | -212.5 |
| L452R |  | -288.1 |
| T478K |  | -229.6 |
| E484Q |  | -250.2 |
| E484K |  | -243.0 |
| N501Y |  | -204.6 |
| L452R/T478K |  | -256.1 |
| L452R/E484Q |  | -252.6 |
| E484K/N501Y |  | -264.5 |
| B. Top 5 non-mutated residues with free binding energy changes |  |  |
| Wild type | Q493 | -19.50 |
|  | Y505 | -19.12 |
|  | F486 | -14.85 |
|  | T500 | -12.55 |
|  | F456 | -11.41 |
| L452R | Q498 | -23.58 |
|  | Q493 | -19.57 |
|  | N501 | -19.54 |
|  | Y505 | -18.89 |
|  | F486 | -16.23 |
| T478K | Y505 | -19.15 |
|  | F486 | -15.92 |
|  | Q493 | -14.16 |
|  | T500 | -13.62 |
|  | F456 | -11.63 |
| E484K | Y505 | -21.61 |
|  | F486 | -15.50 |
|  | T500 | -13.73 |
|  | N501 | -13.36 |
|  | Y489 | -12.77 |
| E484Q | Q493 | -18.24 |
|  | N501 | -17.64 |
|  | Y505 | -17.16 |
|  | F486 | -15.14 |
|  | F456 | -12.33 |
| N501Y | Y501 | -24.05 |
|  | F486 | -15.66 |
|  | Y505 | -14.54 |
|  | F456 | -12.23 |
|  | Q493 | -11.86 |
| L452R/T478K | Y505 | -19.47 |
|  | F486 | -17.49 |
|  | N501 | -16.69 |
|  | Q493 | -13.63 |
|  | T500 | -12.99 |
| L452R/E484Q | F486 | -16.43 |
|  | Q493 | -15.38 |
|  | Y505 | -14.36 |
|  | Y489 | -13.17 |
|  | F456 | -12.56 |
| E484K/N501Y | Y501 | -21.05 |
|  | Y505 | -16.95 |
|  | Q493 | -16.35 |
|  | T500 | -15.79 |
|  | F486 | -14.90 |

We also compared the energy changes in the non-mutated residues in the RBD mutants. In the wild type RBD, a set of residues made high contributions to the overall energy between RBD and ACE2 receptor with the top 5 residues of F456, F486, Q493, T500, and Y505 (Table 1B). However, the top 5 residues in each RBD double mutant were mostly changed from the wild type RBD and their corresponding single mutations: L452/T478K changed to Y505, F486, N501, Q493, and T500; L452R/E484Q to F486, Q493, Y505, Y489 and F456; and E484K/N501Y to Y501, Y505, Q493, T500 and F486. N501 in the wildtype RBD contributed only −7.58 kJ mol^-1^ (9^th^) in the 198 RBD residues (data not shown). However, N501 became a top residue in 2 of the 3 RBD double mutants and 4 of the 5 RBD single mutants, highlighting that N501Y in RBD double mutants and single RBD mutants played significant roles in enhancing the affinity between RBD mutants to ACE2 receptor by enhancing the binding energy (Table 1B, Figure 4).

### Antibody escaping in RBD double mutants

SARS-CoV-2 neutralizing antibody (NAb) C121 (PDB ID: 7K8Z) is a well-known neutralizing antibody to SARS-CoV-2 through binging to RBD and E484K is a well-known mutant resistant to C121 binding (Barnes et al., 2020; Weisblum et al., 2020). We used C121 antibody and E484K as the model to test the relationship between RBD double mutants and antibody resistance. Figure 5 showed that the binding sites of C121 on S protein covering N439, N440, L455, G446, E484 and Q493 shown in yellow and E484K shown in green. Comparison of the binding sites between wild type RBD and double mutant RBD demonstrated that all three RBD double mutants altered C121 binding site, with L452R/E484Q as the most significant one among the three RBD double mutants. The results highlight L452R/E484K not only increased transmission but also facilitated antibody escape ability on SARS-CoV-2.

**Figure 5.**
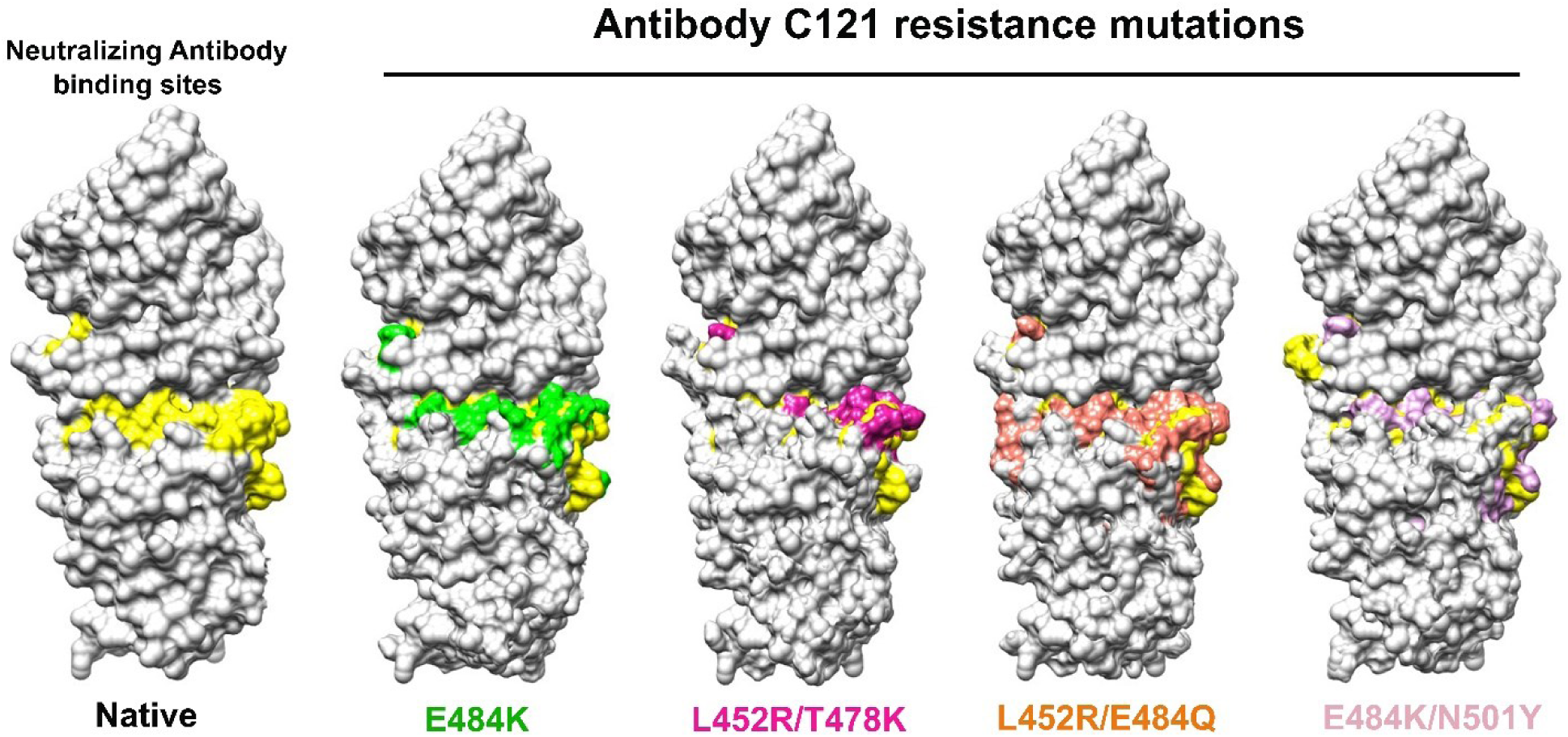
Changes of antibody binding sites in double mutated RBD. The binding site (N439, N440, L455, G446, E484 and Q493) of neutralizing antibody C121 (PDB ID: 7K8X) are compared between the wild type, E484K known escaping C121 binding, and double mutants L452R/T478K, L452R/E484Q and E484K/N501Y. Yellow: C121 binding sites on wild type RBD; Green: C121 binding sites on E484K RBD; Magenta: C121 binding sites on L452R/T478K RBD; Orange: C121 binding sites on L452R/E484Q RBD; Pink: C121 binding sites on E484K/N501Y RBD. The figure shows that all three double mutated RBD altered the C121 biding sites, with L452R/T478K (delta) as the most significant one.

## DISCUSSION

It has been proposed that higher transmissibility of SARS-CoV-2 new mutant strains can be attributed by the increased affinity of the mutated RBD to ACE2 receptor in the host cells, and the RBD double mutations can further increase the affinity and enhance the transmissibility, as represented by SARS-CoV-2 L452R/T478K (delta), L452R/E484Q (kappa), E484K/N501Y (beta and gamma) strains (Scudellari, 2021). Using combinational methods, our study provides structural-based evidence to demonstrate that RBD double mutant-changed RBD structure enhances the affinity of mutated RBD to ACE2 receptor, causing the increased transmissibility of the new SARS-CoV-2 mutant strains.

It is interesting to note that many of the “hot spot” mutants predicted by previous simulation studies have rarely become highly transmissible strains (Gan et al., 2021; Ghorbani et al., 2020; Y. Wang et al., 2020; Yi et al., 2020). Instead, only limited numbers of mostly unpredicted single mutations were truly selected to cause outbreaks. More interesting fact is that the RBD double mutations in the newly emerging strains still inherited these mutations at the same positions with different combinations. This indicates that although MDS is powerful in detecting structural changes caused by mutation, solely based on physical relationship between RBD and ACE2 receptor in the randomly detected mutation pool is not a reliable approach to determine the “hotspot” mutation as the marks for new strain with increased transmissibility. No matter how strong the physical interaction between mutated RBD to ACE2 receptor, only these with best fitness within the mutation pool will be selected, which is determined by multiple selection factors not only by the relationship between RBD and ACE2 receptor. In addition, relying on the accumulated quantity of mutation sequences may also give misleading information for the increased transmissibility, as the number of sequences collected could be biased sampling, not necessarily reflect the actual abundance of the mutant strains in actual infected population. This is reflected by the RBD mutation sequence data that only 4 (N501Y, L452R, E484K, Y478K) were listed among the 7 top mutation sequences, and E484Q was listed at 26^th^ position (Supplementary table 1). Therefore, focusing on the “real world” mutations either single, double and possible triple combinations at these fixed positions can provide more reliable evidence to monitor the newly selected strains and to use the physical evidence to explain the structural basis for these highly selected ones with high transmissibility (Khan et al., 2021).

N501 is particularly interesting residue. It has been determined that N501 enhances RBD binding to ACE2 by maintaining RBD at open conformation (Teruel *et al*., 2021), and this function can be enhanced by E484K (Nelson *et al*, 2021). N501 is also a mutation hot-spot as reflected by the RBD single mutation N501Y in the alpha strain (Fiorentini et al., 2021), and RBD double mutation E484K/N501Y in beta and gamma strains. We observed that each tested RBD single or double mutation increased the free binding energy between N501 and ACE2 receptor (Figure 4B), highlighting that N501 is a key player in enhancing wile type and mutated RBD binding to ACE2, not only for N501Y but also the mutated residues in other RBD locations likely through allosteric activity (Mugnai et al., 2020). For example, E484K and E484Q substantially increased binding affinity of N501 and N501Y to ACE2 receptor (Luan et al., 2021; Nelson et al., 2021). Similar situation was also present in RBD double mutants, and mutations outside RBD can also contribute to the increased affinity of RBD to ACE2 receptor, such as D614G (Korber et al., 2020).Therefore, not only the mutated RBD residues but also non-mutated RBD residues needs to be considered when addressing the increased transmissibility of new mutant strains.

Antibody resistance is one of the crucial determinants of viral transmission. Many potentially neutralizing anti SARS-CoV-2 antibodies target RBD to block its binding to the ACE-2 receptor (Baum et al., 2020; Ju et al., 2020; Starr et al., 2021). Consistent with experimental observations (Rogers et al., 2020), our study showed that all three RBD double mutants can cause antibody escape. Breakthrough infection by delta strain has been observed in the vaccinated individuals (Bergwerk et al., 2021). Our observation indicates that the altered antibody binding sites on RBD by RBD double mutation contributes to the phenomenon.

It is interesting to note that the binding energy between single mutants (L452R, T478K, E484K, E484Q and N501Y) and double mutants (L452/E484Q, L452/T478K and E484K/N501Y) were at similar level, although both were higher than the wild type RBD.

In conclusion, our study revealed that double mutated RBD in SARS-CoV-2 caused the changes of RBD structure, RBD binding energy to ACE2 receptor, and antibody binding sites on RBD. These changes contribute to the increased transmissibility of recent SARS-CoV-2 mutant strains with double mutated RBD.

## MATERIALS & METHODS

### Structures of RBD and mutated RBDs

The structure of SARS-CoV-2 RBD - ACE2 receptor complex (PDB ID: 6M0J) was used as the wild type reference (Lan et al., 2020). The structure was optimized using CHARMM36m force field in GROMACS version 5.1.2. Three RBD double mutant L452R/T478K (delta), L452R/E484Q (Kappa) and E484K/N501Y (beta, gamma), and 5 RBD single mutants of L452R, T478K, E484Q, E484K, and N501Y included in the double mutants were used in the study. The structures with the mutated residues in each mutant were generated using UCSF Chimera following the default parameters (Pettersen et al., 2004).

### Molecular dynamics simulation

GROMACS version 5.1.2, was used for molecular dynamics simulation (Berendsen et al., 1995). CHARMM36m force field was chosen to model RBD - ACE2 receptor complex (Huang et al., 2017). The complex was situated in the simulation box 2 nm away from the box edge. The system was solvated with tip3p water and neutralized with Na+ ions. Steep descent algorithm was applied to the system before 1 ns equilibration run at 298 K and 1 bar in the NPT ensemble using Berendsen thermostat and barostat. The system was set at 298 K and 1 bar in the NPT ensemble by using a V-rescale thermostat and Parrinello-Rahman barostat during the production run (Parrinello & Rahman, 1981). Verlet velocity algorithm was employed with a time step of 2 fs. The Particle Mesh Ewald (PME) method was used to treat the long-range electrostatic interactions with the cut-off distance at 1.0 nm. Hydrogen bond was constrained at equilibrium lengths by the LINC algorithm and the trajectory frame of MD was saved every 30 ps (Hess, 2008). RMSD, RMSF, Rg, SASA and MSD were used to analyze structural changes of the native RBD and its mutants. The 35-40ns simulation trajectories were utilized in each method. XMGRACE program was utilized to generate the corresponding plots (Turner, 2005). Three independent RMSD MD simulations were performed for the wild type RBD-ACE2 for 100 ns, received RBD values at 0.186 nm, 0.173 nm, 0.179 nm, ACE2 values at 0.229 nm, 0.230 nm, 0.239 nm for run 1, run 2, run 3. The results confirmed that RBD - ACE2 structure remained stable throughout the simulation process.

### Superimposed structural comparison

Mutant RBD structures extracted at 100 ns of MD simulations through UCSF CHIMERA were superimposed with the wild type RBD structures to identify the conformation changes using PYMOL.

### Antibody escaping analysis

To determine whether RBD mutants escape the binding by neutralization antibody, structures between the mutant and wildtype RBD were structurally mapped with independent binding sites of antibody C121 on RBD. The RBD binding site for antibody C121, a neutralizing antibody known to bind RBD (PDB: 7K8X) (Barnes et al., 2020; Weisblum et al., 2020), and E484K mutant known to escape C121 binding were used to determine the impact of double mutant L452R/T478K, L452R/E484Q and E484K/N501Y on antibody binding structure.

### Free binding energy calculation

MM/GBSA (molecular mechanics energies combined with generalized Born and surface area continuum solvation) (Genheden & Ryde, 2015) was used to calculate the free binding energy of the RBD to ACE2 (http://cadd.zju.edu.cn/hawkdock,(Weng et al., 2019). ff02 force field (Cieplak et al., 2001) and the GB^OBC1^ model (Onufriev et al., 2004) were assigned upon the proteins (Complex, RBD and ACE2) before steepest descent and conjugate gradient minimization. The free energy of binding, Δ*G*, is calculated based on the following equation:

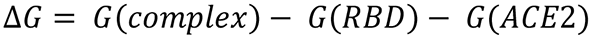

For each component, the total energy was estimated by using the following equation:

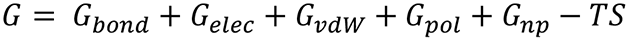

The first three terms represent the standard MM energy terms: bonded (bond, angle, and dihedral), electrostatic and van der Waal interactions. *G*_*pol*_ and *G*_*np*_ are the polar and the non-polar contribution of solvation free energies, calculated by the Generalised Born (GB) implicit solvent method and solvent accessible surface area (SASA). The last term absolute temperature, *T*, is multiplied by entropy, *S*. For the double mutant RBD and their containing single mutant RBD, simulation was performed at 100 ns and a single protein coordinate file was extracted every 10 ns. Overall free binding energy was averaged for each type of RBD.

## Acknowledgements

We are thankful for the Information and Communication Technology Office (ICTO), University of Macau for providing the High-Performance Computing Cluster (HPCC) resource and facilities for the study.

## Contribution

SS, BT: method development, analysis, data acquisition and interpretation, draft manuscript; SMW: conception, design, data analysis and interpretation, draft and revise manuscript, and funding.

## Funding

This work was supported by Macau Science and Technology Development Fund (085/2017/A2, 0077/2019/AMJ), University of Macau (SRG2017-00097-FHS, MYRG2019-00018-FHS), Faculty of Health Sciences, University of Macau (FHSIG/SW/0007/2020P, Startup fund) (SMW).

## Competing interests

None declared.

## Patient consent for publication

Not required.

## Ethics approval

Not required.

## Data availability

All data relevant to the study are included in the article or uploaded as online supplemental information.

## Additional Supplementary Files

**S. Figure 1.**
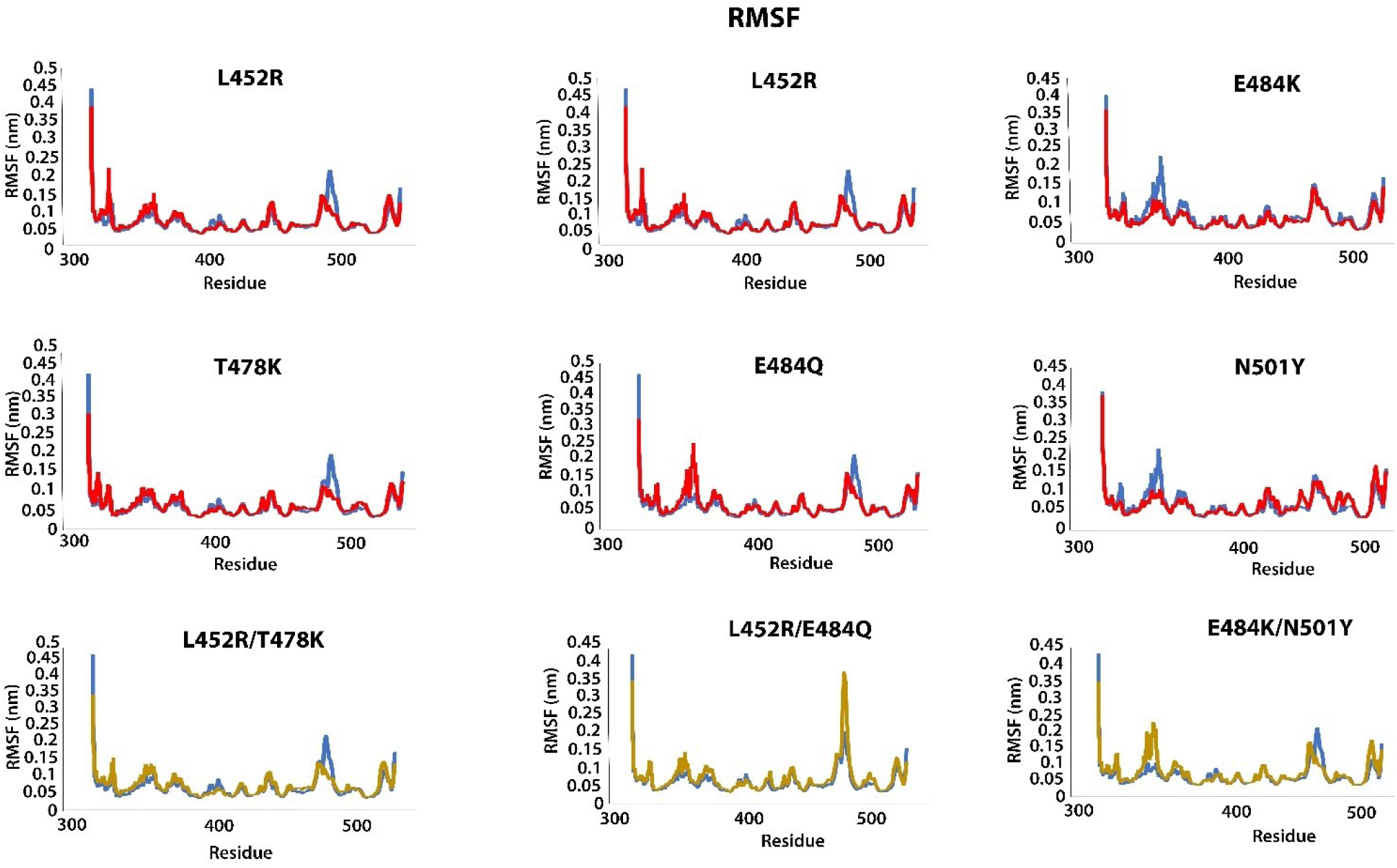
Dynamic changes of RBD structure by RMSF analysis in wildtype, single and double mutant structures. The x-axis represents the residue positions of the RBD domain and y-axis represents the RMSF value for the structures. Blue: wild type; red: single mutant; yellow: double mutants.

**S. Figure 2.**
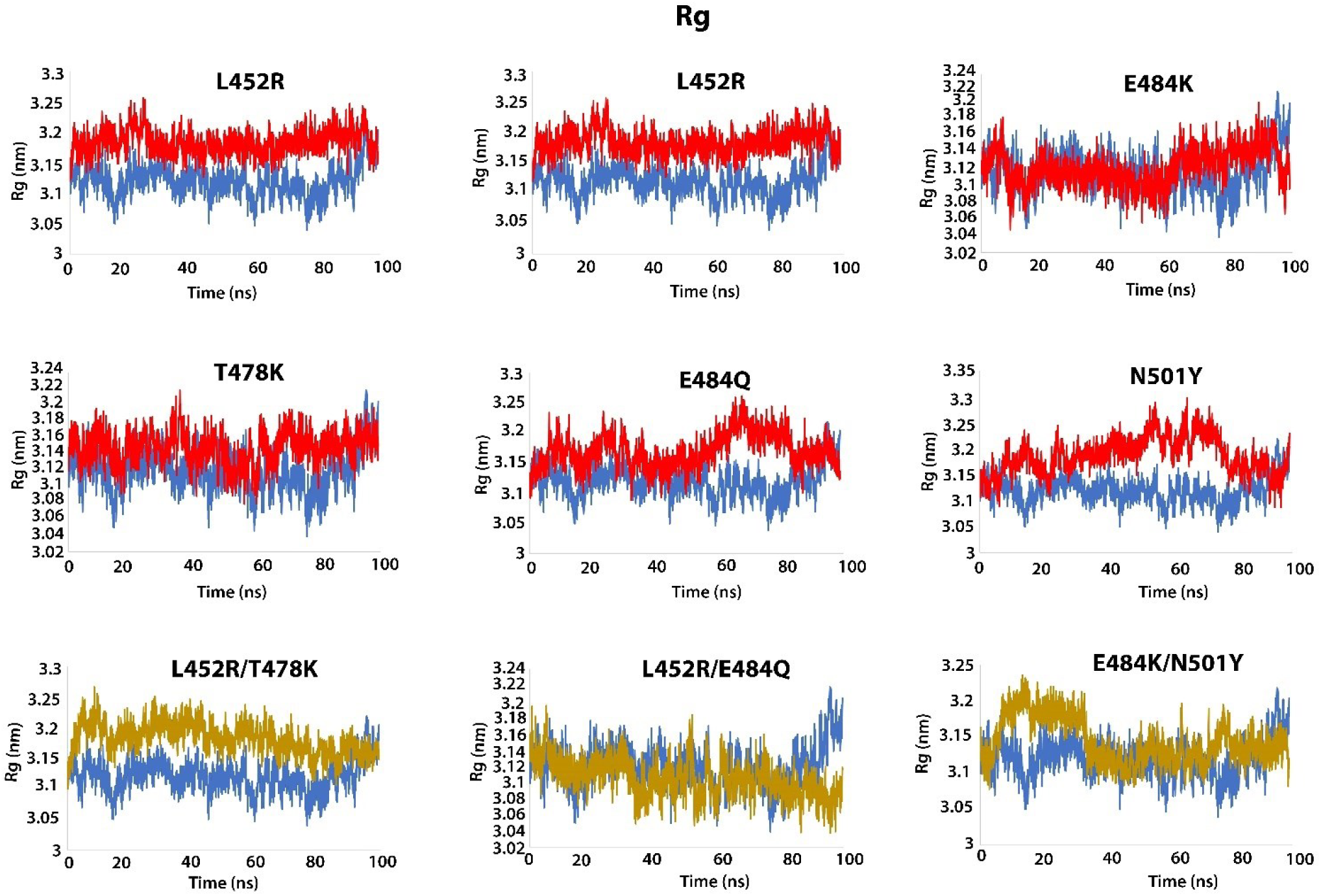
Dynamic changes of RBD structure by Rg analysis in in wildtype, single and double mutant structures. The x-axis represents the time period of 100 ns and y-axis represents the Rg value for the structures. Blue: wild type; red: single mutant; yellow: double mutants.

**S. Figure 3.**
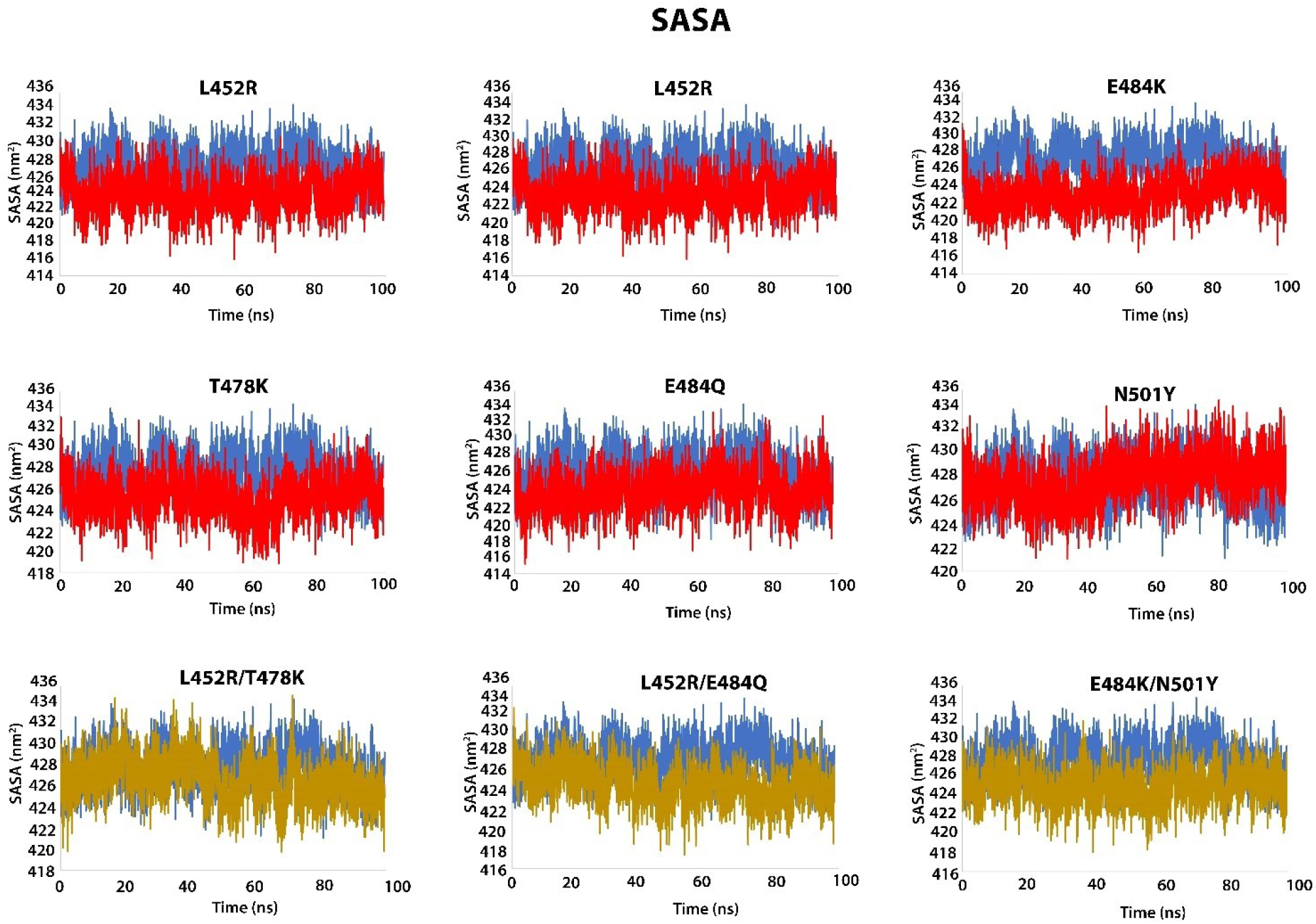
Dynamic changes of RBD structure by SASA analysis in wildtype, single and double mutant structures. The x-axis represents the time period of 100 ns and y-axis represents the SASA value for the structures. Blue: wild type; red: single mutant; yellow: double mutants.

**S. Figure 4.**
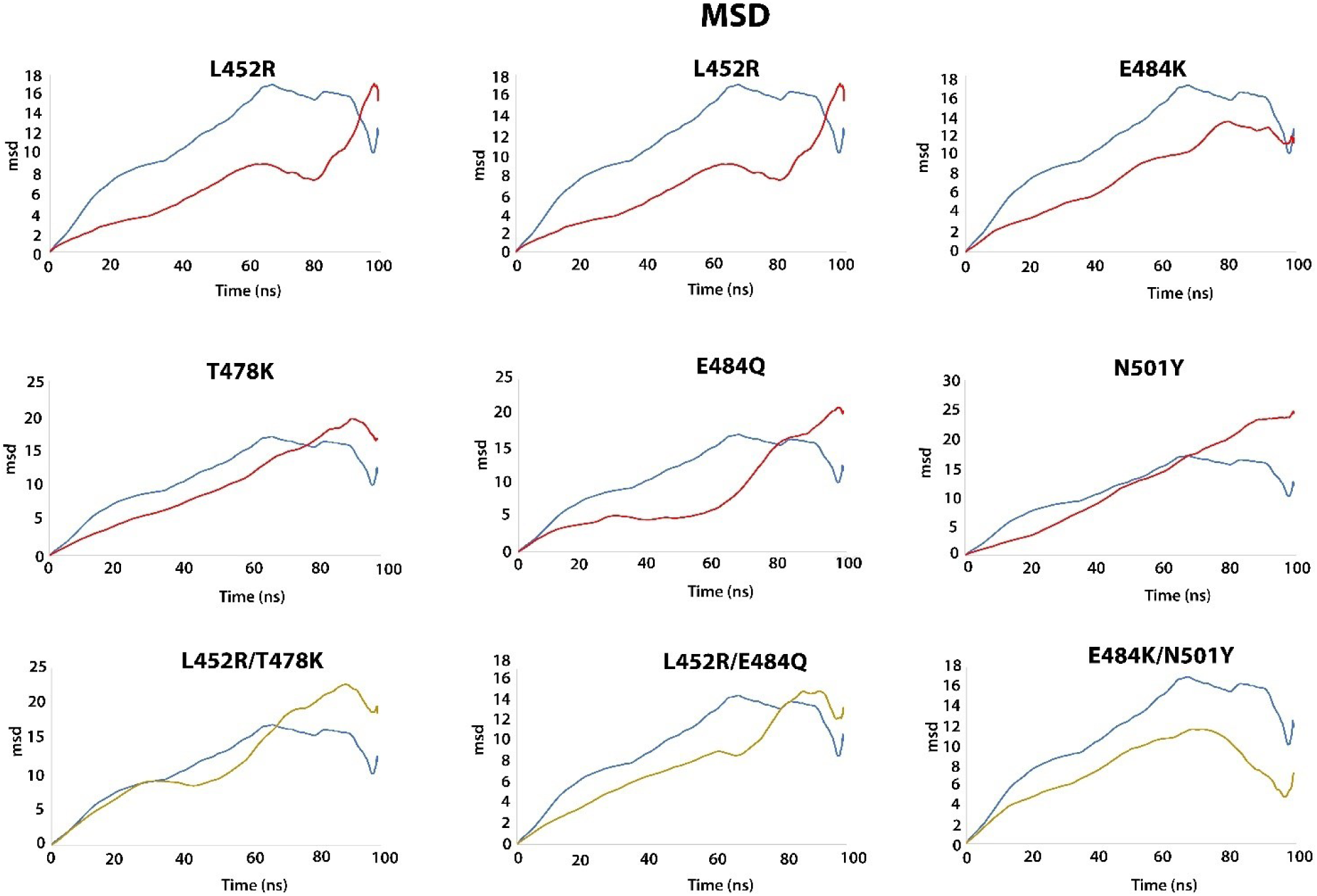
Dynamic changes of RBD structure by MSD analysis in wildtype, single and double mutant structures. The x-axis represents the time period of 100 ns and y-axis represents the MSD value for the structures. Blue: wild type; red: single mutant; yellow: double mutants.

**Supplementary table 1.**
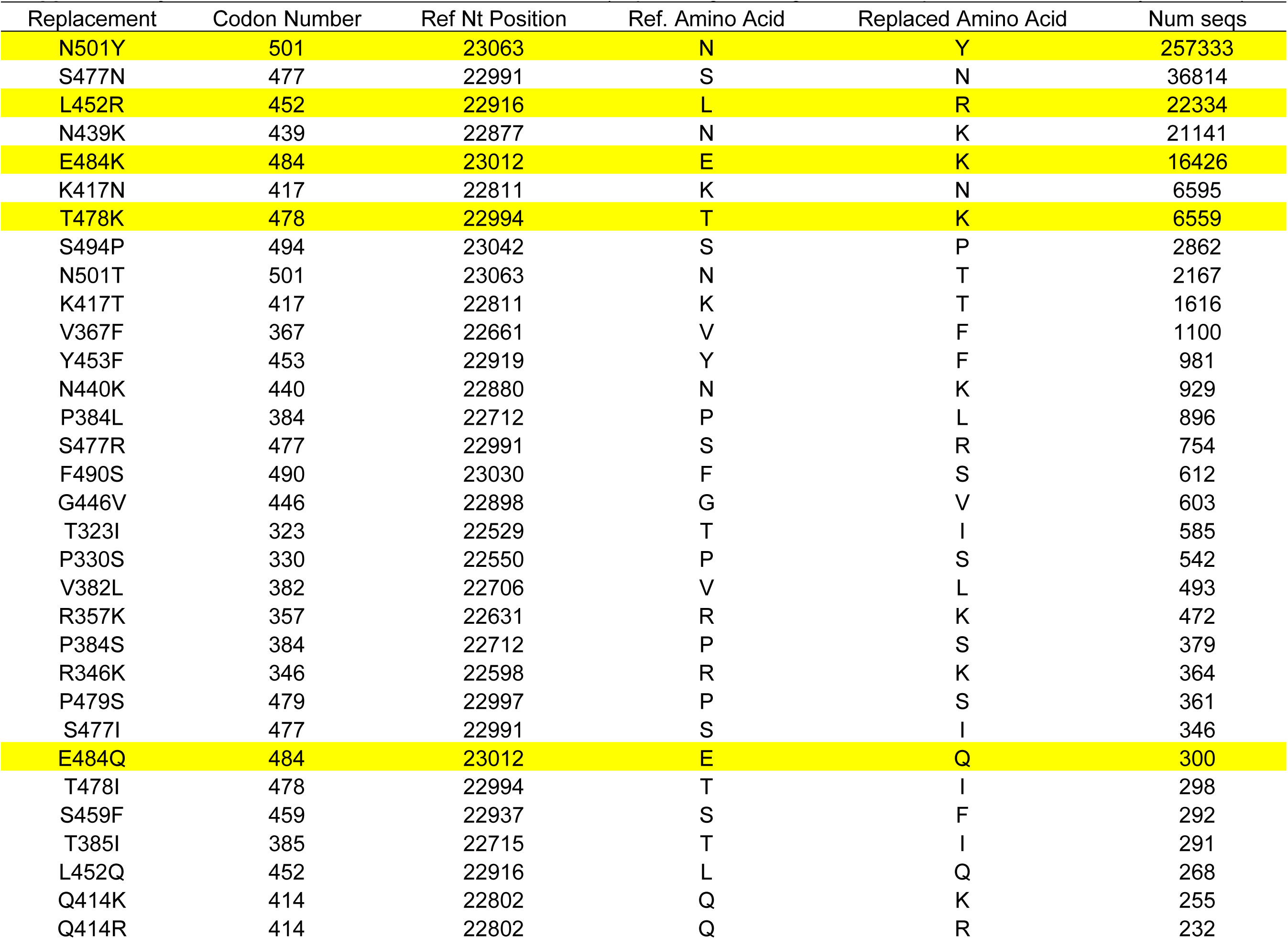
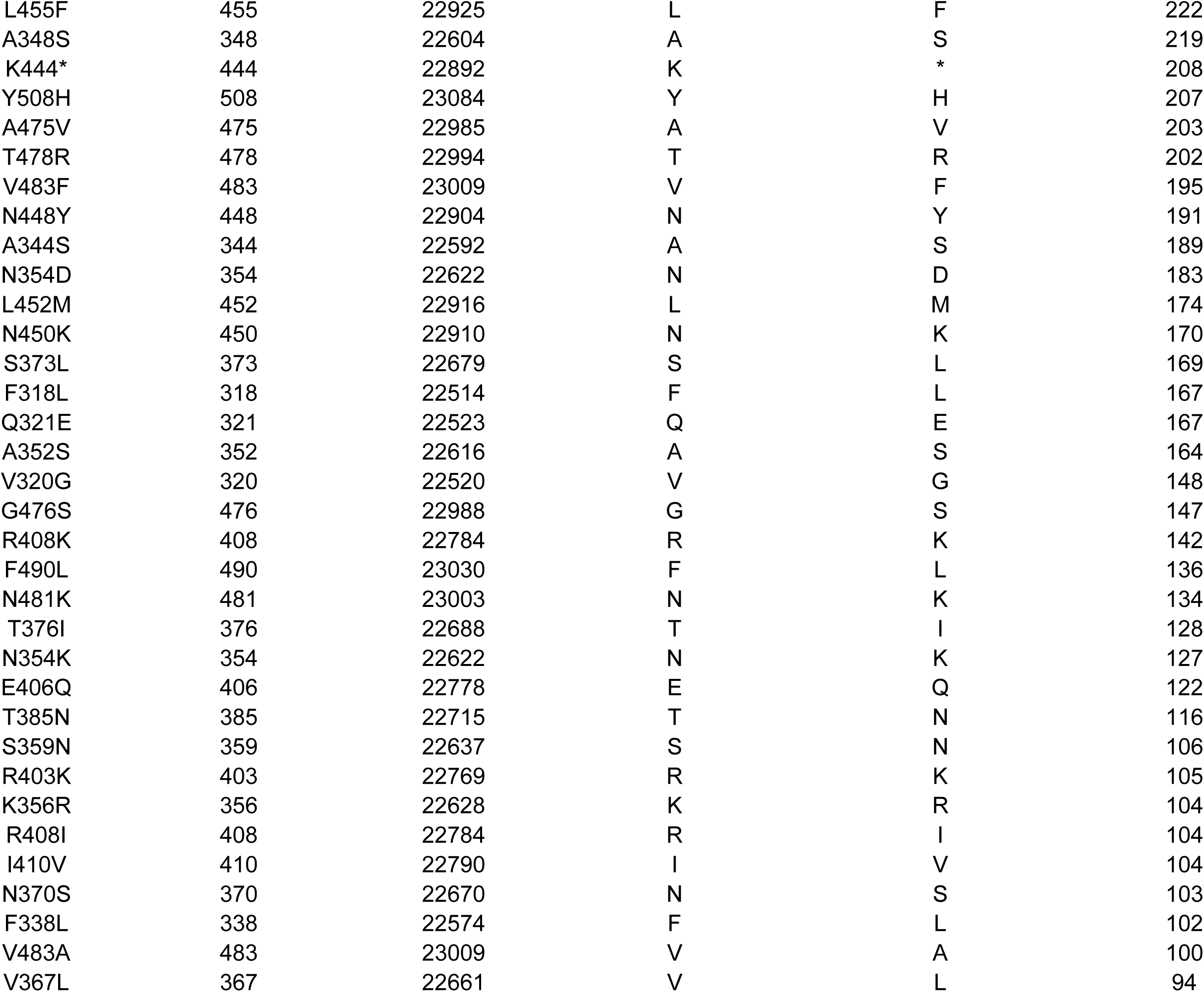
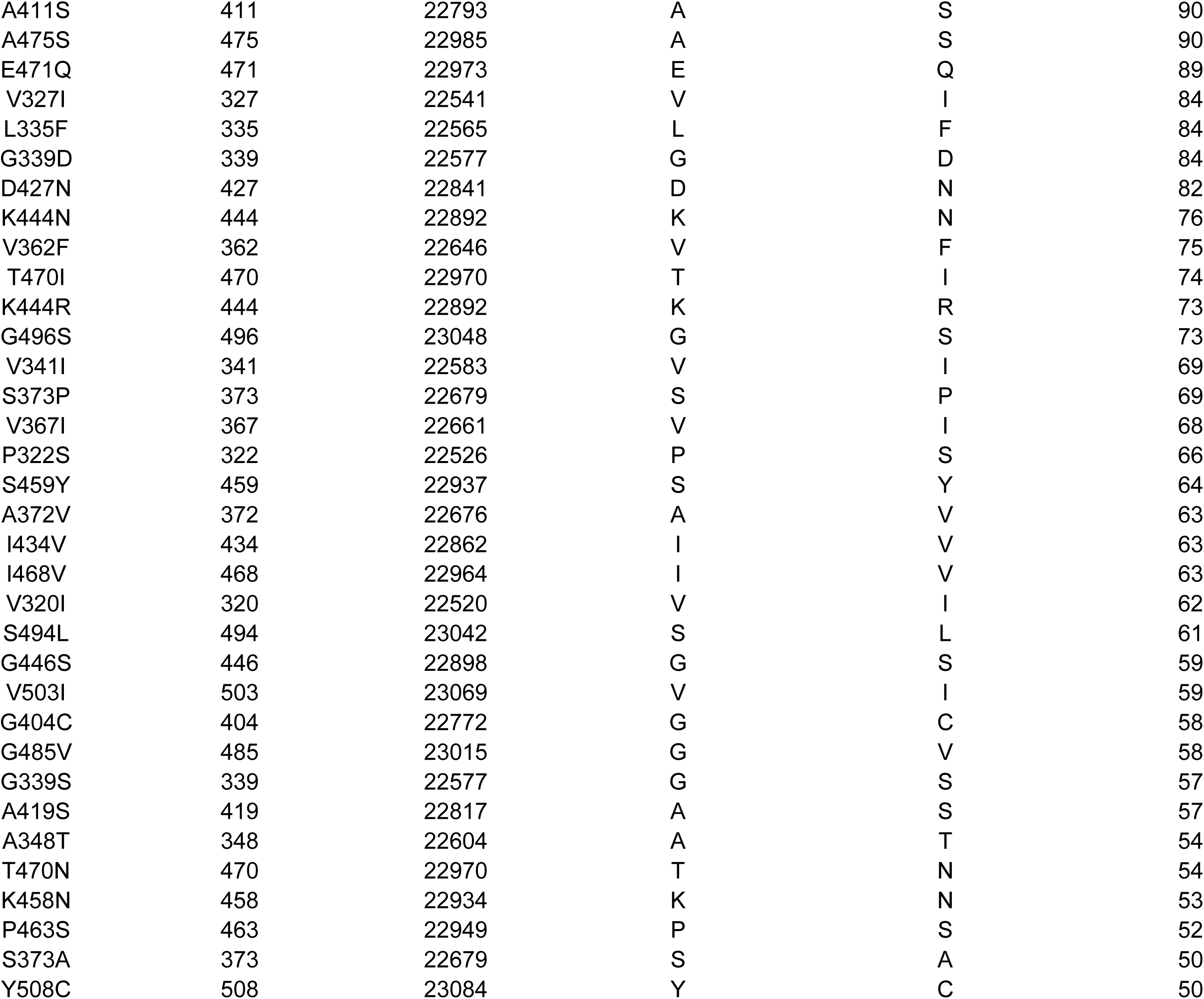
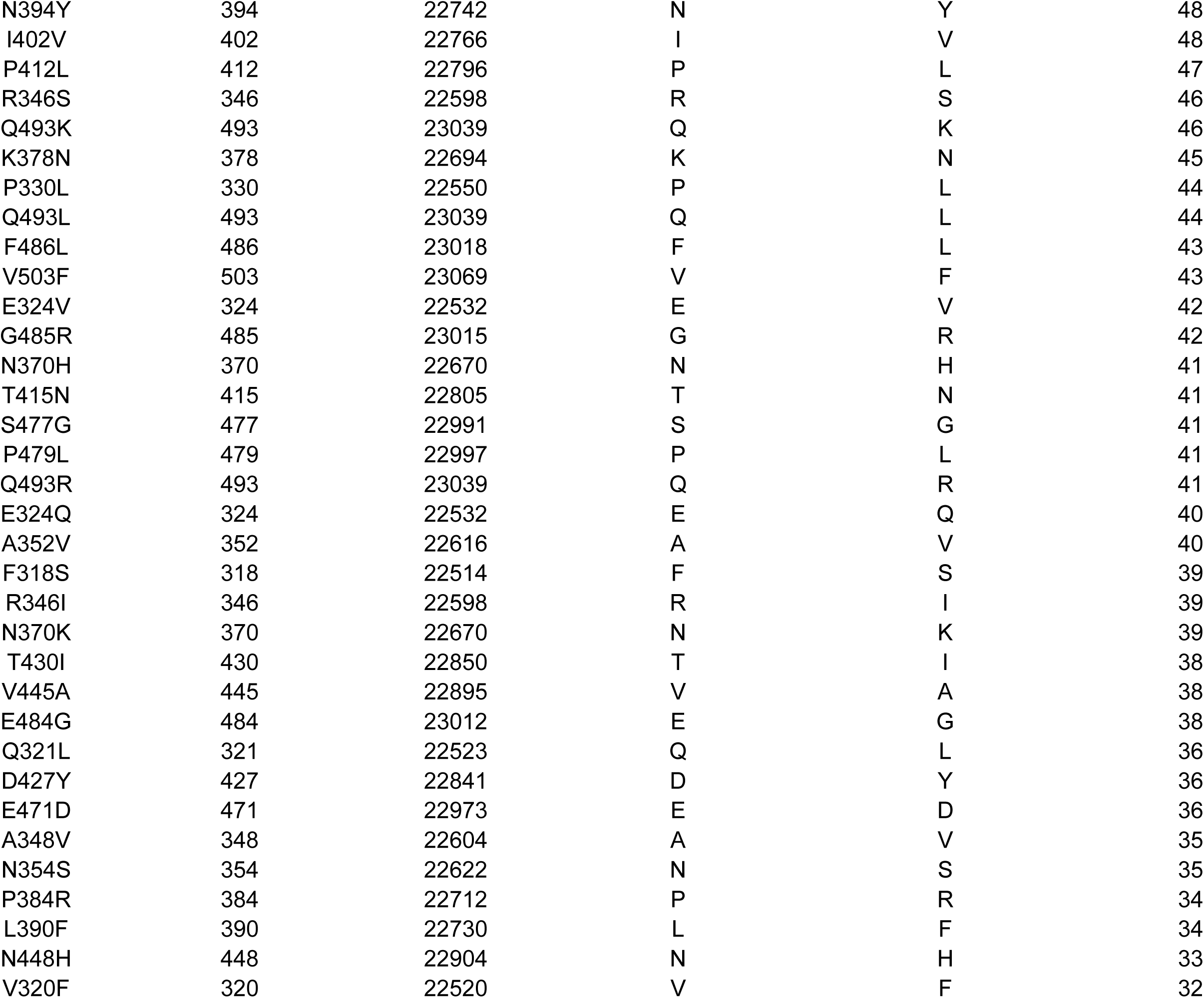
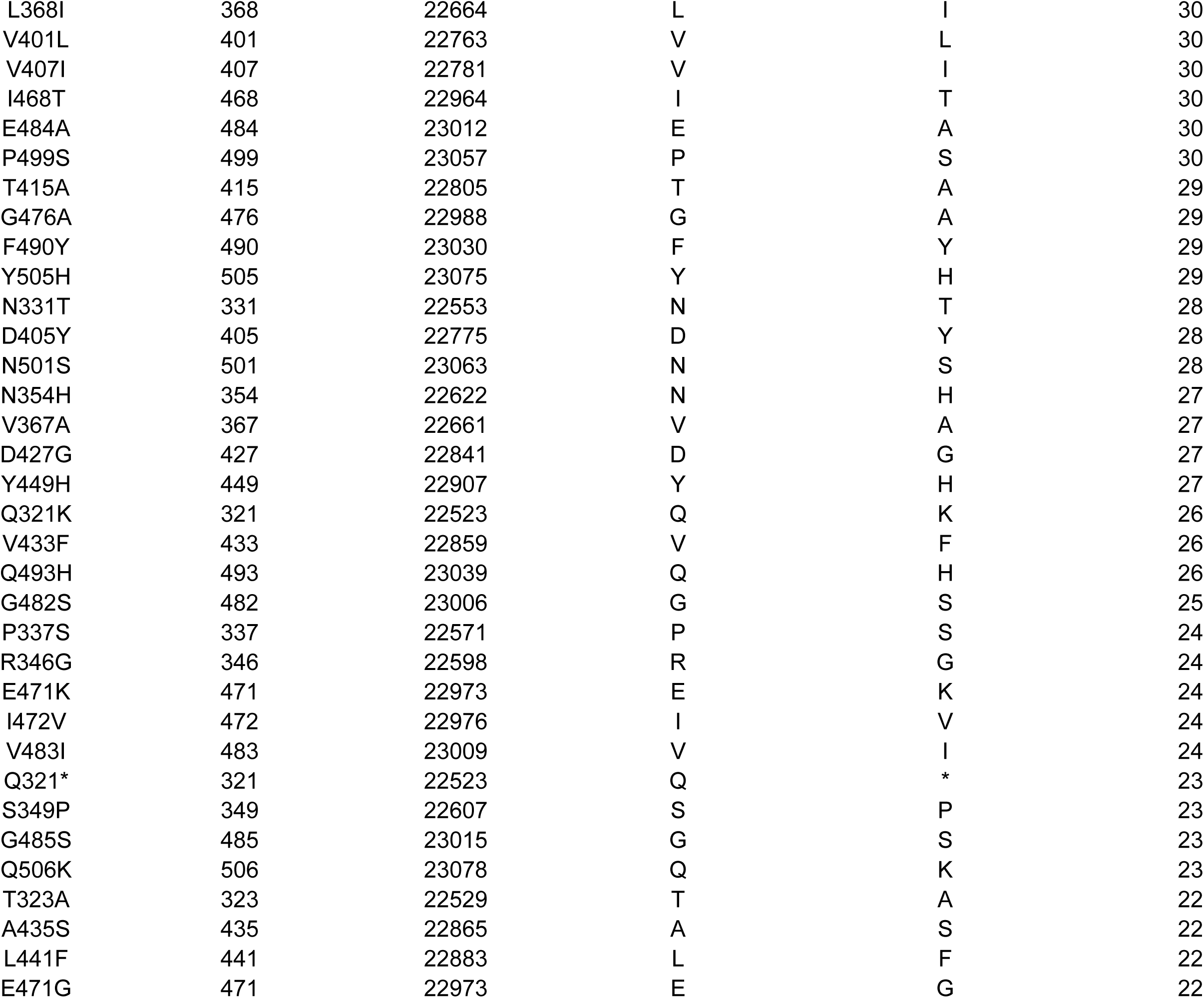
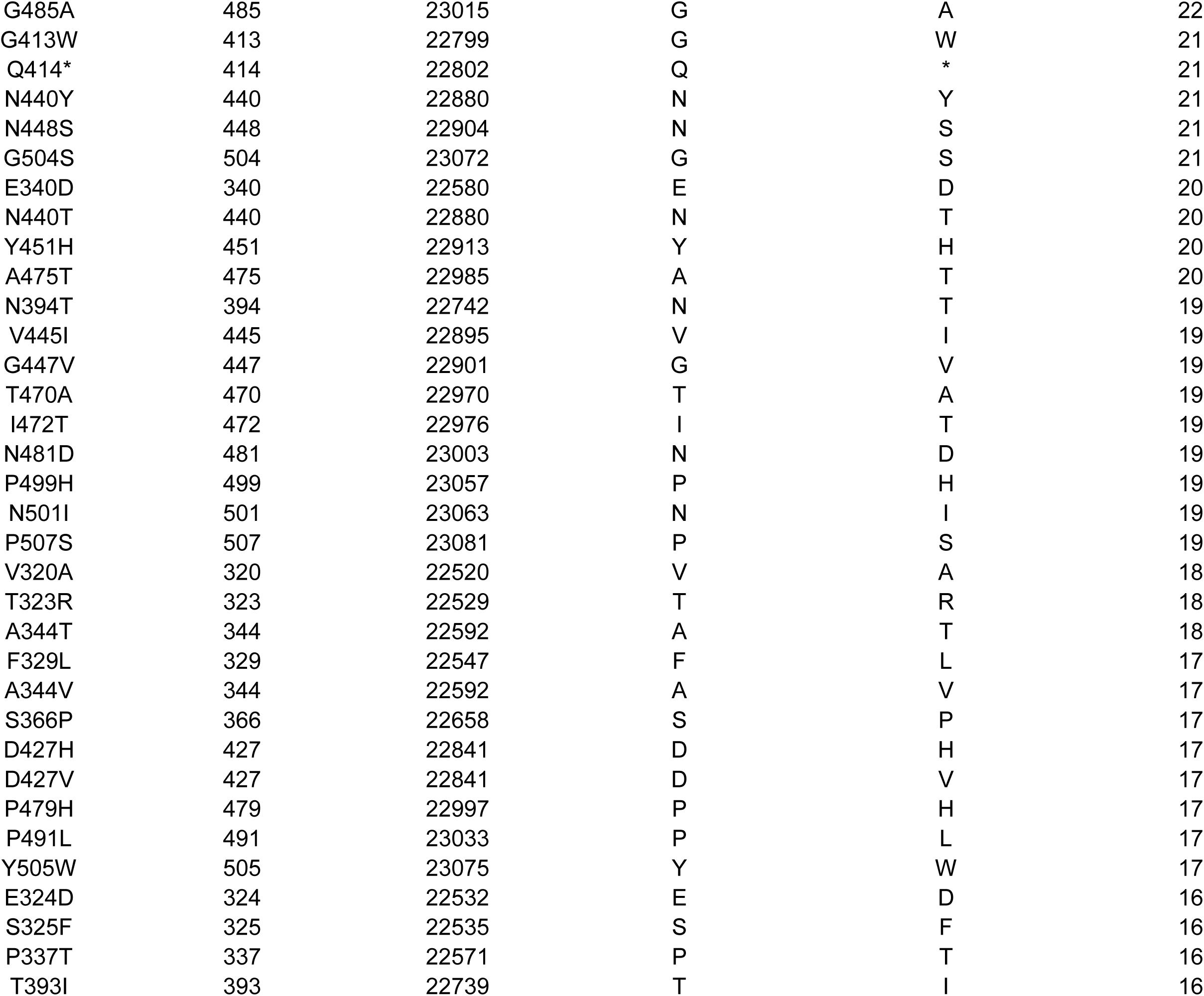
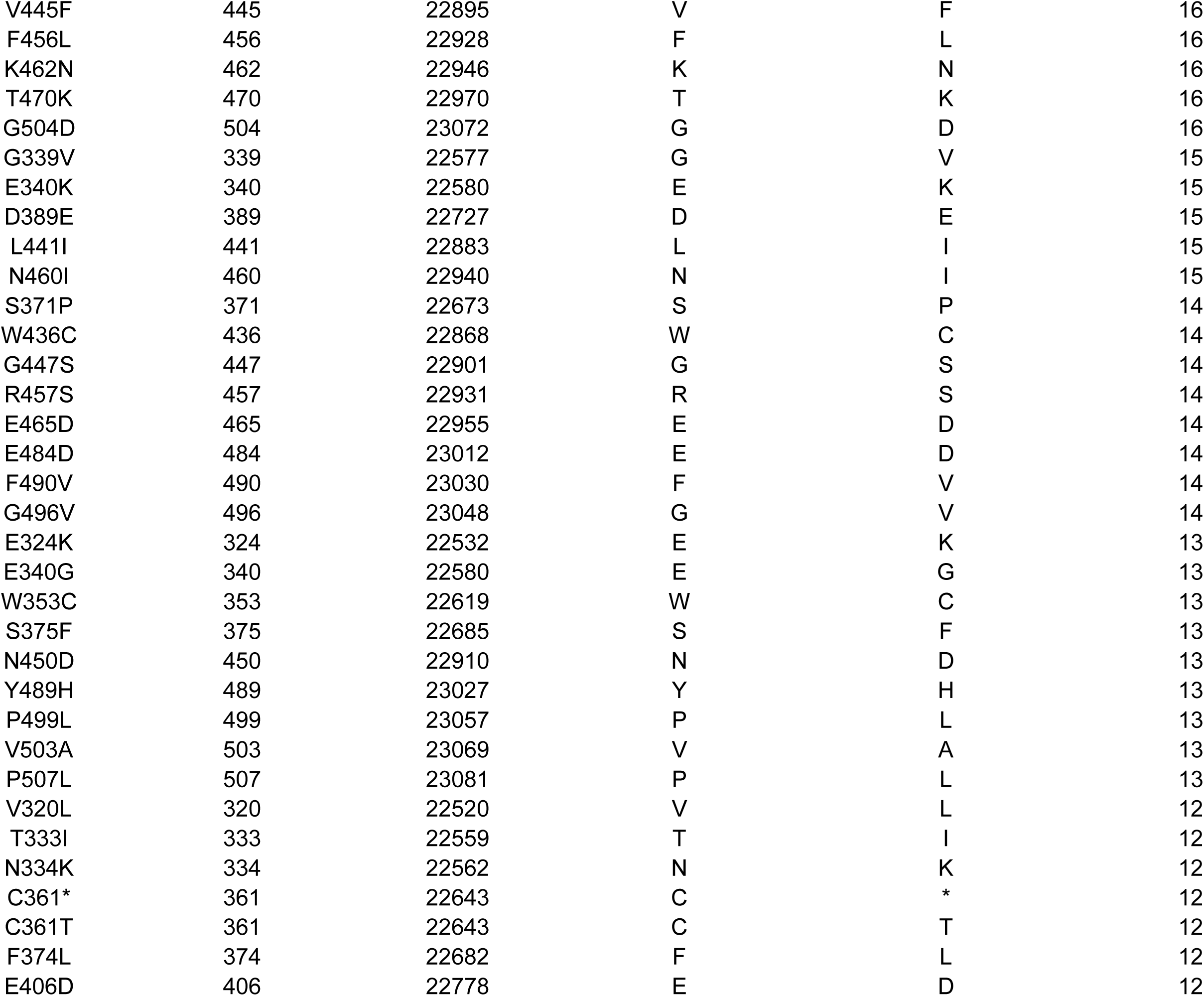
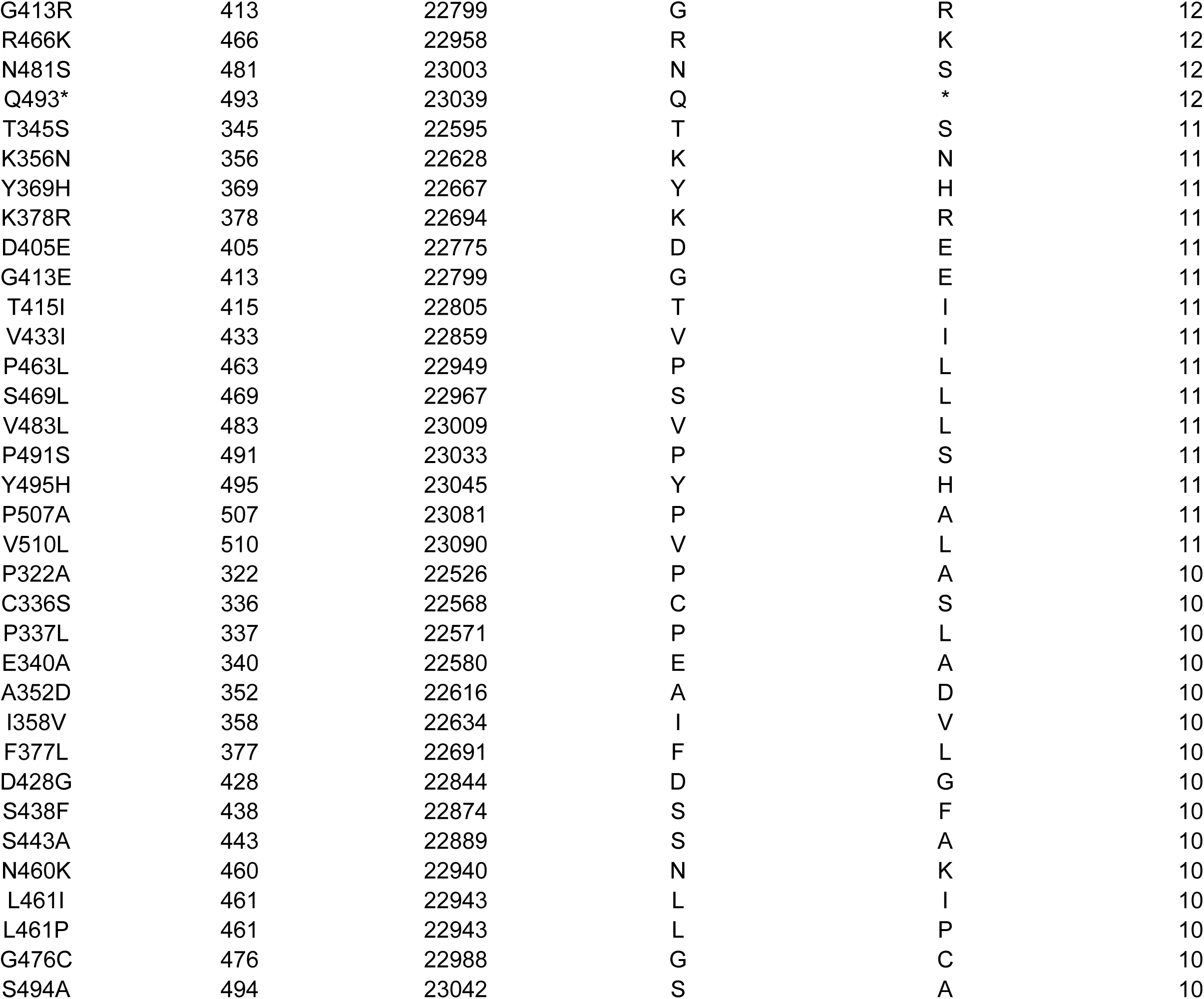
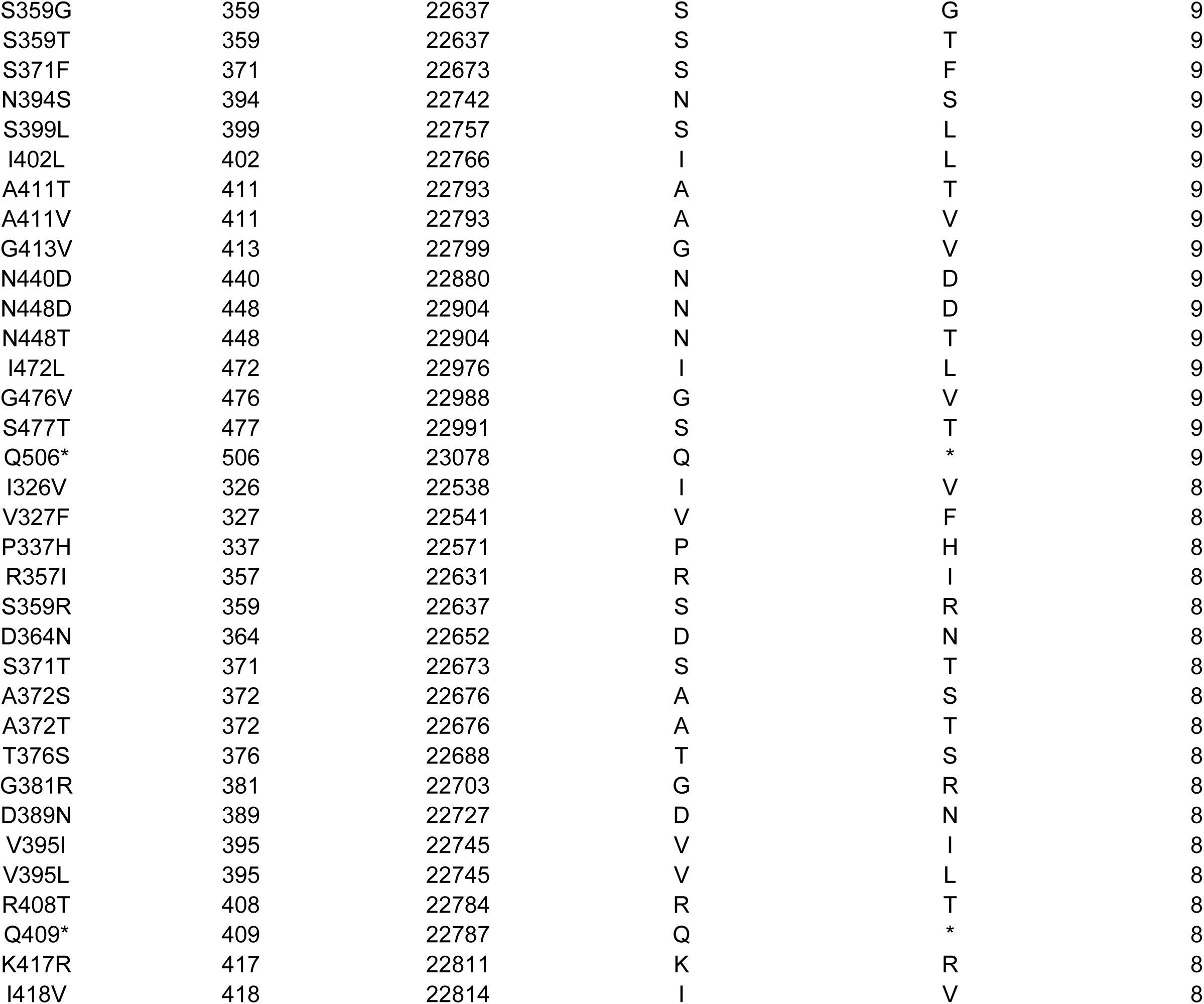
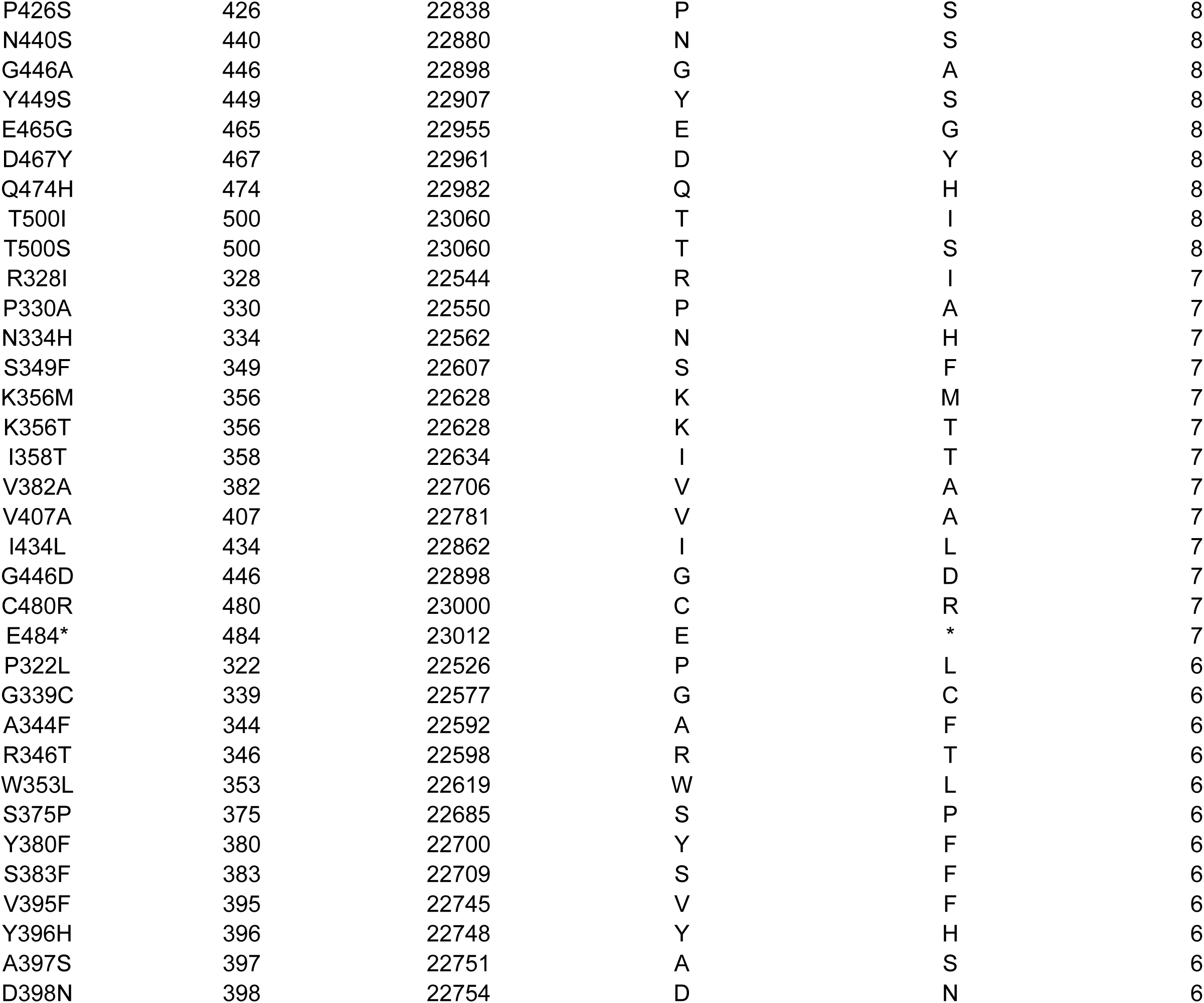
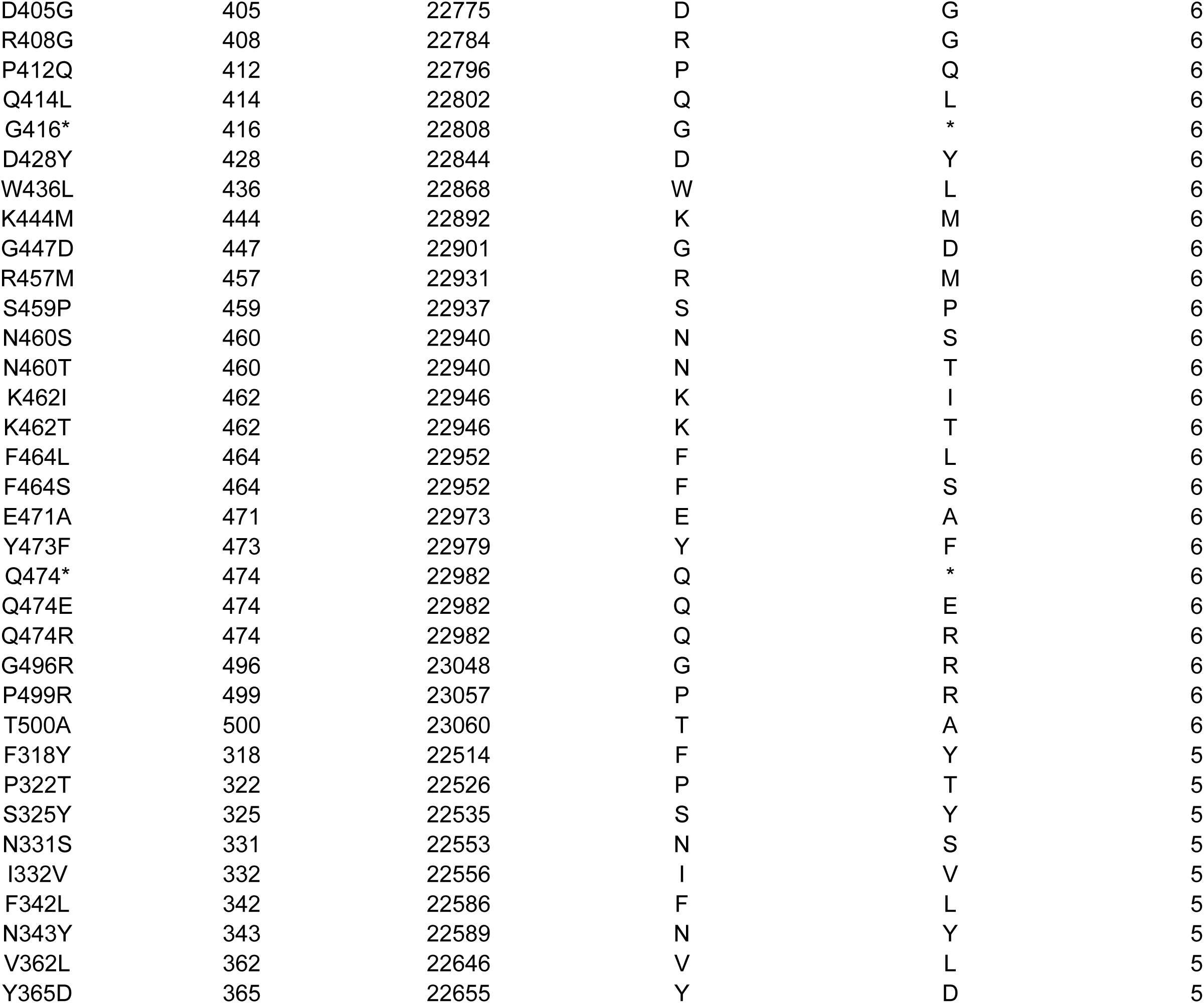
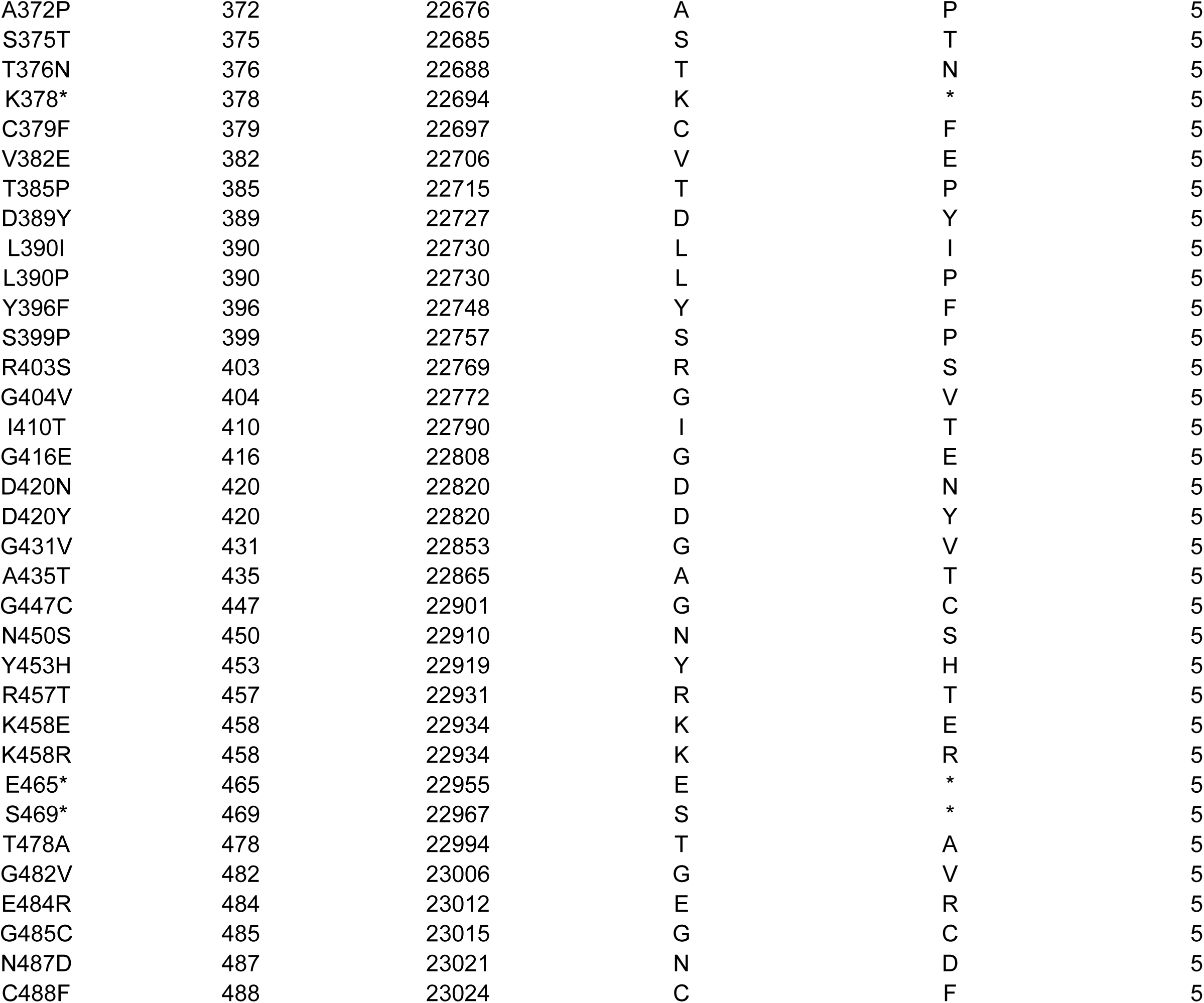
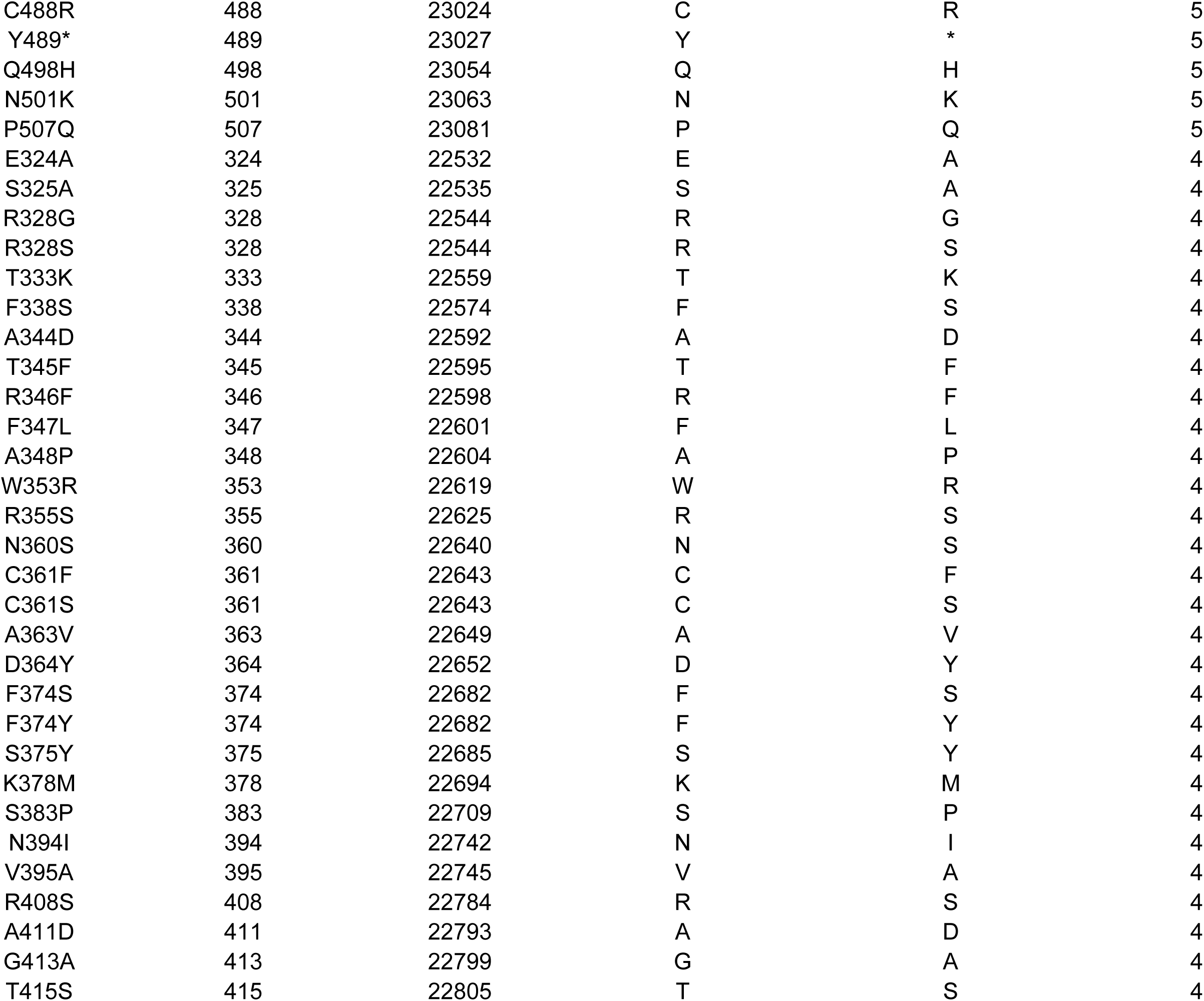
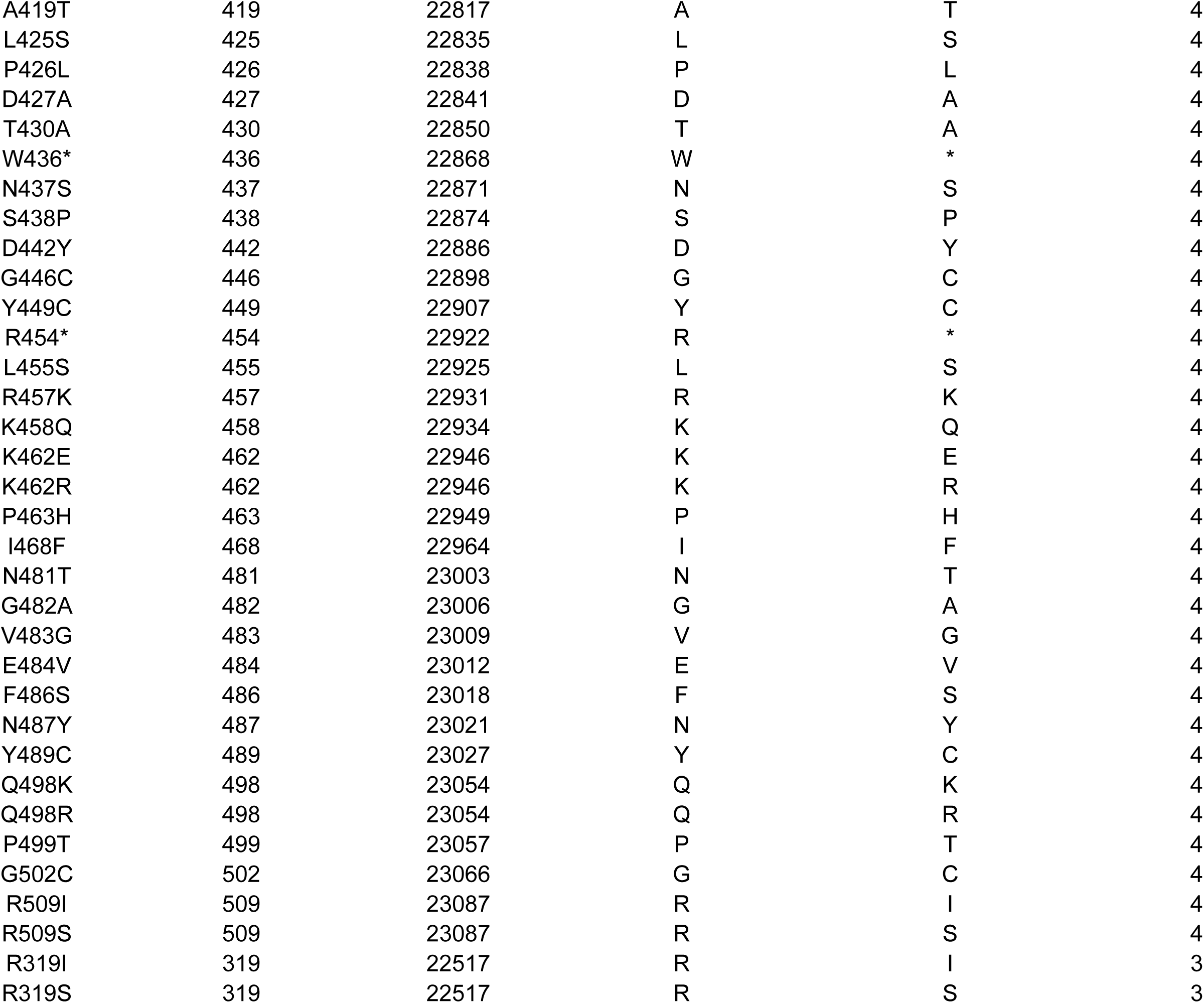
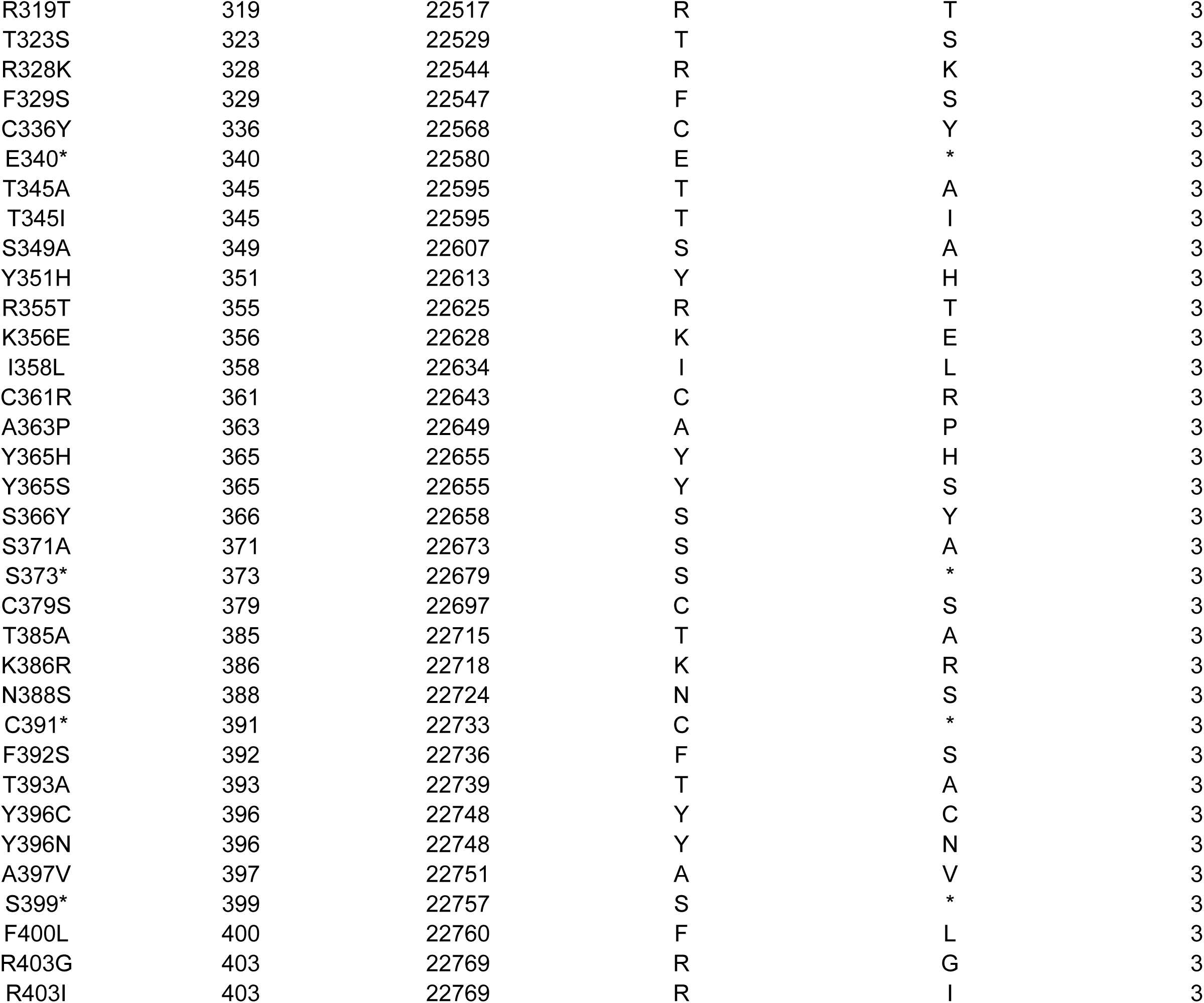
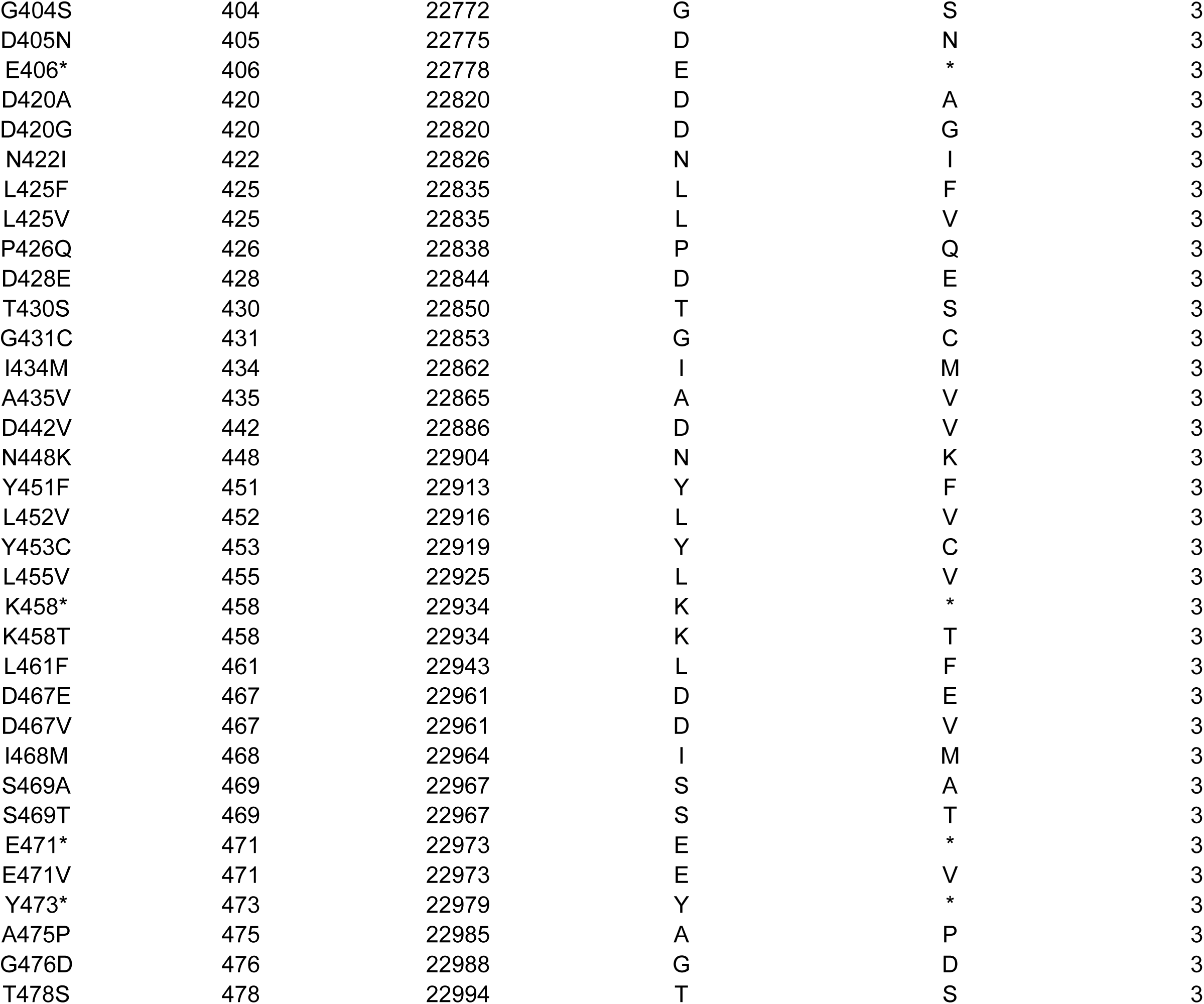
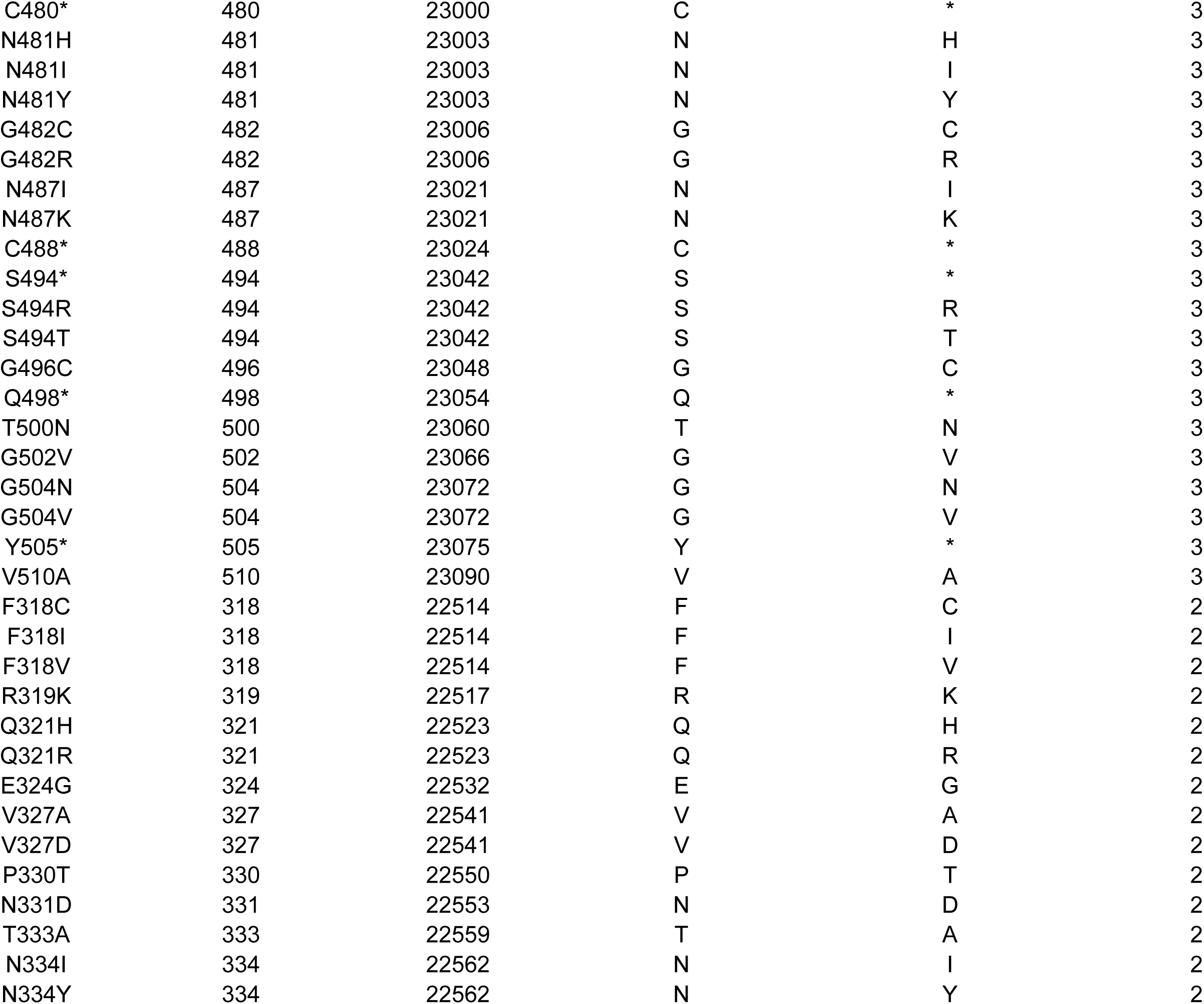
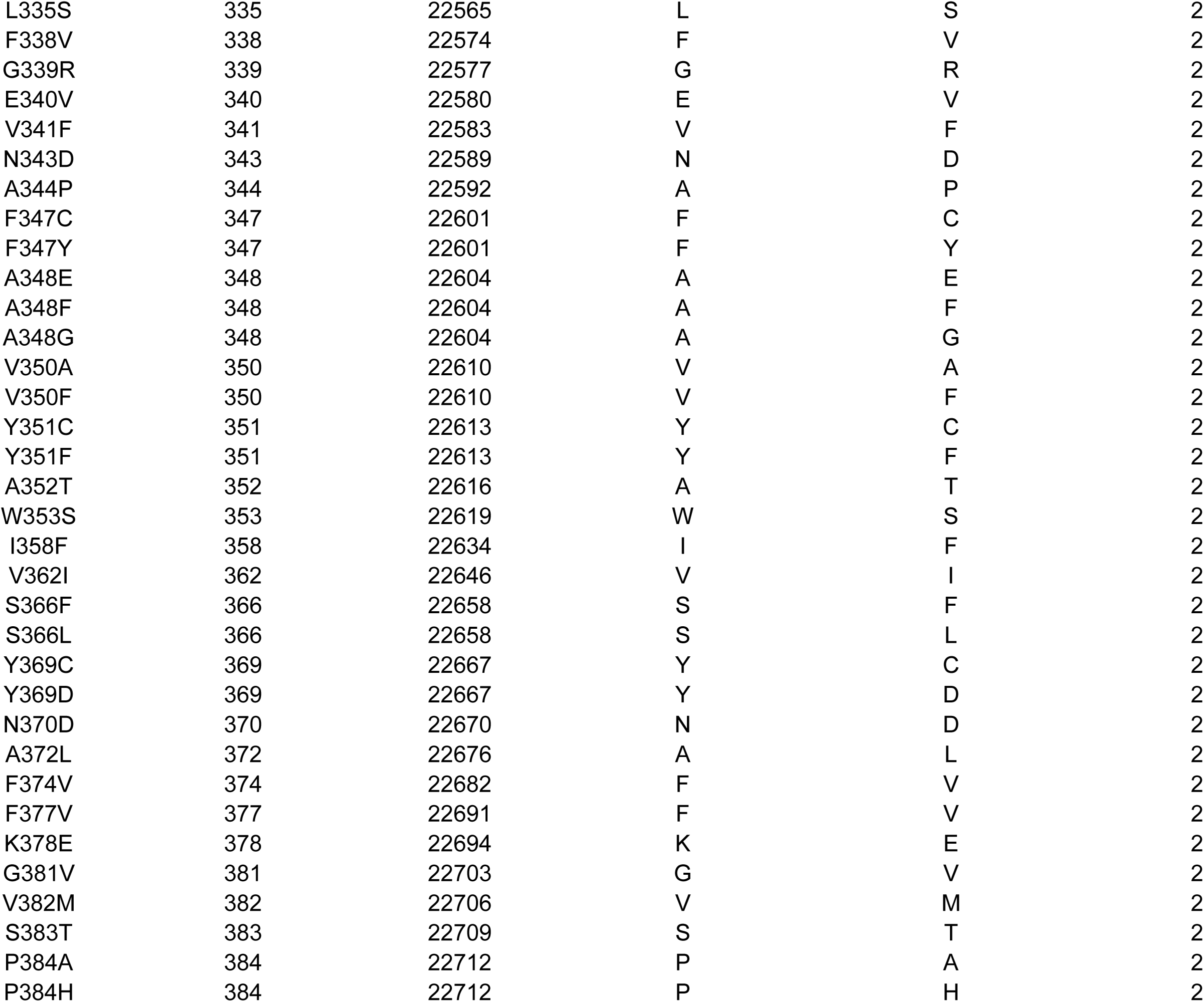
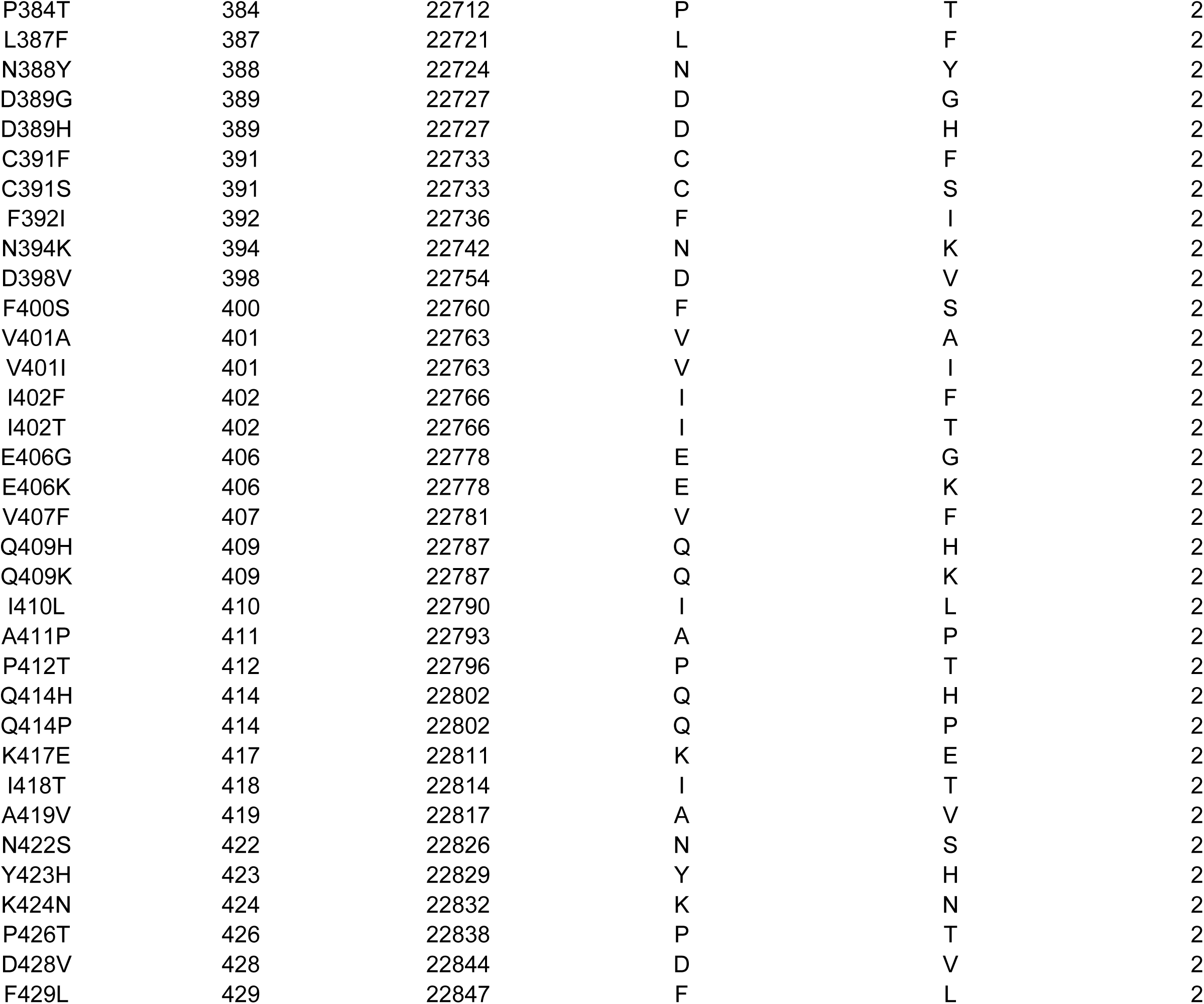
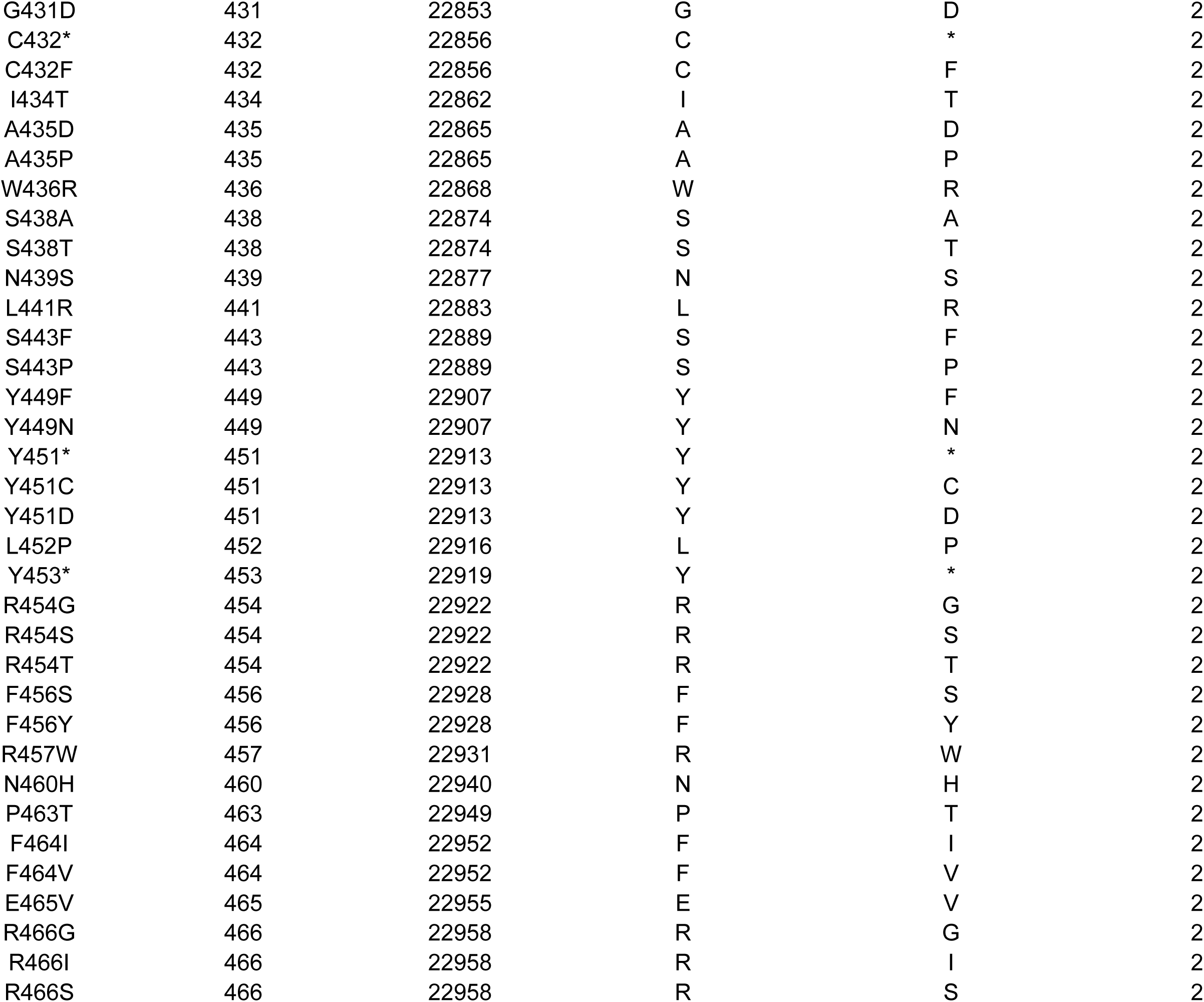
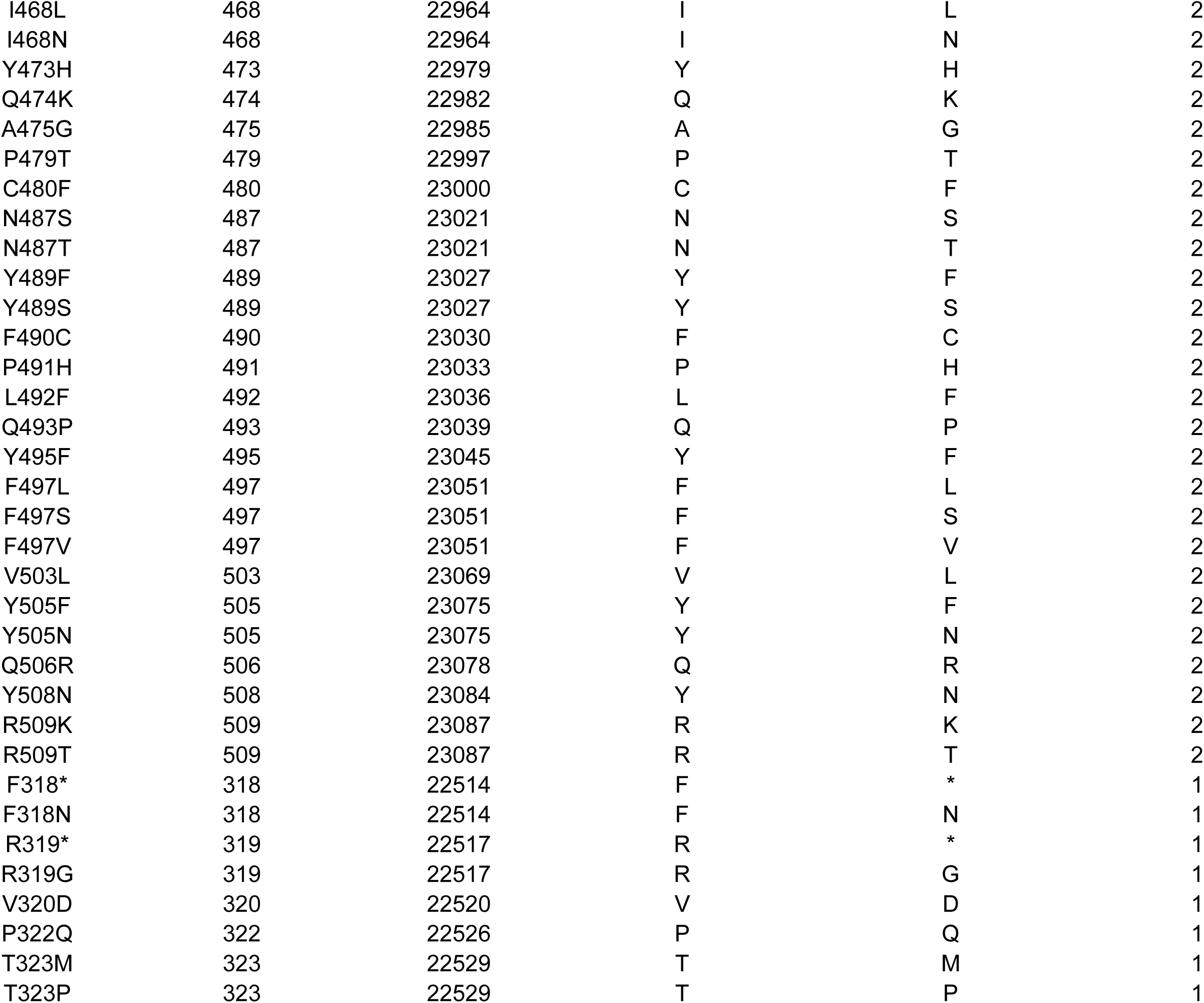
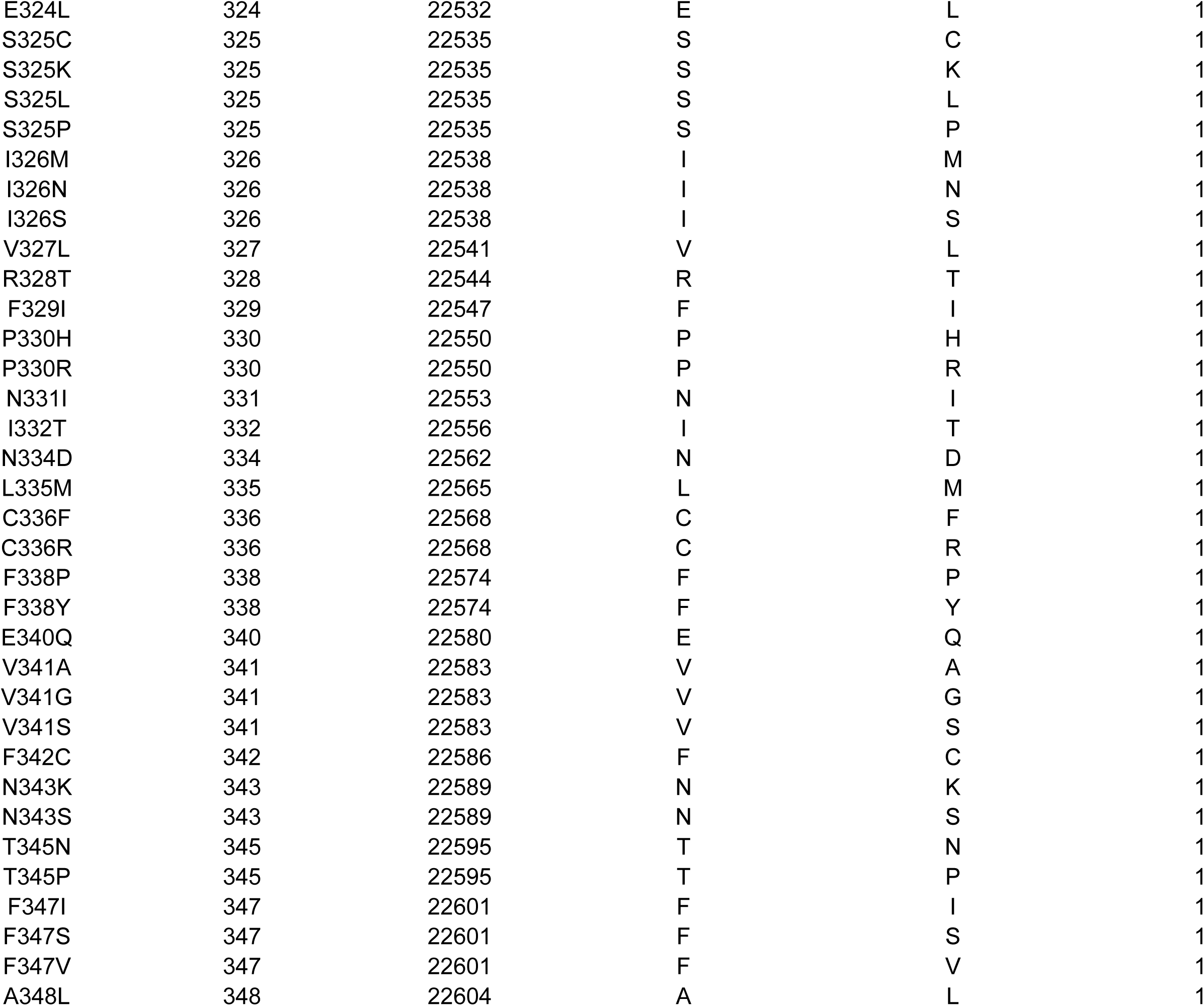
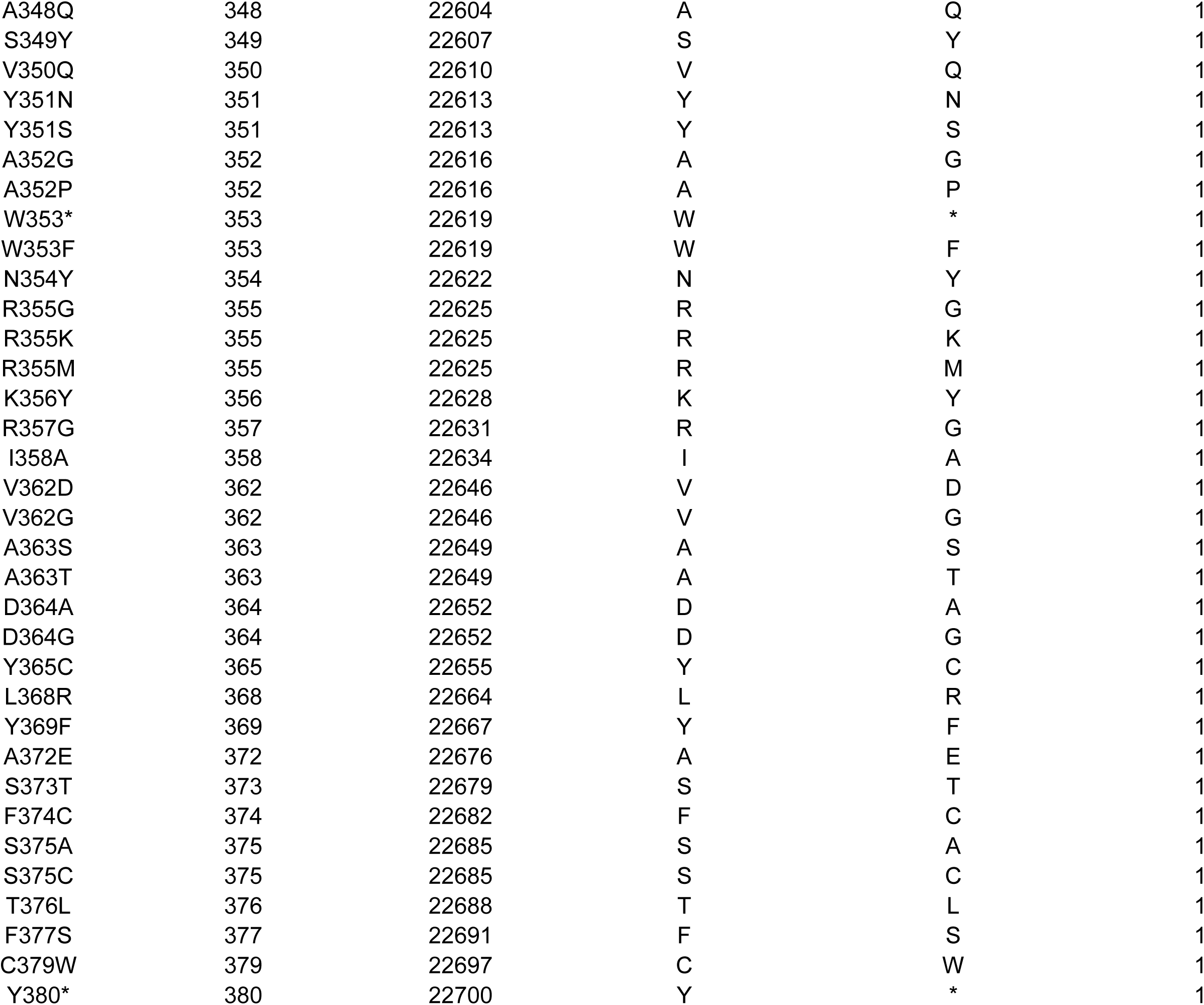
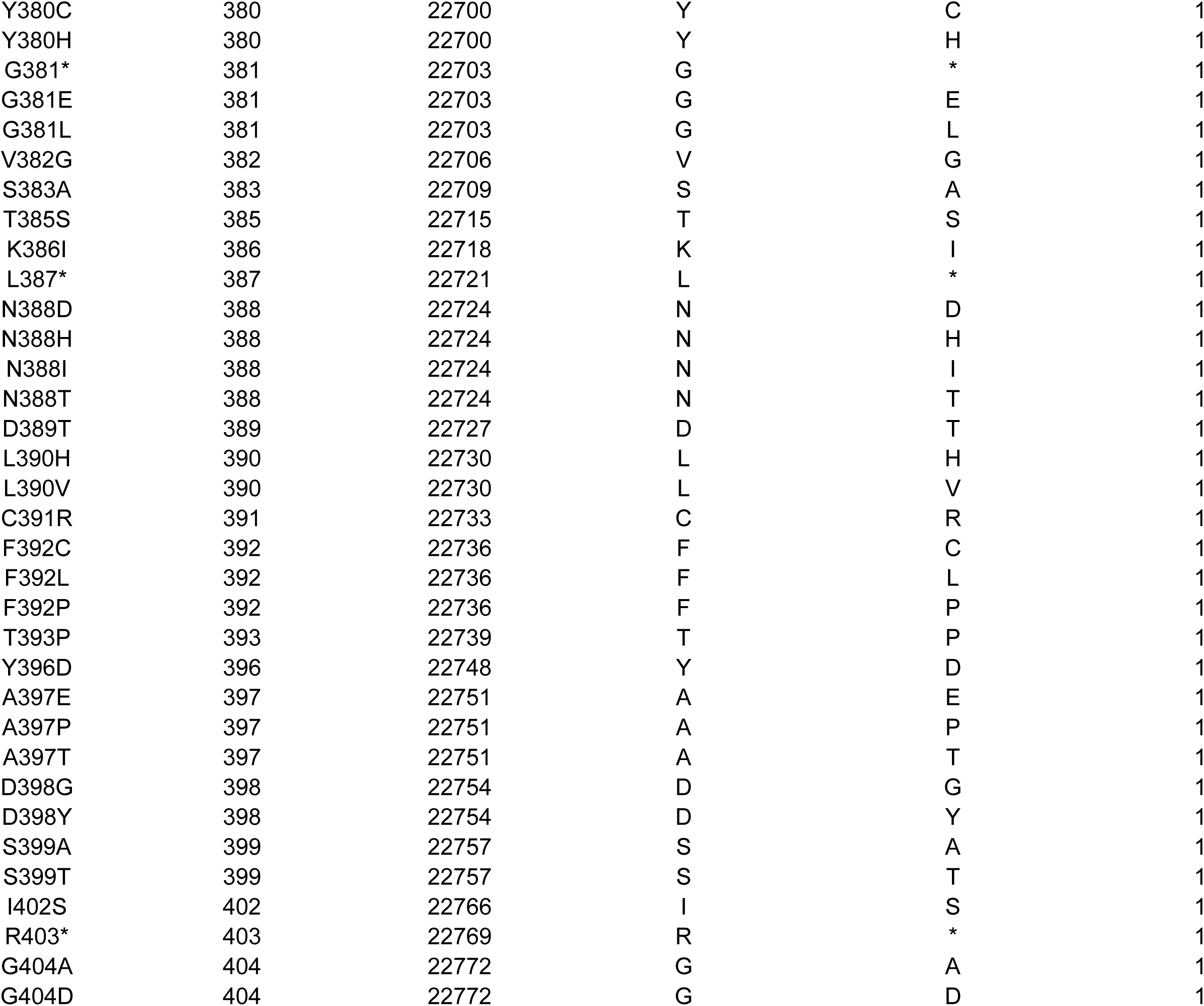
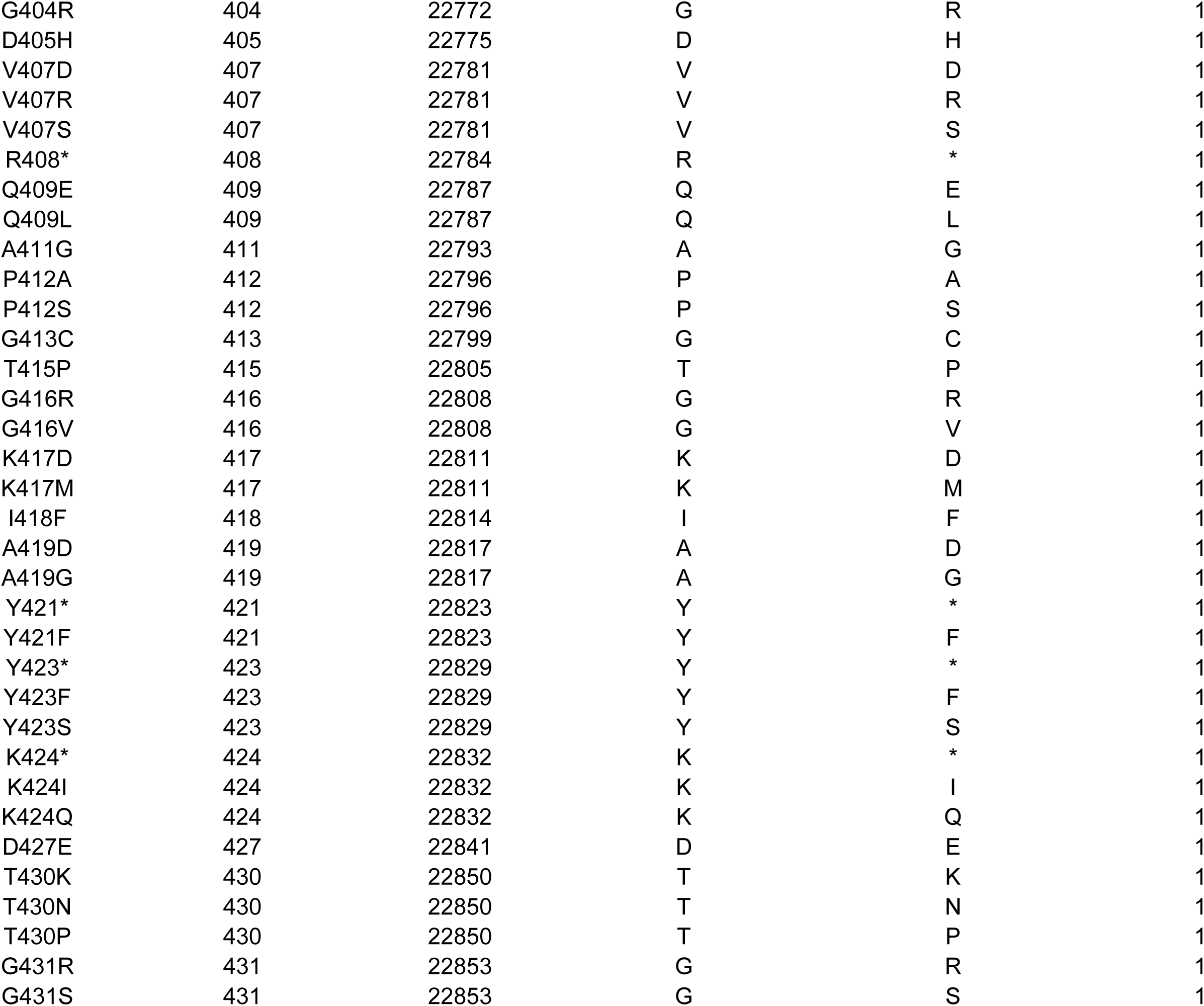
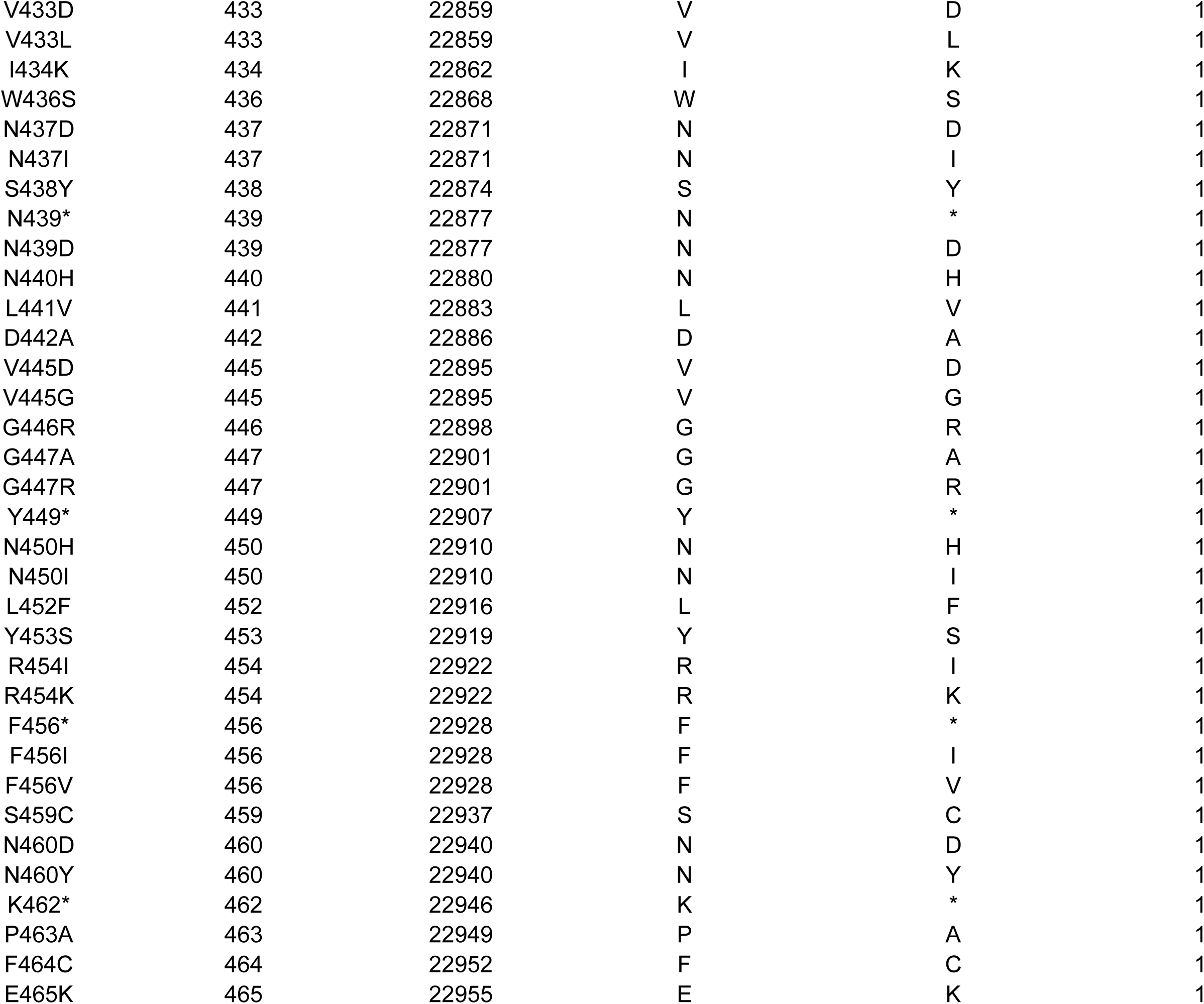
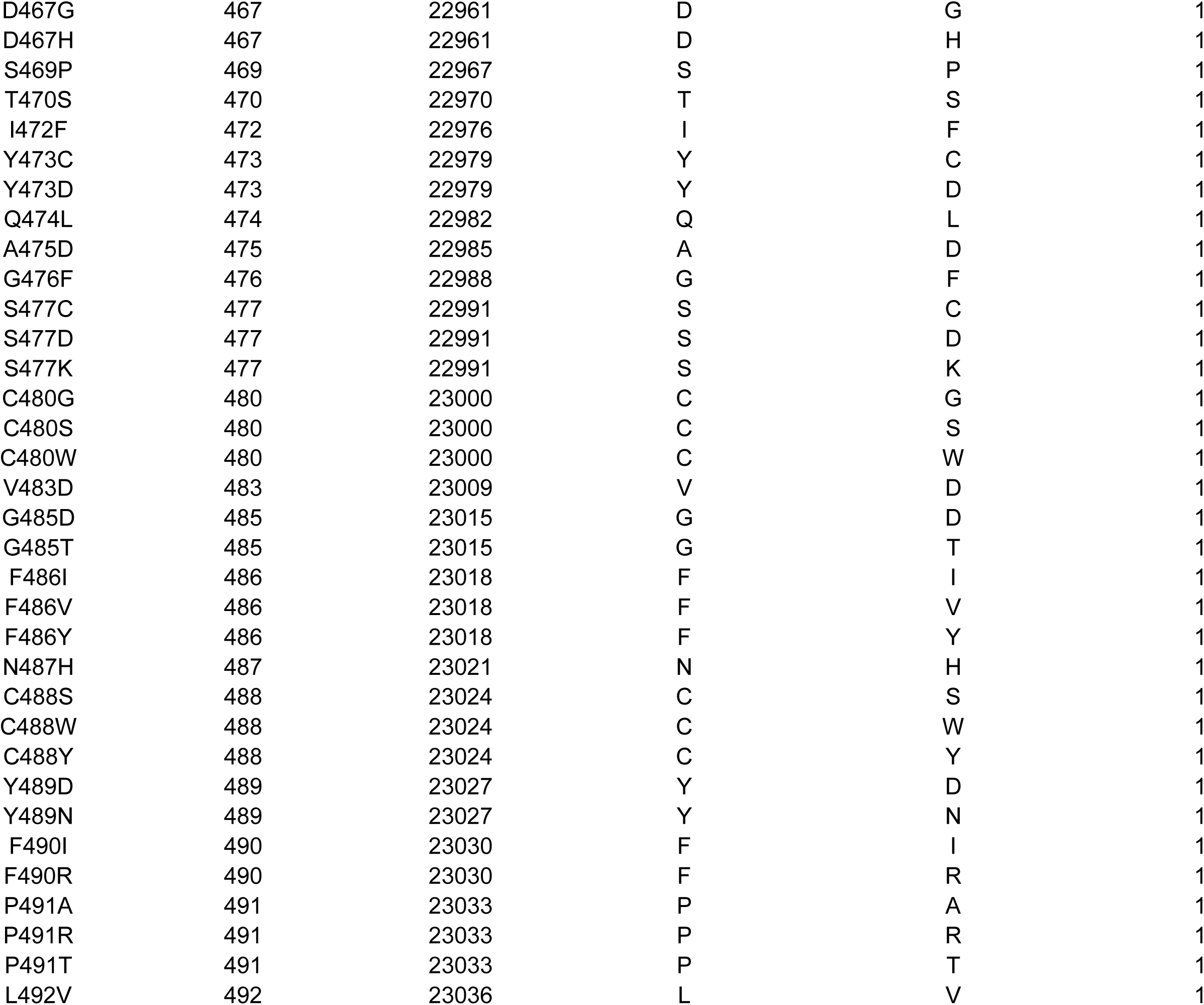
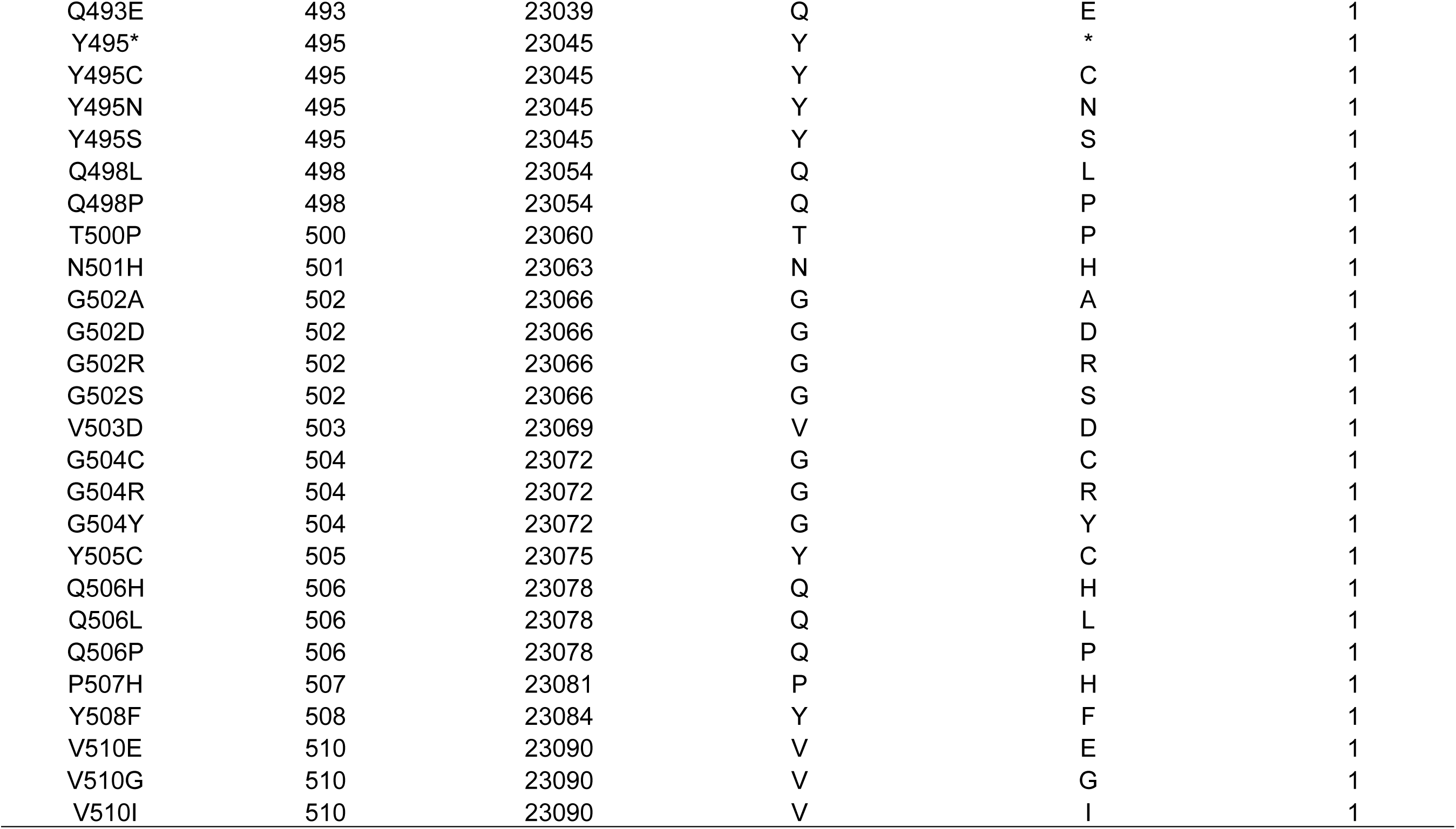
RBD missense mutations identified (http://cov-glue.cvr.gla.ac.uk/#/replacement, access July 31, 2021)

| Replacement | Codon Number | Ref Nt Position | Ref. Amino Acid | Replaced Amino Acid | Num seqs |
| --- | --- | --- | --- | --- | --- |
| N501Y | 501 | 23063 | N | Y | 257333 |
| S477N | 477 | 22991 | S | N | 36814 |
| L452R | 452 | 22916 | L | R | 22334 |
| N439K | 439 | 22877 | N | K | 21141 |
| E484K | 484 | 23012 | E | K | 16426 |
| K417N | 417 | 22811 | K | N | 6595 |
| T478K | 478 | 22994 | T | K | 6559 |
| S494P | 494 | 23042 | S | P | 2862 |
| N501T | 501 | 23063 | N | T | 2167 |
| K417T | 417 | 22811 | K | T | 1616 |
| V367F | 367 | 22661 | V | F | 1100 |
| Y453F | 453 | 22919 | Y | F | 981 |
| N440K | 440 | 22880 | N | K | 929 |
| P384L | 384 | 22712 | P | L | 896 |
| S477R | 477 | 22991 | S | R | 754 |
| F490S | 490 | 23030 | F | S | 612 |
| G446V | 446 | 22898 | G | V | 603 |
| T323I | 323 | 22529 | T | I | 585 |
| P330S | 330 | 22550 | P | S | 542 |
| V382L | 382 | 22706 | V | L | 493 |
| R357K | 357 | 22631 | R | K | 472 |
| P384S | 384 | 22712 | P | S | 379 |
| R346K | 346 | 22598 | R | K | 364 |
| P479S | 479 | 22997 | P | S | 361 |
| S477I | 477 | 22991 | S | I | 346 |
| E484Q | 484 | 23012 | E | Q | 300 |
| T478I | 478 | 22994 | T | I | 298 |
| S459F | 459 | 22937 | S | F | 292 |
| T385I | 385 | 22715 | T | I | 291 |
| L452Q | 452 | 22916 | L | Q | 268 |
| Q414K | 414 | 22802 | Q | K | 255 |
| Q414R | 414 | 22802 | Q | R | 232 |
| L455F | 455 | 22925 | L | F | 222 |
| A348S | 348 | 22604 | A | S | 219 |
| K444* | 444 | 22892 | K | * | 208 |
| Y508H | 508 | 23084 | Y | H | 207 |
| A475V | 475 | 22985 | A | V | 203 |
| T478R | 478 | 22994 | T | R | 202 |
| V483F | 483 | 23009 | V | F | 195 |
| N448Y | 448 | 22904 | N | Y | 191 |
| A344S | 344 | 22592 | A | S | 189 |
| N354D | 354 | 22622 | N | D | 183 |
| L452M | 452 | 22916 | L | M | 174 |
| N450K | 450 | 22910 | N | K | 170 |
| S373L | 373 | 22679 | S | L | 169 |
| F318L | 318 | 22514 | F | L | 167 |
| Q321E | 321 | 22523 | Q | E | 167 |
| A352S | 352 | 22616 | A | S | 164 |
| V320G | 320 | 22520 | V | G | 148 |
| G476S | 476 | 22988 | G | S | 147 |
| R408K | 408 | 22784 | R | K | 142 |
| F490L | 490 | 23030 | F | L | 136 |
| N481K | 481 | 23003 | N | K | 134 |
| T376I | 376 | 22688 | T | I | 128 |
| N354K | 354 | 22622 | N | K | 127 |
| E406Q | 406 | 22778 | E | Q | 122 |
| T385N | 385 | 22715 | T | N | 116 |
| S359N | 359 | 22637 | S | N | 106 |
| R403K | 403 | 22769 | R | K | 105 |
| K356R | 356 | 22628 | K | R | 104 |
| R408I | 408 | 22784 | R | I | 104 |
| I410V | 410 | 22790 | I | V | 104 |
| N370S | 370 | 22670 | N | S | 103 |
| F338L | 338 | 22574 | F | L | 102 |
| V483A | 483 | 23009 | V | A | 100 |
| V367L | 367 | 22661 | V | L | 94 |
| A411S | 411 | 22793 | A | S | 90 |
| A475S | 475 | 22985 | A | S | 90 |
| E471Q | 471 | 22973 | E | Q | 89 |
| V327I | 327 | 22541 | V | I | 84 |
| L335F | 335 | 22565 | L | F | 84 |
| G339D | 339 | 22577 | G | D | 84 |
| D427N | 427 | 22841 | D | N | 82 |
| K444N | 444 | 22892 | K | N | 76 |
| V362F | 362 | 22646 | V | F | 75 |
| T470I | 470 | 22970 | T | I | 74 |
| K444R | 444 | 22892 | K | R | 73 |
| G496S | 496 | 23048 | G | S | 73 |
| V341I | 341 | 22583 | V | I | 69 |
| S373P | 373 | 22679 | S | P | 69 |
| V367I | 367 | 22661 | V | I | 68 |
| P322S | 322 | 22526 | P | S | 66 |
| S459Y | 459 | 22937 | S | Y | 64 |
| A372V | 372 | 22676 | A | V | 63 |
| I434V | 434 | 22862 | I | V | 63 |
| I468V | 468 | 22964 | I | V | 63 |
| V320I | 320 | 22520 | V | I | 62 |
| S494L | 494 | 23042 | S | L | 61 |
| G446S | 446 | 22898 | G | S | 59 |
| V503I | 503 | 23069 | V | I | 59 |
| G404C | 404 | 22772 | G | C | 58 |
| G485V | 485 | 23015 | G | V | 58 |
| G339S | 339 | 22577 | G | S | 57 |
| A419S | 419 | 22817 | A | S | 57 |
| A348T | 348 | 22604 | A | T | 54 |
| T470N | 470 | 22970 | T | N | 54 |
| K458N | 458 | 22934 | K | N | 53 |
| P463S | 463 | 22949 | P | S | 52 |
| S373A | 373 | 22679 | S | A | 50 |
| Y508C | 508 | 23084 | Y | C | 50 |
| N394Y | 394 | 22742 | N | Y | 48 |
| I402V | 402 | 22766 | I | V | 48 |
| P412L | 412 | 22796 | P | L | 47 |
| R346S | 346 | 22598 | R | S | 46 |
| Q493K | 493 | 23039 | Q | K | 46 |
| K378N | 378 | 22694 | K | N | 45 |
| P330L | 330 | 22550 | P | L | 44 |
| Q493L | 493 | 23039 | Q | L | 44 |
| F486L | 486 | 23018 | F | L | 43 |
| V503F | 503 | 23069 | V | F | 43 |
| E324V | 324 | 22532 | E | V | 42 |
| G485R | 485 | 23015 | G | R | 42 |
| N370H | 370 | 22670 | N | H | 41 |
| T415N | 415 | 22805 | T | N | 41 |
| S477G | 477 | 22991 | S | G | 41 |
| P479L | 479 | 22997 | P | L | 41 |
| Q493R | 493 | 23039 | Q | R | 41 |
| E324Q | 324 | 22532 | E | Q | 40 |
| A352V | 352 | 22616 | A | V | 40 |
| F318S | 318 | 22514 | F | S | 39 |
| R346I | 346 | 22598 | R | I | 39 |
| N370K | 370 | 22670 | N | K | 39 |
| T430I | 430 | 22850 | T | I | 38 |
| V445A | 445 | 22895 | V | A | 38 |
| E484G | 484 | 23012 | E | G | 38 |
| Q321L | 321 | 22523 | Q | L | 36 |
| D427Y | 427 | 22841 | D | Y | 36 |
| E471D | 471 | 22973 | E | D | 36 |
| A348V | 348 | 22604 | A | V | 35 |
| N354S | 354 | 22622 | N | S | 35 |
| P384R | 384 | 22712 | P | R | 34 |
| L390F | 390 | 22730 | L | F | 34 |
| N448H | 448 | 22904 | N | H | 33 |
| V320F | 320 | 22520 | V | F | 32 |
| L368I | 368 | 22664 | L | I | 30 |
| V401L | 401 | 22763 | V | L | 30 |
| V407I | 407 | 22781 | V | I | 30 |
| I468T | 468 | 22964 | I | T | 30 |
| E484A | 484 | 23012 | E | A | 30 |
| P499S | 499 | 23057 | P | S | 30 |
| T415A | 415 | 22805 | T | A | 29 |
| G476A | 476 | 22988 | G | A | 29 |
| F490Y | 490 | 23030 | F | Y | 29 |
| Y505H | 505 | 23075 | Y | H | 29 |
| N331T | 331 | 22553 | N | T | 28 |
| D405Y | 405 | 22775 | D | Y | 28 |
| N501S | 501 | 23063 | N | S | 28 |
| N354H | 354 | 22622 | N | H | 27 |
| V367A | 367 | 22661 | V | A | 27 |
| D427G | 427 | 22841 | D | G | 27 |
| Y449H | 449 | 22907 | Y | H | 27 |
| Q321K | 321 | 22523 | Q | K | 26 |
| V433F | 433 | 22859 | V | F | 26 |
| Q493H | 493 | 23039 | Q | H | 26 |
| G482S | 482 | 23006 | G | S | 25 |
| P337S | 337 | 22571 | P | S | 24 |
| R346G | 346 | 22598 | R | G | 24 |
| E471K | 471 | 22973 | E | K | 24 |
| I472V | 472 | 22976 | I | V | 24 |
| V483I | 483 | 23009 | V | I | 24 |
| Q321* | 321 | 22523 | Q | * | 23 |
| S349P | 349 | 22607 | S | P | 23 |
| G485S | 485 | 23015 | G | S | 23 |
| Q506K | 506 | 23078 | Q | K | 23 |
| T323A | 323 | 22529 | T | A | 22 |
| A435S | 435 | 22865 | A | S | 22 |
| L441F | 441 | 22883 | L | F | 22 |
| E471G | 471 | 22973 | E | G | 22 |
| G485A | 485 | 23015 | G | A | 22 |
| G413W | 413 | 22799 | G | W | 21 |
| Q414* | 414 | 22802 | Q | * | 21 |
| N440Y | 440 | 22880 | N | Y | 21 |
| N448S | 448 | 22904 | N | S | 21 |
| G504S | 504 | 23072 | G | S | 21 |
| E340D | 340 | 22580 | E | D | 20 |
| N440T | 440 | 22880 | N | T | 20 |
| Y451H | 451 | 22913 | Y | H | 20 |
| A475T | 475 | 22985 | A | T | 20 |
| N394T | 394 | 22742 | N | T | 19 |
| V445I | 445 | 22895 | V | I | 19 |
| G447V | 447 | 22901 | G | V | 19 |
| T470A | 470 | 22970 | T | A | 19 |
| I472T | 472 | 22976 | I | T | 19 |
| N481D | 481 | 23003 | N | D | 19 |
| P499H | 499 | 23057 | P | H | 19 |
| N501I | 501 | 23063 | N | I | 19 |
| P507S | 507 | 23081 | P | S | 19 |
| V320A | 320 | 22520 | V | A | 18 |
| T323R | 323 | 22529 | T | R | 18 |
| A344T | 344 | 22592 | A | T | 18 |
| F329L | 329 | 22547 | F | L | 17 |
| A344V | 344 | 22592 | A | V | 17 |
| S366P | 366 | 22658 | S | P | 17 |
| D427H | 427 | 22841 | D | H | 17 |
| D427V | 427 | 22841 | D | V | 17 |
| P479H | 479 | 22997 | P | H | 17 |
| P491L | 491 | 23033 | P | L | 17 |
| Y505W | 505 | 23075 | Y | W | 17 |
| E324D | 324 | 22532 | E | D | 16 |
| S325F | 325 | 22535 | S | F | 16 |
| P337T | 337 | 22571 | P | T | 16 |
| T393I | 393 | 22739 | T | I | 16 |
| V445F | 445 | 22895 | V | F | 16 |
| F456L | 456 | 22928 | F | L | 16 |
| K462N | 462 | 22946 | K | N | 16 |
| T470K | 470 | 22970 | T | K | 16 |
| G504D | 504 | 23072 | G | D | 16 |
| G339V | 339 | 22577 | G | V | 15 |
| E340K | 340 | 22580 | E | K | 15 |
| D389E | 389 | 22727 | D | E | 15 |
| L441I | 441 | 22883 | L | I | 15 |
| N460I | 460 | 22940 | N | I | 15 |
| S371P | 371 | 22673 | S | P | 14 |
| W436C | 436 | 22868 | W | C | 14 |
| G447S | 447 | 22901 | G | S | 14 |
| R457S | 457 | 22931 | R | S | 14 |
| E465D | 465 | 22955 | E | D | 14 |
| E484D | 484 | 23012 | E | D | 14 |
| F490V | 490 | 23030 | F | V | 14 |
| G496V | 496 | 23048 | G | V | 14 |
| E324K | 324 | 22532 | E | K | 13 |
| E340G | 340 | 22580 | E | G | 13 |
| W353C | 353 | 22619 | W | C | 13 |
| S375F | 375 | 22685 | S | F | 13 |
| N450D | 450 | 22910 | N | D | 13 |
| Y489H | 489 | 23027 | Y | H | 13 |
| P499L | 499 | 23057 | P | L | 13 |
| V503A | 503 | 23069 | V | A | 13 |
| P507L | 507 | 23081 | P | L | 13 |
| V320L | 320 | 22520 | V | L | 12 |
| T333I | 333 | 22559 | T | I | 12 |
| N334K | 334 | 22562 | N | K | 12 |
| C361* | 361 | 22643 | C | * | 12 |
| C361T | 361 | 22643 | C | T | 12 |
| F374L | 374 | 22682 | F | L | 12 |
| E406D | 406 | 22778 | E | D | 12 |
| G413R | 413 | 22799 | G | R | 12 |
| R466K | 466 | 22958 | R | K | 12 |
| N481S | 481 | 23003 | N | S | 12 |
| Q493* | 493 | 23039 | Q | * | 12 |
| T345S | 345 | 22595 | T | S | 11 |
| K356N | 356 | 22628 | K | N | 11 |
| Y369H | 369 | 22667 | Y | H | 11 |
| K378R | 378 | 22694 | K | R | 11 |
| D405E | 405 | 22775 | D | E | 11 |
| G413E | 413 | 22799 | G | E | 11 |
| T415I | 415 | 22805 | T | I | 11 |
| V433I | 433 | 22859 | V | I | 11 |
| P463L | 463 | 22949 | P | L | 11 |
| S469L | 469 | 22967 | S | L | 11 |
| V483L | 483 | 23009 | V | L | 11 |
| P491S | 491 | 23033 | P | S | 11 |
| Y495H | 495 | 23045 | Y | H | 11 |
| P507A | 507 | 23081 | P | A | 11 |
| V510L | 510 | 23090 | V | L | 11 |
| P322A | 322 | 22526 | P | A | 10 |
| C336S | 336 | 22568 | C | S | 10 |
| P337L | 337 | 22571 | P | L | 10 |
| E340A | 340 | 22580 | E | A | 10 |
| A352D | 352 | 22616 | A | D | 10 |
| I358V | 358 | 22634 | I | V | 10 |
| F377L | 377 | 22691 | F | L | 10 |
| D428G | 428 | 22844 | D | G | 10 |
| S438F | 438 | 22874 | S | F | 10 |
| S443A | 443 | 22889 | S | A | 10 |
| N460K | 460 | 22940 | N | K | 10 |
| L461I | 461 | 22943 | L | I | 10 |
| L461P | 461 | 22943 | L | P | 10 |
| G476C | 476 | 22988 | G | C | 10 |
| S494A | 494 | 23042 | S | A | 10 |
| S359G | 359 | 22637 | S | G | 9 |
| S359T | 359 | 22637 | S | T | 9 |
| S371F | 371 | 22673 | S | F | 9 |
| N394S | 394 | 22742 | N | S | 9 |
| S399L | 399 | 22757 | S | L | 9 |
| I402L | 402 | 22766 | I | L | 9 |
| A411T | 411 | 22793 | A | T | 9 |
| A411V | 411 | 22793 | A | V | 9 |
| G413V | 413 | 22799 | G | V | 9 |
| N440D | 440 | 22880 | N | D | 9 |
| N448D | 448 | 22904 | N | D | 9 |
| N448T | 448 | 22904 | N | T | 9 |
| I472L | 472 | 22976 | I | L | 9 |
| G476V | 476 | 22988 | G | V | 9 |
| S477T | 477 | 22991 | S | T | 9 |
| Q506* | 506 | 23078 | Q | * | 9 |
| I326V | 326 | 22538 | I | V | 8 |
| V327F | 327 | 22541 | V | F | 8 |
| P337H | 337 | 22571 | P | H | 8 |
| R357I | 357 | 22631 | R | I | 8 |
| S359R | 359 | 22637 | S | R | 8 |
| D364N | 364 | 22652 | D | N | 8 |
| S371T | 371 | 22673 | S | T | 8 |
| A372S | 372 | 22676 | A | S | 8 |
| A372T | 372 | 22676 | A | T | 8 |
| T376S | 376 | 22688 | T | S | 8 |
| G381R | 381 | 22703 | G | R | 8 |
| D389N | 389 | 22727 | D | N | 8 |
| V395I | 395 | 22745 | V | I | 8 |
| V395L | 395 | 22745 | V | L | 8 |
| R408T | 408 | 22784 | R | T | 8 |
| Q409* | 409 | 22787 | Q | * | 8 |
| K417R | 417 | 22811 | K | R | 8 |
| I418V | 418 | 22814 | I | V | 8 |
| P426S | 426 | 22838 | P | S | 8 |
| N440S | 440 | 22880 | N | S | 8 |
| G446A | 446 | 22898 | G | A | 8 |
| Y449S | 449 | 22907 | Y | S | 8 |
| E465G | 465 | 22955 | E | G | 8 |
| D467Y | 467 | 22961 | D | Y | 8 |
| Q474H | 474 | 22982 | Q | H | 8 |
| T500I | 500 | 23060 | T | I | 8 |
| T500S | 500 | 23060 | T | S | 8 |
| R328I | 328 | 22544 | R | I | 7 |
| P330A | 330 | 22550 | P | A | 7 |
| N334H | 334 | 22562 | N | H | 7 |
| S349F | 349 | 22607 | S | F | 7 |
| K356M | 356 | 22628 | K | M | 7 |
| K356T | 356 | 22628 | K | T | 7 |
| I358T | 358 | 22634 | I | T | 7 |
| V382A | 382 | 22706 | V | A | 7 |
| V407A | 407 | 22781 | V | A | 7 |
| I434L | 434 | 22862 | I | L | 7 |
| G446D | 446 | 22898 | G | D | 7 |
| C480R | 480 | 23000 | C | R | 7 |
| E484* | 484 | 23012 | E | * | 7 |
| P322L | 322 | 22526 | P | L | 6 |
| G339C | 339 | 22577 | G | C | 6 |
| A344F | 344 | 22592 | A | F | 6 |
| R346T | 346 | 22598 | R | T | 6 |
| W353L | 353 | 22619 | W | L | 6 |
| S375P | 375 | 22685 | S | P | 6 |
| Y380F | 380 | 22700 | Y | F | 6 |
| S383F | 383 | 22709 | S | F | 6 |
| V395F | 395 | 22745 | V | F | 6 |
| Y396H | 396 | 22748 | Y | H | 6 |
| A397S | 397 | 22751 | A | S | 6 |
| D398N | 398 | 22754 | D | N | 6 |
| D405G | 405 | 22775 | D | G | 6 |
| R408G | 408 | 22784 | R | G | 6 |
| P412Q | 412 | 22796 | P | Q | 6 |
| Q414L | 414 | 22802 | Q | L | 6 |
| G416* | 416 | 22808 | G | * | 6 |
| D428Y | 428 | 22844 | D | Y | 6 |
| W436L | 436 | 22868 | W | L | 6 |
| K444M | 444 | 22892 | K | M | 6 |
| G447D | 447 | 22901 | G | D | 6 |
| R457M | 457 | 22931 | R | M | 6 |
| S459P | 459 | 22937 | S | P | 6 |
| N460S | 460 | 22940 | N | S | 6 |
| N460T | 460 | 22940 | N | T | 6 |
| K462I | 462 | 22946 | K | I | 6 |
| K462T | 462 | 22946 | K | T | 6 |
| F464L | 464 | 22952 | F | L | 6 |
| F464S | 464 | 22952 | F | S | 6 |
| E471A | 471 | 22973 | E | A | 6 |
| Y473F | 473 | 22979 | Y | F | 6 |
| Q474* | 474 | 22982 | Q | * | 6 |
| Q474E | 474 | 22982 | Q | E | 6 |
| Q474R | 474 | 22982 | Q | R | 6 |
| G496R | 496 | 23048 | G | R | 6 |
| P499R | 499 | 23057 | P | R | 6 |
| T500A | 500 | 23060 | T | A | 6 |
| F318Y | 318 | 22514 | F | Y | 5 |
| P322T | 322 | 22526 | P | T | 5 |
| S325Y | 325 | 22535 | S | Y | 5 |
| N331S | 331 | 22553 | N | S | 5 |
| I332V | 332 | 22556 | I | V | 5 |
| F342L | 342 | 22586 | F | L | 5 |
| N343Y | 343 | 22589 | N | Y | 5 |
| V362L | 362 | 22646 | V | L | 5 |
| Y365D | 365 | 22655 | Y | D | 5 |
| A372P | 372 | 22676 | A | P | 5 |
| S375T | 375 | 22685 | S | T | 5 |
| T376N | 376 | 22688 | T | N | 5 |
| K378* | 378 | 22694 | K | * | 5 |
| C379F | 379 | 22697 | C | F | 5 |
| V382E | 382 | 22706 | V | E | 5 |
| T385P | 385 | 22715 | T | P | 5 |
| D389Y | 389 | 22727 | D | Y | 5 |
| L390I | 390 | 22730 | L | I | 5 |
| L390P | 390 | 22730 | L | P | 5 |
| Y396F | 396 | 22748 | Y | F | 5 |
| S399P | 399 | 22757 | S | P | 5 |
| R403S | 403 | 22769 | R | S | 5 |
| G404V | 404 | 22772 | G | V | 5 |
| I410T | 410 | 22790 | I | T | 5 |
| G416E | 416 | 22808 | G | E | 5 |
| D420N | 420 | 22820 | D | N | 5 |
| D420Y | 420 | 22820 | D | Y | 5 |
| G431V | 431 | 22853 | G | V | 5 |
| A435T | 435 | 22865 | A | T | 5 |
| G447C | 447 | 22901 | G | C | 5 |
| N450S | 450 | 22910 | N | S | 5 |
| Y453H | 453 | 22919 | Y | H | 5 |
| R457T | 457 | 22931 | R | T | 5 |
| K458E | 458 | 22934 | K | E | 5 |
| K458R | 458 | 22934 | K | R | 5 |
| E465* | 465 | 22955 | E | * | 5 |
| S469* | 469 | 22967 | S | * | 5 |
| T478A | 478 | 22994 | T | A | 5 |
| G482V | 482 | 23006 | G | V | 5 |
| E484R | 484 | 23012 | E | R | 5 |
| G485C | 485 | 23015 | G | C | 5 |
| N487D | 487 | 23021 | N | D | 5 |
| C488F | 488 | 23024 | C | F | 5 |
| C488R | 488 | 23024 | C | R | 5 |
| Y489* | 489 | 23027 | Y | * | 5 |
| Q498H | 498 | 23054 | Q | H | 5 |
| N501K | 501 | 23063 | N | K | 5 |
| P507Q | 507 | 23081 | P | Q | 5 |
| E324A | 324 | 22532 | E | A | 4 |
| S325A | 325 | 22535 | S | A | 4 |
| R328G | 328 | 22544 | R | G | 4 |
| R328S | 328 | 22544 | R | S | 4 |
| T333K | 333 | 22559 | T | K | 4 |
| F338S | 338 | 22574 | F | S | 4 |
| A344D | 344 | 22592 | A | D | 4 |
| T345F | 345 | 22595 | T | F | 4 |
| R346F | 346 | 22598 | R | F | 4 |
| F347L | 347 | 22601 | F | L | 4 |
| A348P | 348 | 22604 | A | P | 4 |
| W353R | 353 | 22619 | W | R | 4 |
| R355S | 355 | 22625 | R | S | 4 |
| N360S | 360 | 22640 | N | S | 4 |
| C361F | 361 | 22643 | C | F | 4 |
| C361S | 361 | 22643 | C | S | 4 |
| A363V | 363 | 22649 | A | V | 4 |
| D364Y | 364 | 22652 | D | Y | 4 |
| F374S | 374 | 22682 | F | S | 4 |
| F374Y | 374 | 22682 | F | Y | 4 |
| S375Y | 375 | 22685 | S | Y | 4 |
| K378M | 378 | 22694 | K | M | 4 |
| S383P | 383 | 22709 | S | P | 4 |
| N394I | 394 | 22742 | N | I | 4 |
| V395A | 395 | 22745 | V | A | 4 |
| R408S | 408 | 22784 | R | S | 4 |
| A411D | 411 | 22793 | A | D | 4 |
| G413A | 413 | 22799 | G | A | 4 |
| T415S | 415 | 22805 | T | S | 4 |
| A419T | 419 | 22817 | A | T | 4 |
| L425S | 425 | 22835 | L | S | 4 |
| P426L | 426 | 22838 | P | L | 4 |
| D427A | 427 | 22841 | D | A | 4 |
| T430A | 430 | 22850 | T | A | 4 |
| W436* | 436 | 22868 | W | * | 4 |
| N437S | 437 | 22871 | N | S | 4 |
| S438P | 438 | 22874 | S | P | 4 |
| D442Y | 442 | 22886 | D | Y | 4 |
| G446C | 446 | 22898 | G | C | 4 |
| Y449C | 449 | 22907 | Y | C | 4 |
| R454* | 454 | 22922 | R | * | 4 |
| L455S | 455 | 22925 | L | S | 4 |
| R457K | 457 | 22931 | R | K | 4 |
| K458Q | 458 | 22934 | K | Q | 4 |
| K462E | 462 | 22946 | K | E | 4 |
| K462R | 462 | 22946 | K | R | 4 |
| P463H | 463 | 22949 | P | H | 4 |
| I468F | 468 | 22964 | I | F | 4 |
| N481T | 481 | 23003 | N | T | 4 |
| G482A | 482 | 23006 | G | A | 4 |
| V483G | 483 | 23009 | V | G | 4 |
| E484V | 484 | 23012 | E | V | 4 |
| F486S | 486 | 23018 | F | S | 4 |
| N487Y | 487 | 23021 | N | Y | 4 |
| Y489C | 489 | 23027 | Y | C | 4 |
| Q498K | 498 | 23054 | Q | K | 4 |
| Q498R | 498 | 23054 | Q | R | 4 |
| P499T | 499 | 23057 | P | T | 4 |
| G502C | 502 | 23066 | G | C | 4 |
| R509I | 509 | 23087 | R | I | 4 |
| R509S | 509 | 23087 | R | S | 4 |
| R319I | 319 | 22517 | R | I | 3 |
| R319S | 319 | 22517 | R | S | 3 |
| R319T | 319 | 22517 | R | T | 3 |
| T323S | 323 | 22529 | T | S | 3 |
| R328K | 328 | 22544 | R | K | 3 |
| F329S | 329 | 22547 | F | S | 3 |
| C336Y | 336 | 22568 | C | Y | 3 |
| E340* | 340 | 22580 | E | * | 3 |
| T345A | 345 | 22595 | T | A | 3 |
| T345I | 345 | 22595 | T | I | 3 |
| S349A | 349 | 22607 | S | A | 3 |
| Y351H | 351 | 22613 | Y | H | 3 |
| R355T | 355 | 22625 | R | T | 3 |
| K356E | 356 | 22628 | K | E | 3 |
| I358L | 358 | 22634 | I | L | 3 |
| C361R | 361 | 22643 | C | R | 3 |
| A363P | 363 | 22649 | A | P | 3 |
| Y365H | 365 | 22655 | Y | H | 3 |
| Y365S | 365 | 22655 | Y | S | 3 |
| S366Y | 366 | 22658 | S | Y | 3 |
| S371A | 371 | 22673 | S | A | 3 |
| S373* | 373 | 22679 | S | * | 3 |
| C379S | 379 | 22697 | C | S | 3 |
| T385A | 385 | 22715 | T | A | 3 |
| K386R | 386 | 22718 | K | R | 3 |
| N388S | 388 | 22724 | N | S | 3 |
| C391* | 391 | 22733 | C | * | 3 |
| F392S | 392 | 22736 | F | S | 3 |
| T393A | 393 | 22739 | T | A | 3 |
| Y396C | 396 | 22748 | Y | C | 3 |
| Y396N | 396 | 22748 | Y | N | 3 |
| A397V | 397 | 22751 | A | V | 3 |
| S399* | 399 | 22757 | S | * | 3 |
| F400L | 400 | 22760 | F | L | 3 |
| R403G | 403 | 22769 | R | G | 3 |
| R403I | 403 | 22769 | R | I | 3 |
| G404S | 404 | 22772 | G | S | 3 |
| D405N | 405 | 22775 | D | N | 3 |
| E406* | 406 | 22778 | E | * | 3 |
| D420A | 420 | 22820 | D | A | 3 |
| D420G | 420 | 22820 | D | G | 3 |
| N422I | 422 | 22826 | N | I | 3 |
| L425F | 425 | 22835 | L | F | 3 |
| L425V | 425 | 22835 | L | V | 3 |
| P426Q | 426 | 22838 | P | Q | 3 |
| D428E | 428 | 22844 | D | E | 3 |
| T430S | 430 | 22850 | T | S | 3 |
| G431C | 431 | 22853 | G | C | 3 |
| I434M | 434 | 22862 | I | M | 3 |
| A435V | 435 | 22865 | A | V | 3 |
| D442V | 442 | 22886 | D | V | 3 |
| N448K | 448 | 22904 | N | K | 3 |
| Y451F | 451 | 22913 | Y | F | 3 |
| L452V | 452 | 22916 | L | V | 3 |
| Y453C | 453 | 22919 | Y | C | 3 |
| L455V | 455 | 22925 | L | V | 3 |
| K458* | 458 | 22934 | K | * | 3 |
| K458T | 458 | 22934 | K | T | 3 |
| L461F | 461 | 22943 | L | F | 3 |
| D467E | 467 | 22961 | D | E | 3 |
| D467V | 467 | 22961 | D | V | 3 |
| I468M | 468 | 22964 | I | M | 3 |
| S469A | 469 | 22967 | S | A | 3 |
| S469T | 469 | 22967 | S | T | 3 |
| E471* | 471 | 22973 | E | * | 3 |
| E471V | 471 | 22973 | E | V | 3 |
| Y473* | 473 | 22979 | Y | * | 3 |
| A475P | 475 | 22985 | A | P | 3 |
| G476D | 476 | 22988 | G | D | 3 |
| T478S | 478 | 22994 | T | S | 3 |
| C480* | 480 | 23000 | C | * | 3 |
| N481H | 481 | 23003 | N | H | 3 |
| N481I | 481 | 23003 | N | I | 3 |
| N481Y | 481 | 23003 | N | Y | 3 |
| G482C | 482 | 23006 | G | C | 3 |
| G482R | 482 | 23006 | G | R | 3 |
| N487I | 487 | 23021 | N | I | 3 |
| N487K | 487 | 23021 | N | K | 3 |
| C488* | 488 | 23024 | C | * | 3 |
| S494* | 494 | 23042 | S | * | 3 |
| S494R | 494 | 23042 | S | R | 3 |
| S494T | 494 | 23042 | S | T | 3 |
| G496C | 496 | 23048 | G | C | 3 |
| Q498* | 498 | 23054 | Q | * | 3 |
| T500N | 500 | 23060 | T | N | 3 |
| G502V | 502 | 23066 | G | V | 3 |
| G504N | 504 | 23072 | G | N | 3 |
| G504V | 504 | 23072 | G | V | 3 |
| Y505* | 505 | 23075 | Y | * | 3 |
| V510A | 510 | 23090 | V | A | 3 |
| F318C | 318 | 22514 | F | C | 2 |
| F318I | 318 | 22514 | F | I | 2 |
| F318V | 318 | 22514 | F | V | 2 |
| R319K | 319 | 22517 | R | K | 2 |
| Q321H | 321 | 22523 | Q | H | 2 |
| Q321R | 321 | 22523 | Q | R | 2 |
| E324G | 324 | 22532 | E | G | 2 |
| V327A | 327 | 22541 | V | A | 2 |
| V327D | 327 | 22541 | V | D | 2 |
| P330T | 330 | 22550 | P | T | 2 |
| N331D | 331 | 22553 | N | D | 2 |
| T333A | 333 | 22559 | T | A | 2 |
| N334I | 334 | 22562 | N | I | 2 |
| N334Y | 334 | 22562 | N | Y | 2 |
| L335S | 335 | 22565 | L | S | 2 |
| F338V | 338 | 22574 | F | V | 2 |
| G339R | 339 | 22577 | G | R | 2 |
| E340V | 340 | 22580 | E | V | 2 |
| V341F | 341 | 22583 | V | F | 2 |
| N343D | 343 | 22589 | N | D | 2 |
| A344P | 344 | 22592 | A | P | 2 |
| F347C | 347 | 22601 | F | C | 2 |
| F347Y | 347 | 22601 | F | Y | 2 |
| A348E | 348 | 22604 | A | E | 2 |
| A348F | 348 | 22604 | A | F | 2 |
| A348G | 348 | 22604 | A | G | 2 |
| V350A | 350 | 22610 | V | A | 2 |
| V350F | 350 | 22610 | V | F | 2 |
| Y351C | 351 | 22613 | Y | C | 2 |
| Y351F | 351 | 22613 | Y | F | 2 |
| A352T | 352 | 22616 | A | T | 2 |
| W353S | 353 | 22619 | W | S | 2 |
| I358F | 358 | 22634 | I | F | 2 |
| V362I | 362 | 22646 | V | I | 2 |
| S366F | 366 | 22658 | S | F | 2 |
| S366L | 366 | 22658 | S | L | 2 |
| Y369C | 369 | 22667 | Y | C | 2 |
| Y369D | 369 | 22667 | Y | D | 2 |
| N370D | 370 | 22670 | N | D | 2 |
| A372L | 372 | 22676 | A | L | 2 |
| F374V | 374 | 22682 | F | V | 2 |
| F377V | 377 | 22691 | F | V | 2 |
| K378E | 378 | 22694 | K | E | 2 |
| G381V | 381 | 22703 | G | V | 2 |
| V382M | 382 | 22706 | V | M | 2 |
| S383T | 383 | 22709 | S | T | 2 |
| P384A | 384 | 22712 | P | A | 2 |
| P384H | 384 | 22712 | P | H | 2 |
| P384T | 384 | 22712 | P | T | 2 |
| L387F | 387 | 22721 | L | F | 2 |
| N388Y | 388 | 22724 | N | Y | 2 |
| D389G | 389 | 22727 | D | G | 2 |
| D389H | 389 | 22727 | D | H | 2 |
| C391F | 391 | 22733 | C | F | 2 |
| C391S | 391 | 22733 | C | S | 2 |
| F392I | 392 | 22736 | F | I | 2 |
| N394K | 394 | 22742 | N | K | 2 |
| D398V | 398 | 22754 | D | V | 2 |
| F400S | 400 | 22760 | F | S | 2 |
| V401A | 401 | 22763 | V | A | 2 |
| V401I | 401 | 22763 | V | I | 2 |
| I402F | 402 | 22766 | I | F | 2 |
| I402T | 402 | 22766 | I | T | 2 |
| E406G | 406 | 22778 | E | G | 2 |
| E406K | 406 | 22778 | E | K | 2 |
| V407F | 407 | 22781 | V | F | 2 |
| Q409H | 409 | 22787 | Q | H | 2 |
| Q409K | 409 | 22787 | Q | K | 2 |
| I410L | 410 | 22790 | I | L | 2 |
| A411P | 411 | 22793 | A | P | 2 |
| P412T | 412 | 22796 | P | T | 2 |
| Q414H | 414 | 22802 | Q | H | 2 |
| Q414P | 414 | 22802 | Q | P | 2 |
| K417E | 417 | 22811 | K | E | 2 |
| I418T | 418 | 22814 | I | T | 2 |
| A419V | 419 | 22817 | A | V | 2 |
| N422S | 422 | 22826 | N | S | 2 |
| Y423H | 423 | 22829 | Y | H | 2 |
| K424N | 424 | 22832 | K | N | 2 |
| P426T | 426 | 22838 | P | T | 2 |
| D428V | 428 | 22844 | D | V | 2 |
| F429L | 429 | 22847 | F | L | 2 |
| G431D | 431 | 22853 | G | D | 2 |
| C432* | 432 | 22856 | C | * | 2 |
| C432F | 432 | 22856 | C | F | 2 |
| I434T | 434 | 22862 | I | T | 2 |
| A435D | 435 | 22865 | A | D | 2 |
| A435P | 435 | 22865 | A | P | 2 |
| W436R | 436 | 22868 | W | R | 2 |
| S438A | 438 | 22874 | S | A | 2 |
| S438T | 438 | 22874 | S | T | 2 |
| N439S | 439 | 22877 | N | S | 2 |
| L441R | 441 | 22883 | L | R | 2 |
| S443F | 443 | 22889 | S | F | 2 |
| S443P | 443 | 22889 | S | P | 2 |
| Y449F | 449 | 22907 | Y | F | 2 |
| Y449N | 449 | 22907 | Y | N | 2 |
| Y451* | 451 | 22913 | Y | * | 2 |
| Y451C | 451 | 22913 | Y | C | 2 |
| Y451D | 451 | 22913 | Y | D | 2 |
| L452P | 452 | 22916 | L | P | 2 |
| Y453* | 453 | 22919 | Y | * | 2 |
| R454G | 454 | 22922 | R | G | 2 |
| R454S | 454 | 22922 | R | S | 2 |
| R454T | 454 | 22922 | R | T | 2 |
| F456S | 456 | 22928 | F | S | 2 |
| F456Y | 456 | 22928 | F | Y | 2 |
| R457W | 457 | 22931 | R | W | 2 |
| N460H | 460 | 22940 | N | H | 2 |
| P463T | 463 | 22949 | P | T | 2 |
| F464I | 464 | 22952 | F | I | 2 |
| F464V | 464 | 22952 | F | V | 2 |
| E465V | 465 | 22955 | E | V | 2 |
| R466G | 466 | 22958 | R | G | 2 |
| R466I | 466 | 22958 | R | I | 2 |
| R466S | 466 | 22958 | R | S | 2 |
| I468L | 468 | 22964 | I | L | 2 |
| I468N | 468 | 22964 | I | N | 2 |
| Y473H | 473 | 22979 | Y | H | 2 |
| Q474K | 474 | 22982 | Q | K | 2 |
| A475G | 475 | 22985 | A | G | 2 |
| P479T | 479 | 22997 | P | T | 2 |
| C480F | 480 | 23000 | C | F | 2 |
| N487S | 487 | 23021 | N | S | 2 |
| N487T | 487 | 23021 | N | T | 2 |
| Y489F | 489 | 23027 | Y | F | 2 |
| Y489S | 489 | 23027 | Y | S | 2 |
| F490C | 490 | 23030 | F | C | 2 |
| P491H | 491 | 23033 | P | H | 2 |
| L492F | 492 | 23036 | L | F | 2 |
| Q493P | 493 | 23039 | Q | P | 2 |
| Y495F | 495 | 23045 | Y | F | 2 |
| F497L | 497 | 23051 | F | L | 2 |
| F497S | 497 | 23051 | F | S | 2 |
| F497V | 497 | 23051 | F | V | 2 |
| V503L | 503 | 23069 | V | L | 2 |
| Y505F | 505 | 23075 | Y | F | 2 |
| Y505N | 505 | 23075 | Y | N | 2 |
| Q506R | 506 | 23078 | Q | R | 2 |
| Y508N | 508 | 23084 | Y | N | 2 |
| R509K | 509 | 23087 | R | K | 2 |
| R509T | 509 | 23087 | R | T | 2 |
| F318* | 318 | 22514 | F | * | 1 |
| F318N | 318 | 22514 | F | N | 1 |
| R319* | 319 | 22517 | R | * | 1 |
| R319G | 319 | 22517 | R | G | 1 |
| V320D | 320 | 22520 | V | D | 1 |
| P322Q | 322 | 22526 | P | Q | 1 |
| T323M | 323 | 22529 | T | M | 1 |
| T323P | 323 | 22529 | T | P | 1 |
| E324L | 324 | 22532 | E | L | 1 |
| S325C | 325 | 22535 | S | C | 1 |
| S325K | 325 | 22535 | S | K | 1 |
| S325L | 325 | 22535 | S | L | 1 |
| S325P | 325 | 22535 | S | P | 1 |
| I326M | 326 | 22538 | I | M | 1 |
| I326N | 326 | 22538 | I | N | 1 |
| I326S | 326 | 22538 | I | S | 1 |
| V327L | 327 | 22541 | V | L | 1 |
| R328T | 328 | 22544 | R | T | 1 |
| F329I | 329 | 22547 | F | I | 1 |
| P330H | 330 | 22550 | P | H | 1 |
| P330R | 330 | 22550 | P | R | 1 |
| N331I | 331 | 22553 | N | I | 1 |
| I332T | 332 | 22556 | I | T | 1 |
| N334D | 334 | 22562 | N | D | 1 |
| L335M | 335 | 22565 | L | M | 1 |
| C336F | 336 | 22568 | C | F | 1 |
| C336R | 336 | 22568 | C | R | 1 |
| F338P | 338 | 22574 | F | P | 1 |
| F338Y | 338 | 22574 | F | Y | 1 |
| E340Q | 340 | 22580 | E | Q | 1 |
| V341A | 341 | 22583 | V | A | 1 |
| V341G | 341 | 22583 | V | G | 1 |
| V341S | 341 | 22583 | V | S | 1 |
| F342C | 342 | 22586 | F | C | 1 |
| N343K | 343 | 22589 | N | K | 1 |
| N343S | 343 | 22589 | N | S | 1 |
| T345N | 345 | 22595 | T | N | 1 |
| T345P | 345 | 22595 | T | P | 1 |
| F347I | 347 | 22601 | F | I | 1 |
| F347S | 347 | 22601 | F | S | 1 |
| F347V | 347 | 22601 | F | V | 1 |
| A348L | 348 | 22604 | A | L | 1 |
| A348Q | 348 | 22604 | A | Q | 1 |
| S349Y | 349 | 22607 | S | Y | 1 |
| V350Q | 350 | 22610 | V | Q | 1 |
| Y351N | 351 | 22613 | Y | N | 1 |
| Y351S | 351 | 22613 | Y | S | 1 |
| A352G | 352 | 22616 | A | G | 1 |
| A352P | 352 | 22616 | A | P | 1 |
| W353* | 353 | 22619 | W | * | 1 |
| W353F | 353 | 22619 | W | F | 1 |
| N354Y | 354 | 22622 | N | Y | 1 |
| R355G | 355 | 22625 | R | G | 1 |
| R355K | 355 | 22625 | R | K | 1 |
| R355M | 355 | 22625 | R | M | 1 |
| K356Y | 356 | 22628 | K | Y | 1 |
| R357G | 357 | 22631 | R | G | 1 |
| I358A | 358 | 22634 | I | A | 1 |
| V362D | 362 | 22646 | V | D | 1 |
| V362G | 362 | 22646 | V | G | 1 |
| A363S | 363 | 22649 | A | S | 1 |
| A363T | 363 | 22649 | A | T | 1 |
| D364A | 364 | 22652 | D | A | 1 |
| D364G | 364 | 22652 | D | G | 1 |
| Y365C | 365 | 22655 | Y | C | 1 |
| L368R | 368 | 22664 | L | R | 1 |
| Y369F | 369 | 22667 | Y | F | 1 |
| A372E | 372 | 22676 | A | E | 1 |
| S373T | 373 | 22679 | S | T | 1 |
| F374C | 374 | 22682 | F | C | 1 |
| S375A | 375 | 22685 | S | A | 1 |
| S375C | 375 | 22685 | S | C | 1 |
| T376L | 376 | 22688 | T | L | 1 |
| F377S | 377 | 22691 | F | S | 1 |
| C379W | 379 | 22697 | C | W | 1 |
| Y380* | 380 | 22700 | Y | * | 1 |
| Y380C | 380 | 22700 | Y | C | 1 |
| Y380H | 380 | 22700 | Y | H | 1 |
| G381* | 381 | 22703 | G | * | 1 |
| G381E | 381 | 22703 | G | E | 1 |
| G381L | 381 | 22703 | G | L | 1 |
| V382G | 382 | 22706 | V | G | 1 |
| S383A | 383 | 22709 | S | A | 1 |
| T385S | 385 | 22715 | T | S | 1 |
| K386I | 386 | 22718 | K | I | 1 |
| L387* | 387 | 22721 | L | * | 1 |
| N388D | 388 | 22724 | N | D | 1 |
| N388H | 388 | 22724 | N | H | 1 |
| N388I | 388 | 22724 | N | I | 1 |
| N388T | 388 | 22724 | N | T | 1 |
| D389T | 389 | 22727 | D | T | 1 |
| L390H | 390 | 22730 | L | H | 1 |
| L390V | 390 | 22730 | L | V | 1 |
| C391R | 391 | 22733 | C | R | 1 |
| F392C | 392 | 22736 | F | C | 1 |
| F392L | 392 | 22736 | F | L | 1 |
| F392P | 392 | 22736 | F | P | 1 |
| T393P | 393 | 22739 | T | P | 1 |
| Y396D | 396 | 22748 | Y | D | 1 |
| A397E | 397 | 22751 | A | E | 1 |
| A397P | 397 | 22751 | A | P | 1 |
| A397T | 397 | 22751 | A | T | 1 |
| D398G | 398 | 22754 | D | G | 1 |
| D398Y | 398 | 22754 | D | Y | 1 |
| S399A | 399 | 22757 | S | A | 1 |
| S399T | 399 | 22757 | S | T | 1 |
| I402S | 402 | 22766 | I | S | 1 |
| R403* | 403 | 22769 | R | * | 1 |
| G404A | 404 | 22772 | G | A | 1 |
| G404D | 404 | 22772 | G | D | 1 |
| G404R | 404 | 22772 | G | R | 1 |
| D405H | 405 | 22775 | D | H | 1 |
| V407D | 407 | 22781 | V | D | 1 |
| V407R | 407 | 22781 | V | R | 1 |
| V407S | 407 | 22781 | V | S | 1 |
| R408* | 408 | 22784 | R | * | 1 |
| Q409E | 409 | 22787 | Q | E | 1 |
| Q409L | 409 | 22787 | Q | L | 1 |
| A411G | 411 | 22793 | A | G | 1 |
| P412A | 412 | 22796 | P | A | 1 |
| P412S | 412 | 22796 | P | S | 1 |
| G413C | 413 | 22799 | G | C | 1 |
| T415P | 415 | 22805 | T | P | 1 |
| G416R | 416 | 22808 | G | R | 1 |
| G416V | 416 | 22808 | G | V | 1 |
| K417D | 417 | 22811 | K | D | 1 |
| K417M | 417 | 22811 | K | M | 1 |
| I418F | 418 | 22814 | I | F | 1 |
| A419D | 419 | 22817 | A | D | 1 |
| A419G | 419 | 22817 | A | G | 1 |
| Y421* | 421 | 22823 | Y | * | 1 |
| Y421F | 421 | 22823 | Y | F | 1 |
| Y423* | 423 | 22829 | Y | * | 1 |
| Y423F | 423 | 22829 | Y | F | 1 |
| Y423S | 423 | 22829 | Y | S | 1 |
| K424* | 424 | 22832 | K | * | 1 |
| K424I | 424 | 22832 | K | I | 1 |
| K424Q | 424 | 22832 | K | Q | 1 |
| D427E | 427 | 22841 | D | E | 1 |
| T430K | 430 | 22850 | T | K | 1 |
| T430N | 430 | 22850 | T | N | 1 |
| T430P | 430 | 22850 | T | P | 1 |
| G431R | 431 | 22853 | G | R | 1 |
| G431S | 431 | 22853 | G | S | 1 |
| V433D | 433 | 22859 | V | D | 1 |
| V433L | 433 | 22859 | V | L | 1 |
| I434K | 434 | 22862 | I | K | 1 |
| W436S | 436 | 22868 | W | S | 1 |
| N437D | 437 | 22871 | N | D | 1 |
| N437I | 437 | 22871 | N | I | 1 |
| S438Y | 438 | 22874 | S | Y | 1 |
| N439* | 439 | 22877 | N | * | 1 |
| N439D | 439 | 22877 | N | D | 1 |
| N440H | 440 | 22880 | N | H | 1 |
| L441V | 441 | 22883 | L | V | 1 |
| D442A | 442 | 22886 | D | A | 1 |
| V445D | 445 | 22895 | V | D | 1 |
| V445G | 445 | 22895 | V | G | 1 |
| G446R | 446 | 22898 | G | R | 1 |
| G447A | 447 | 22901 | G | A | 1 |
| G447R | 447 | 22901 | G | R | 1 |
| Y449* | 449 | 22907 | Y | * | 1 |
| N450H | 450 | 22910 | N | H | 1 |
| N450I | 450 | 22910 | N | I | 1 |
| L452F | 452 | 22916 | L | F | 1 |
| Y453S | 453 | 22919 | Y | S | 1 |
| R454I | 454 | 22922 | R | I | 1 |
| R454K | 454 | 22922 | R | K | 1 |
| F456* | 456 | 22928 | F | * | 1 |
| F456I | 456 | 22928 | F | I | 1 |
| F456V | 456 | 22928 | F | V | 1 |
| S459C | 459 | 22937 | S | C | 1 |
| N460D | 460 | 22940 | N | D | 1 |
| N460Y | 460 | 22940 | N | Y | 1 |
| K462* | 462 | 22946 | K | * | 1 |
| P463A | 463 | 22949 | P | A | 1 |
| F464C | 464 | 22952 | F | C | 1 |
| E465K | 465 | 22955 | E | K | 1 |
| D467G | 467 | 22961 | D | G | 1 |
| D467H | 467 | 22961 | D | H | 1 |
| S469P | 469 | 22967 | S | P | 1 |
| T470S | 470 | 22970 | T | S | 1 |
| I472F | 472 | 22976 | I | F | 1 |
| Y473C | 473 | 22979 | Y | C | 1 |
| Y473D | 473 | 22979 | Y | D | 1 |
| Q474L | 474 | 22982 | Q | L | 1 |
| A475D | 475 | 22985 | A | D | 1 |
| G476F | 476 | 22988 | G | F | 1 |
| S477C | 477 | 22991 | S | C | 1 |
| S477D | 477 | 22991 | S | D | 1 |
| S477K | 477 | 22991 | S | K | 1 |
| C480G | 480 | 23000 | C | G | 1 |
| C480S | 480 | 23000 | C | S | 1 |
| C480W | 480 | 23000 | C | W | 1 |
| V483D | 483 | 23009 | V | D | 1 |
| G485D | 485 | 23015 | G | D | 1 |
| G485T | 485 | 23015 | G | T | 1 |
| F486I | 486 | 23018 | F | I | 1 |
| F486V | 486 | 23018 | F | V | 1 |
| F486Y | 486 | 23018 | F | Y | 1 |
| N487H | 487 | 23021 | N | H | 1 |
| C488S | 488 | 23024 | C | S | 1 |
| C488W | 488 | 23024 | C | W | 1 |
| C488Y | 488 | 23024 | C | Y | 1 |
| Y489D | 489 | 23027 | Y | D | 1 |
| Y489N | 489 | 23027 | Y | N | 1 |
| F490I | 490 | 23030 | F | I | 1 |
| F490R | 490 | 23030 | F | R | 1 |
| P491A | 491 | 23033 | P | A | 1 |
| P491R | 491 | 23033 | P | R | 1 |
| P491T | 491 | 23033 | P | T | 1 |
| L492V | 492 | 23036 | L | V | 1 |
| Q493E | 493 | 23039 | Q | E | 1 |
| Y495* | 495 | 23045 | Y | * | 1 |
| Y495C | 495 | 23045 | Y | C | 1 |
| Y495N | 495 | 23045 | Y | N | 1 |
| Y495S | 495 | 23045 | Y | S | 1 |
| Q498L | 498 | 23054 | Q | L | 1 |
| Q498P | 498 | 23054 | Q | P | 1 |
| T500P | 500 | 23060 | T | P | 1 |
| N501H | 501 | 23063 | N | H | 1 |
| G502A | 502 | 23066 | G | A | 1 |
| G502D | 502 | 23066 | G | D | 1 |
| G502R | 502 | 23066 | G | R | 1 |
| G502S | 502 | 23066 | G | S | 1 |
| V503D | 503 | 23069 | V | D | 1 |
| G504C | 504 | 23072 | G | C | 1 |
| G504R | 504 | 23072 | G | R | 1 |
| G504Y | 504 | 23072 | G | Y | 1 |
| Y505C | 505 | 23075 | Y | C | 1 |
| Q506H | 506 | 23078 | Q | H | 1 |
| Q506L | 506 | 23078 | Q | L | 1 |
| Q506P | 506 | 23078 | Q | P | 1 |
| P507H | 507 | 23081 | P | H | 1 |
| Y508F | 508 | 23084 | Y | F | 1 |
| V510E | 510 | 23090 | V | E | 1 |
| V510G | 510 | 23090 | V | G | 1 |
| V510I | 510 | 23090 | V | I | 1 |

## Notes

### Competing Interest Statement

The authors have declared no competing interest.

## REFERENCES

Lan, J., Ge, J., Yu, J., Shan, S., Zhou, H., Fan, S., Zhang, Q., Shi, X., Wang, Q., Zhang, L., & Wang, X. (2020). Structure of the SARS-CoV-2 spike receptor-binding domain bound to the ACE2 receptor. Nature, 581(7807), 215–220. https://doi.org/10.1038/s41586-020-2180-5

Wang, Q., Zhang, Y., Wu, L., Niu, S., Song, C., Zhang, Z., Lu, G., Qiao, C., Hu, Y., Yuen, K. Y., Wang, Q., Zhou, H., Yan, J., & Qi, J. (2020). Structural and Functional Basis of SARS-CoV-2 Entry by Using Human ACE2. Cell, 181(4), 894–904.e899. https://doi.org/10.1016/j.cell.2020.03.045

Leung, K., Shum, M. H., Leung, G. M., Lam, T. T., & Wu, J. T. (2021). Early transmissibility assessment of the N501Y mutant strains of SARS-CoV-2 in the United Kingdom, October to November 2020. Euro Surveillance, 26(1). https://doi.org/10.2807/1560-7917.Es.2020.26.1.2002106

Torjesen, I. (2021). Covid-19: Delta variant is now UK’s most dominant strain and spreading through schools. BMJ, 373, n1445. https://doi.org/10.1136/bmj.n1445

Karplus, M. (2002). Molecular Dynamics Simulations of Biomolecules. Accounts of Chemical Research, 35(6), 321–323. https://doi.org/10.1021/ar020082r

Sinha, S., & Wang, S. M. (2020). Classification of VUS and unclassified variants in BRCA1 BRCT repeats by molecular dynamics simulation. Computational and Structural Biotechnology Journal, 18, 723–736. https://doi.org/10.1016/j.csbj.2020.03.013

Kufareva, I., & Abagyan, R. (2012). Methods of protein structure comparison. Methods in Molecular Biology, 857, 231–257. https://doi.org/10.1007/978-1-61779-588-6_10

Gumbart, J. C., Roux, B., & Chipot, C. (2013). Efficient Determination of Protein–Protein Standard Binding Free Energies from First Principles. Journal of Chemical Theory and Computation, 9(8), 3789–3798. https://doi.org/10.1021/ct400273t

Dong, Y. W., Liao, M. L., Meng, X. L., & Somero, G. N. (2018). Structural flexibility and protein adaptation to temperature: Molecular dynamics analysis of malate dehydrogenases of marine molluscs. Proceedings of the National Academy of Sciences, 115(6), 1274–1279. https://doi.org/10.1073/pnas.1718910115

Benson, N. C., & Daggett, V. (2012). A Comparison of Multiscale Methods for the Analysis of Molecular Dynamics Simulations. The Journal of Physical Chemistry B, 116(29), 8722–8731. https://doi.org/10.1021/jp302103t

Daidone, I., Amadei, A., Roccatano, D., & Nola, A. D. (2003). Molecular Dynamics Simulation of Protein Folding by Essential Dynamics Sampling: Folding Landscape of Horse Heart Cytochrome c. Biophysical Journal, 85(5), 2865–2871. https://doi.org/10.1016/S0006-3495(03)74709-2

Zhang, D., & Lazim, R. (2017). Application of conventional molecular dynamics simulation in evaluating the stability of apomyoglobin in urea solution. Scientific Reports, 7, 44651. https://doi.org/10.1038/srep44651

Yousefpour, A., Modarress, H., Goharpey, F., & Amjad-Iranagh, S. (2015). Interaction of PEGylated anti-hypertensive drugs, amlodipine, atenolol and lisinopril with lipid bilayer membrane: A molecular dynamics simulation study. Biochim Biophys Acta, 1848(8), 1687–1698. https://doi.org/10.1016/j.bbamem.2015.04.016

Barnes, C. O., Jette, C. A., Abernathy, M. E., Dam, K. A., Esswein, S. R., Gristick, H. B., Malyutin, A. G., Sharaf, N. G., Huey-Tubman, K. E., Lee, Y. E., Robbiani, D. F., Nussenzweig, M. C., West, A. P., Jr., & Bjorkman, P. J. (2020). SARS-CoV-2 neutralizing antibody structures inform therapeutic strategies. Nature, 588(7839), 682–687. https://doi.org/10.1038/s41586-020-2852-1

Weisblum, Y., Schmidt, F., Zhang, F., DaSilva, J., Poston, D., Lorenzi, J. C., Muecksch, F., Rutkowska, M., Hoffmann, H. H., Michailidis, E., Gaebler, C., Agudelo, M., Cho, A., Wang, Z., Gazumyan, A., Cipolla, M., Luchsinger, L., Hillyer, C. D., Caskey, M., Robbiani, D. F., Rice, C. M., Nussenzweig, M. C., Hatziioannou, T., & Bieniasz, P. D. (2020). Escape from neutralizing antibodies by SARS-CoV-2 spike protein variants. Elife, 9, e61312. https://doi.org/10.7554/eLife.61312

Scudellari, M. (2021). How the coronavirus infects cells - and why Delta is so dangerous. Nature, 595(7869), 640–644. https://doi.org/10.1038/d41586-021-02039-y

Gan, H. H., Twaddle, A., Marchand, B., & Gunsalus, K. C. (2021). Structural Modeling of the SARS-CoV-2 Spike/Human ACE2 Complex Interface can Identify High-Affinity Variants Associated with Increased Transmissibility. J Mol Biol, 433(15), 167051. https://doi.org/10.1016/j.jmb.2021.167051

Ghorbani, M., Brooks, B. R., & Klauda, J. B. (2020). Critical Sequence Hotspots for Binding of Novel Coronavirus to Angiotensin Converter Enzyme as Evaluated by Molecular Simulations. The Journal of Physical Chemistry B, 124(45), 10034–10047. https://doi.org/10.1021/acs.jpcb.0c05994

Wang, Y., Liu, M., & Gao, J. (2020). Enhanced receptor binding of SARS-CoV-2 through networks of hydrogen-bonding and hydrophobic interactions. Proceedings of the National Academy of Sciences, 117(25), 13967–13974. https://doi.org/10.1073/pnas.2008209117

Yi, C., Sun, X., Ye, J., Ding, L., Liu, M., Yang, Z., Lu, X., Zhang, Y., Ma, L., Gu, W., Qu, A., Xu, J., Shi, Z., Ling, Z., & Sun, B. (2020). Key residues of the receptor binding motif in the spike protein of SARS-CoV-2 that interact with ACE2 and neutralizing antibodies. Cellular & Molecular Immunology, 17(6), 621–630. https://doi.org/10.1038/s41423-020-0458-z

Khan, A., Zia, T., Suleman, M., Khan, T., Ali, S. S., Abbasi, A. A., Mohammad, A., & Wei, D. Q. (2021). Higher infectivity of the SARS-CoV-2 new variants is associated with K417N/T, E484K, and N501Y mutants: An insight from structural data. Journal of Cellular Physiology. https://doi.org/10.1002/jcp.30367

Teruel N, Mailhot O, Najmanovich RJ (2021) Modelling conformational state dynamics and its role on infection for SARS-CoV-2 Spike protein variants. PLoS Comput Biol 17(8): e1009286. https://doi.org/10.1371/journal.pcbi.1009286

Fiorentini, S., Messali, S., Zani, A., Caccuri, F., Giovanetti, M., Ciccozzi, M., & Caruso, A. (2021). First detection of SARS-CoV-2 spike protein N501 mutation in Italy in August, 2020. The Lancet Infectious Diseases, 21(6), e147. https://doi.org/10.1016/S1473-3099(21)00007-4

Mugnai, M. L., Templeton, C., Elber, R., & Thirumalai, D. (2020). Role of Long-range Allosteric Communication in Determining the Stability and Disassembly of SARS-COV-2 in Complex with ACE2. bioRxiv. https://doi.org/10.1101/2020.11.30.405340

Luan, B., Wang, H., & Huynh, T. (2021). Molecular Mechanism of the N501Y Mutation for Enhanced Binding between SARS-CoV-2’s Spike Protein and Human ACE2 Receptor. bioRxiv, 2021.2001.2004.425316. https://doi.org/10.1101/2021.01.04.425316

Nelson, G., Buzko, O., Spilman, P., Niazi, K., Rabizadeh, S., & Soon-Shiong, P. (2021). Molecular dynamic simulation reveals E484K mutation enhances spike RBD-ACE2 affinity and the combination of E484K, K417N and N501Y mutations (501Y.V2 variant) induces conformational change greater than N501Y mutant alone, potentially resulting in an escape mutant. bioRxiv, 2021.2001.2013.426558. https://doi.org/10.1101/2021.01.13.426558

Korber, B., Fischer, W. M., Gnanakaran, S., Yoon, H., Theiler, J., Abfalterer, W., Hengartner, N., Giorgi, E. E., Bhattacharya, T., Foley, B., Hastie, K. M., Parker, M. D., Partridge, D. G., Evans, C. M., Freeman, T. M., de Silva, T. I., Angyal, A., Brown, R. L., Carrilero, L., Green, L. R., Groves, D. C., Johnson, K. J., Keeley, A. J., Lindsey, B. B., Parsons, P. J., Raza, M., Rowland-Jones, S., Smith, N., Tucker, R. M., Wang, D., Wyles, M. D., McDanal, C., Perez, L. G., Tang, H., Moon-Walker, A., Whelan, S. P., LaBranche, C. C., Saphire, E. O., & Montefiori, D. C. (2020). Tracking Changes in SARS-CoV-2 Spike: Evidence that D614G Increases Infectivity of the COVID-19 Virus. Cell, 182(4), 812–827.e819. https://doi.org/10.1016/j.cell.2020.06.043

Baum, A., Fulton, B. O., Wloga, E., Copin, R., Pascal, K. E., Russo, V., Giordano, S., Lanza, K., Negron, N., Ni, M., Wei, Y., Atwal, G. S., Murphy, A. J., Stahl, N., Yancopoulos, G. D., & Kyratsous, C. A. (2020). Antibody cocktail to SARS-CoV-2 spike protein prevents rapid mutational escape seen with individual antibodies. Science, 369(6506), 1014–1018. https://doi.org/10.1126/science.abd0831

Ju, B., Zhang, Q., Ge, J., Wang, R., Sun, J., Ge, X., Yu, J., Shan, S., Zhou, B., Song, S., Tang, X., Yu, J., Lan, J., Yuan, J., Wang, H., Zhao, J., Zhang, S., Wang, Y., Shi, X., Liu, L., Zhao, J., Wang, X., Zhang, Z., & Zhang, L. (2020). Human neutralizing antibodies elicited by SARS-CoV-2 infection. Nature, 584(7819), 115–119. https://doi.org/10.1038/s41586-020-2380-z

Starr, T. N., Czudnochowski, N., Liu, Z., Zatta, F., Park, Y. J., Addetia, A., Pinto, D., Beltramello, M., Hernandez, P., Greaney, A. J., Marzi, R., Glass, W. G., Zhang, I., Dingens, A. S., Bowen, J. E., Tortorici, M. A., Walls, A. C., Wojcechowskyj, J. A., De Marco, A., Rosen, L. E., Zhou, J., Montiel-Ruiz, M., Kaiser, H., Dillen, J., Tucker, H., Bassi, J., Silacci-Fregni, C., Housley, M. P., di Iulio, J., Lombardo, G., Agostini, M., Sprugasci, N., Culap, K., Jaconi, S., Meury, M., Dellota, E., Abdelnabi, R., Foo, S. C., Cameroni, E., Stumpf, S., Croll, T. I., Nix, J. C., Havenar-Daughton, C., Piccoli, L., Benigni, F., Neyts, J., Telenti, A., Lempp, F. A., Pizzuto, M. S., Chodera, J. D., Hebner, C. M., Virgin, H. W., Whelan, S. P. J., Veesler, D., Corti, D., Bloom, J. D., & Snell, G. (2021). SARS-CoV-2 RBD antibodies that maximize breadth and resistance to escape. Nature. https://doi.org/10.1038/s41586-021-03807-6

Rogers, T. F., Zhao, F., Huang, D., Beutler, N., Burns, A., He, W. T., Limbo, O., Smith, C., Song, G., Woehl, J., Yang, L., Abbott, R. K., Callaghan, S., Garcia, E., Hurtado, J., Parren, M., Peng, L., Ramirez, S., Ricketts, J., Ricciardi, M. J., Rawlings, S. A., Wu, N. C., Yuan, M., Smith, D. M., Nemazee, D., Teijaro, J. R., Voss, J. E., Wilson, I. A., Andrabi, R., Briney, B., Landais, E., Sok, D., Jardine, J. G., & Burton, D. R. (2020). Isolation of potent SARS-CoV-2 neutralizing antibodies and protection from disease in a small animal model. Science (New York, N.Y.), 369(6506), 956–963. https://doi.org/10.1126/science.abc7520

Bergwerk, M., Gonen, T., Lustig, Y., Amit, S., Lipsitch, M., Cohen, C., Mandelboim, M., Gal Levin, E., Rubin, C., Indenbaum, V., Tal, I., Zavitan, M., Zuckerman, N., Bar-Chaim, A., Kreiss, Y., & Regev-Yochay, G. (2021). Covid-19 Breakthrough Infections in Vaccinated Health Care Workers. New England Journal of Medicine. https://doi.org/10.1056/NEJMoa2109072

Pettersen, E. F., Goddard, T. D., Huang, C. C., Couch, G. S., Greenblatt, D. M., Meng, E. C., & Ferrin, T. E. (2004). UCSF Chimera—A visualization system for exploratory research and analysis. Journal of Computational Chemistry, 25(13), 1605–1612. https://doi.org/10.1002/jcc.20084

Berendsen, H. J. C., van der Spoel, D., & van Drunen, R. (1995). GROMACS: A message-passing parallel molecular dynamics implementation. Computer Physics Communications, 91(1), 43–56. https://doi.org/10.1016/0010-4655(95)00042-E

Hess, B. (2008). P-LINCS: A Parallel Linear Constraint Solver for Molecular Simulation. Journal of Chemical Theory and Computation, 4(1), 116–122. https://doi.org/10.1021/ct700200b

Parrinello, M., & Rahman, A. (1981). Polymorphic transitions in single crystals: A new molecular dynamics method. Journal of Applied Physics, 52(12), 7182–7190. https://doi.org/10.1063/1.328693

Turner, P. (2005). XMGRACE, Version 5.1. 19. Center for Coastal and Land-Margin Research, Oregon Graduate Institute of Science and Technology, Beaverton, OR.

Genheden, S., & Ryde, U. (2015). The MM/PBSA and MM/GBSA methods to estimate ligand-binding affinities. Expert opinion on drug discovery, 10(5), 449–461. https://doi.org/10.1517/17460441.2015.1032936

Cieplak, P., Caldwell, J., & Kollman, P. (2001). Molecular mechanical models for organic and biological systems going beyond the atom centered two body additive approximation: aqueous solution free energies of methanol and N-methyl acetamide, nucleic acid base, and amide hydrogen bonding and chloroform/water partition coefficients of the nucleic acid bases. Journal of Computational Chemistry, 22(10), 1048–1057. https://doi.org/10.1002/jcc.1065

Onufriev, A., Bashford, D., & Case, D. A. (2004). Exploring protein native states and large-scale conformational changes with a modified generalized born model. Proteins, 55(2), 383–394. https://doi.org/10.1002/prot.20033

